# ACID: A Comprehensive Toolbox for Image Processing and Modeling of Brain, Spinal Cord, and Ex Vivo Diffusion MRI Data

**DOI:** 10.1101/2023.10.13.562027

**Authors:** Gergely David, Björn Fricke, Jan Malte Oeschger, Lars Ruthotto, Francisco J. Fritz, Ora Ohana, Laurin Mordhorst, Thomas Sauvigny, Patrick Freund, Karsten Tabelow, Siawoosh Mohammadi

## Abstract

Diffusion MRI (dMRI) has become a crucial imaging technique in the field of neuroscience, with a growing number of clinical applications. Although most studies still focus on the brain, there is a growing interest in utilizing dMRI to investigate the healthy or injured spinal cord. The past decade has also seen the development of biophysical models that link MR-based diffusion measures to underlying microscopic tissue characteristics, which necessitates validation through ex vivo dMRI measurements. Building upon 13 years of research and development, we present an open-source, MATLAB-based academic software toolkit dubbed ACID: **A** Comprehensive Toolbox for Image Processing and Modeling of Brain, Spinal Cord, and Ex Vivo Diffusion MRI Data. ACID is an extension to the Statistical Parametric Mapping (SPM) software, designed to process and model dMRI data of the brain, spinal cord, and ex vivo specimens by incorporating state-of-the-art artifact correction tools, diffusion and kurtosis tensor imaging, and biophysical models that enable the estimation of microstructural properties in white matter. Additionally, the software includes an array of linear and non-linear fitting algorithms for accurate diffusion parameter estimation. By adhering to the Brain Imaging Data Structure (BIDS) data organization principles, ACID facilitates standardized analysis, ensures compatibility with other BIDS-compliant software, and aligns with the growing availability of large databases utilizing the BIDS format. Furthermore, being integrated into the popular SPM framework, ACID benefits from a wide range of segmentation, spatial processing, and statistical analysis tools as well as a large and growing number of SPM extensions. As such, this comprehensive toolbox covers the entire processing chain from raw DICOM data to group-level statistics, all within a single software package.

## 1. Introduction

Diffusion MRI (dMRI) exploits the self-diffusion of water molecules to produce images that are sensitive to tissue microstructure by measuring the diffusion along various spatial directions (Callaghan et al., 1988; Le Bihan et al., 1988; Stejskal & Tanner, 1965). dMRI has been applied to study a number of phenomena including normal brain development (Dubois et al2014; Miller et al 2002), aging (Draganski et al., 2011; Sullivan et al., 2010), training-induced plasticity (Scholz et al., 2009), and monitoring progression of and recovery from neurological diseases (Farbota et al., 2012; Meinzer et al., 2010). Clinical applications of dMRI include the diagnosis of ischemic stroke (Urbach et al., 2000), multiple sclerosis (Horsfield et al., 1996), cancer and metastases (Gerstner and Sorensen, 2011), and surgical planning of brain tumors (Chun et al., 2005). Although the vast majority of dMRI applications has focused on the brain, there is a growing interest in spinal cord dMRI, as researchers seek sensitive and predictive markers of spinal cord white matter damage (Cohen et al., 2017; Martin et al., 2016). Furthermore, an increasing number of studies utilize dMRI on ex vivo specimens for comparative analysis with other imaging modalities, such as electron microscopy (Barazany et al., 2009; Keim et al., 2016; Papazoglou et al., 2023).

To fully utilize the sensitivity of dMRI to tissue microstructure, expert knowledge is required to minimize artifacts both during acquisition, e.g., by cardiac gating or twice-refocused spin-echo sequences, and through dedicated retrospective correction methods. Commonly used retrospective correction techniques include motion and eddy current correction (J. L. R. Andersson & Sotiropoulos, 2016; Mohammadi et al., 2010), susceptibility distortion correction (Gu & Eklund, 2019; Ruthotto et al., 2012), denoising (Becker et al., 2014; Veraart et al., 2016), Rician bias correction (Oeschger et al., 2023a; Sijbers et al., 1998), and robust tensor fitting techniques (Chang et al., 2005; Mohammadi et al., 2013). Retrospective artifact correction techniques, along with diffusion signal modeling capabilities, are widely available in open-source toolboxes such as FSL-FDT (Smith et al., 2004), DiPY (Garyfallidis et al., 2014), DESIGNER (Ades-Aron et al., 2018), ExploreDTI (Leemans et al., 2009), MRtrix3 (Tournier et al., 2019), TORTOISE (Pierpaoli et al., 2010), AFNI-FATCAT (Taylor & Saad, 2013), and others.

While the majority of toolboxes have been designed for brain dMRI, ACID has introduced several features and utilities that make it particularly suitable for spinal cord and ex vivo dMRI as well. Specifically, ACID addresses the higher level and different nature of artifacts in spinal cord dMRI (Barker, 2001; Stroman et al., 2014), and the highly variable geometry and diffusion properties in ex vivo dMRI (see Sébille et al., 2019 for a list of ex vivo/post-mortem dMRI studies). Although there are some software options available for processing spinal cord images, most notably the Spinal Cord Toolbox (De Leener et al., 2017), these tools lack comprehensive artifact correction and biophysical modeling capabilities for estimation of dMRI-based metrics related to microscopic tissue properties.

Biophysical modeling estimates microstructural properties, such as axonal water fraction and orientation dispersion, as aggregated measures on the voxel level, providing greater specificity than standard diffusion tensor (DTI) or diffusion kurtosis imaging (DKI). Toolboxes dedicated for biophysical modelling of the dMRI signal, such as the NODDI (Zhang et al., 2012) or SMI toolbox (Coelho et al., 2022), typically do not include a comprehensive processing pipeline to correct for artifacts in dMRI data. In addition, to date, only a few of the dMRI toolboxes support the Brain Imaging Data Structure (BIDS, Gorgolewski et al., 2016) standard for organizing and annotating raw and processed dMRI data. The lack of standardization complicates not only the sharing and aggregation of processed dMRI data but also the application of automated image analysis tools designed for big data, such as machine learning techniques. Over the past two decades, tens of thousands of dMRI datasets have been made openly available in large neuroimaging databases (e.g., HCP (Van Essen et al., 2013) and the UK Biobank (Littlejohns et al., 2020)), underscoring the importance of consistent data storage practices.

Building upon 13 years of research and development, we introduce an open-source MATLAB-based extension to the Statistical Parametric Mapping (SPM) software, the ACID toolbox: A Comprehensive Toolbox for Image Processing and Modeling of Brain, Spinal Cord, and Ex Vivo Diffusion MRI Data. ACID was initially developed as a collection of artifact correction tools but has now been extended to a comprehensive toolbox for processing and modeling of dMRI data. In particular, ACID offers (i) state-of-the-art image processing tools as well as (ii) DTI, DKI, and white matter biophysical model parameter estimation methods optimized for brain, spinal cord, and ex vivo dMRI data. Additionally, (iii) ACID adheres to the BIDS standard for organizing the output, making the processed images compliant with other BIDS software and facilitating data sharing. Finally, (iv) ACID is embedded in the SPM framework to benefit from its established functions including spatial processing tools and statistical inference schemes. ACID tools can be combined with other SPM functions to create pipelines in SPM batch system, which offers an all-in-one software solution from conversion of DICOM data to statistical group analysis. ACID also benefits from a large and growing number of SPM extensions. For example, ACID can be combined with the SPM12-based hMRI toolbox (Tabelow et al., 2019) to perform multi-contrast analysis of dMRI and other quantitative MRI data, such as relaxation rates, acquired from the same subject, all within a single pipeline. Many of the methods used in the ACID toolbox have already been published in the scientific dMRI literature (Table 1). In this paper, we detail the design and function of the ACID modules and provide guidance on their optimal combination for brain, spinal cord, and ex vivo applications.

**Table 1.**
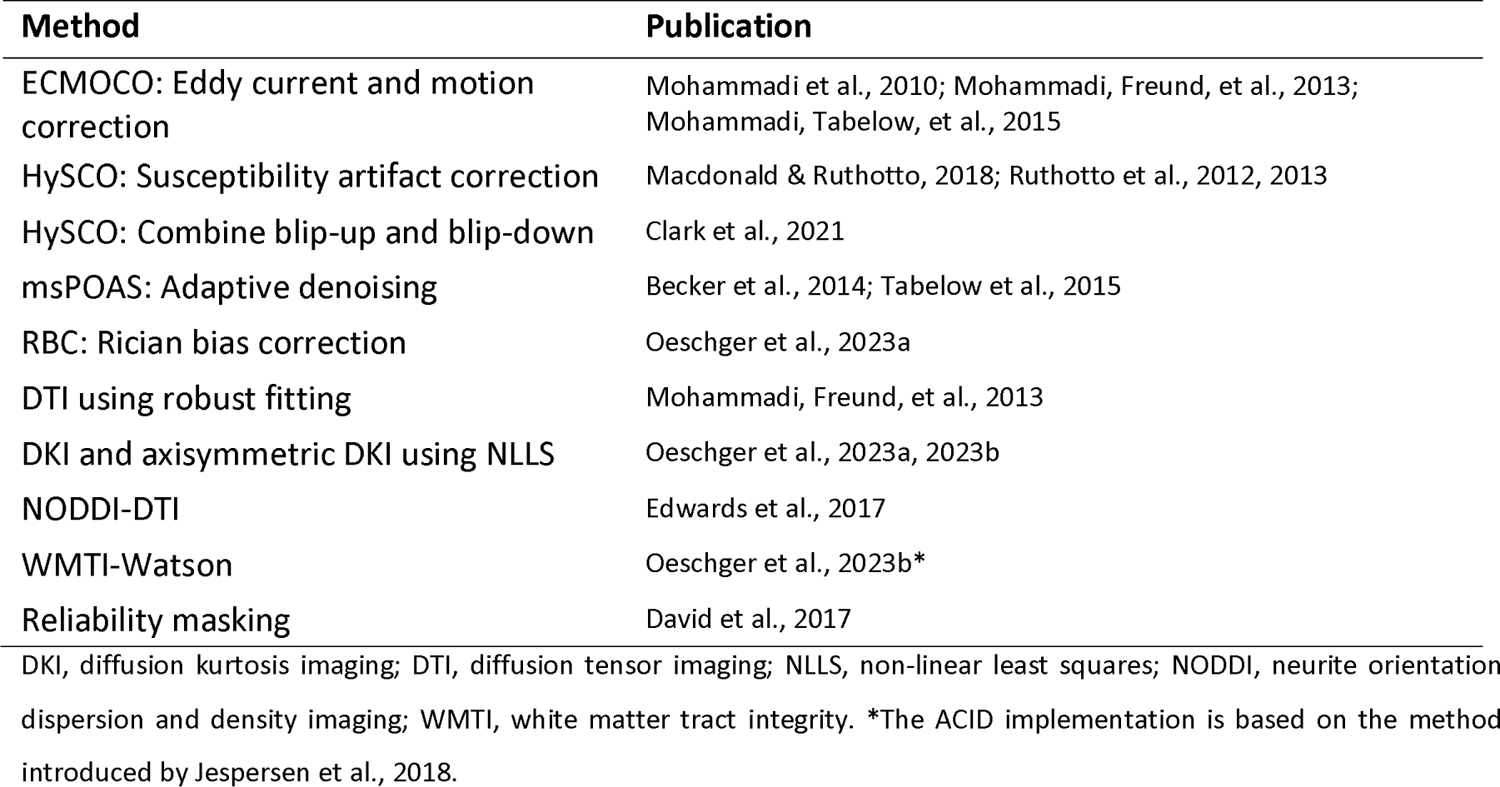
Peer-reviewed methods used in the ACID toolbox.

## 2. Methods

### 2.1 Overview

The ACID toolbox is a comprehensive toolbox for processing and analyzing dMRI data, built upon the following four pillars: (1) pre-processing of dMRI data *(Pre-processing* module), (2) physical models of the diffusion signal *(Diffusion tensor/kurtosis imaging* module), (3) white matter biophysical models of the diffusion signal *(Biophysical models* module), and (4) additional features referred to as *Utilities.* The *Pre-processing* module consists of state-of-the-art methods for retrospective correction of the dMRI data. The *Diffusion tensor/kurtosis imaging* module contains tensor and kurtosis models that can be applied to dMRI data from various tissues or samples, including gray and white matter, as well as diffusion phantoms (Woletz et al., 2024). In contrast, the *Biophysical models* module can only be applied to samples that fall within their validity ranges (see Section 4.2.2). The *Utilities* module contains various useful tools, including masking and noise estimation. The ACID toolbox follows the BIDS convention and enables the seamless integration of external tools into its processing pipeline in a modular fashion *(External tools* module). More details about the implementation and organization of ACID are provided in Appendix A.

### 2.2 Pre-processing

In this chapter, we provide brief descriptions of each artifact correction tool currently implemented in ACID. For detailed recommendations on various dMRI datasets (in vivo brain, in vivo spinal cord, ex vivo/post-mortem), refer to Sections 3.2 and 4.1, as well as Table 5.

#### 2.2.1 Eddy current and motion correction (ECMOCO)

ACID uses the eddy current and motion correction (ECMOCO) algorithm (Mohammadi et al., 2010) to correct for spatial misalignments that may occur between dMRI volumes. These misalignments can be caused by motion and eddy currents induced by the rapidly varying field of the diffusion-sensitizing gradients (Jezzard et al., 1998), which may lead to biased diffusion estimates (Mohammadi et al., 2013). ECMOCO aligns all source volumes to a target volume using a co­registration algorithm with an affine transformation (Friston & Ashburner, 1997) implemented in the SPM function *spm_coreg.* It was previously shown that the robustness of registration can be increased by separately registering diffusion-weighted (DW) and non-diffusion-weighted (bO) volumes to their corresponding target volumes (Mohammadi et al., 2015a). ECMOCO features the multi-target registration mode, where source volumes from each diffusion shell (b-value) are co­registered to their shell-specific target volume (Fig. B1). ECMOCO rotates the b-vectors by the obtained rotational parameters; the rotated b-vectors can be passed on to subsequent processing steps. Of note, the affine transformation of ECMOCO can only correct for first-order eddy-current displacements. The advantages and disadvantages of ECMOCO compared to other established tools, such as FSL eddy, are discussed in Section 4.1.

In spinal cord dMRI, eddy current and motion correction is more challenging than in brain dMRI due to the considerably lower number of voxels and lower signal-to-noise ratio (SNR), particularly in volumes with high b-values (>1000 s/mm^2^) or with diffusion-sensitizing gradients parallel to the spinal cord. While movement of the brain can be considered approximately rigid, the spinal cord may experience varying degrees of displacement along the rostro-caudal axis caused by factors such as breathing, pulsation of the cerebrospinal fluid, or swallowing (Yiannakas et al., 2012). To address this, we introduced *slice-wise* (2D) registration, which independently aligns each slice of the source volume to the corresponding slice of the target volume, thereby correcting for non-rigid, slice-dependent displacements (Mohammadi, Freund, et al., 2013). For more details on ECMOCO, including other recently introduced features *(initialized registration* and *exclusion mode),* refer to Appendix B.

#### 2.2.2 Adaptive denoising (msPOAS)

The Multi-shell Position-Orientation Adaptive Smoothing (msPOAS) is an iterative adaptive denoising algorithm designed to adaptively reduce noise-induced variance in dMRI data while preserving tissue boundaries, as illustrated in Fig. 3 (Becker et al., 2012, 2014; Tabelow et al., 2015). The algorithm adapts to the intensity values and their distance in both voxel space and the spherical space of diffusion directions, allowing smoothing only within spatially homogeneous areas of the DW images. One of the key advantages of msPOAS is its compatibility with all diffusion models as it operates on the raw dMRI data. Adjustable parameters include *kstar* (number of iterations that define the image smoothness), *lambda* (adaptation parameter that defines the strength of edge detection), *kappa* (initial ratio of the amount of smoothing between the local space of neighboring voxels and the spherical space of diffusion gradients), *ncoils* (i.e., the effective number of receiver coils that contributed to the measured signal). To distinguish random fluctuations from structural differences, msPOAS requires an estimate of SNR, or equivalently the noise standard deviation *(sigma).* A higher *kstar* leads to greater smoothness within homogeneous image regions, while a larger *lambda* results in weaker adaptation and more blurring at tissue edges. The optimal *kappa* depends on the number of directions per shell, while *ncoils* should be the same as the value used for noise estimation. When using msPOAS, we recommend starting with the default parameters and the *sigma* estimated with the *Noise estimation* utility function (Table 2). In case of insufficient noise reduction, parameters should be adjusted according to Appendix D.

**Table 2.** List of the ACID utility functions.

| FUNCTION | DESCRIPTION |
| --- | --- |
| **Cropping** | Crops images to a smaller size for less storage space and faster processing.<br>*Input:* image(s) to crop, new matrix size, and voxel coordinates of the center of cropping. The center of cropping can also be selected manually through a pop-up window.<br>*Output:* cropped image(s) and the cropping parameters.<br>*Application:* typically in spinal cord dMRI, where the spinal cord occupies a small portion of the image. |
| **Resampling** | Resamples images to the desired resolution.<br>*Input:* image(s) to be resampled, desired resolution, and type of interpolation (as defined in *spm_slice_vol*). Available types of interpolation: nearest neighbor, trilinear, higher-order Lagrange polynomial (2 to 127), and different orders of sinc interpolation (-1 to -127); default: -7, i.e., 7<sup>th</sup>-order sinc interpolation.<br>*Output:* resampled image(s).<br>*Application:* for example, when performing voxel-wise arithmetic between two or more images with different resolutions (e.g., g-ratio mapping). |
| **Slice-wise realignment** | Enables manual translation and scaling of images along the x and y dimensions on a slice-by-slice basis, facilitated by intensity contour lines of the source image superimposed on the target image.<br>*Input:* image to be realigned, target image, and other images to which the realignment parameters are applied.<br>*Output:* realigned image(s) and the realignment parameters.<br>*Application:* useful for realigning spinal cord images, where residual misalignments are often slice-dependent. |
| **Fusion** | Merges two images with different field of views (FOV), such as a brain and a spinal cord image, into a single combined image (Fig. 5).<br>*Inputs:* two images to be merged and a target image (typically a structural image with a larger FOV).<br>*Output:* a combined image, resampled onto to the target image. The voxel intensity values in overlapping regions are the average of the intensity values in both images. Note that before merging the images, they must be in the correct spatial position; if necessary, image realignment can be performed using the SPM *Realign* or the *Slice-wise realignment* utility function.<br>*Application:* useful for merging a brain and a spinal cord image into a single image |
|  | before applying a warping field obtained from a large-FOV structural image. |
| **Create brain mask** | Creates a binary brain mask by (i) segmenting the brain image into gray matter, white matter, and cerebrospinal fluid using SPM12's unified segmentation tool (Ashburner & Friston, 2005), (ii) summing up the resulting probability maps, and (iii) thresholding it at a certain value (accessible through the script `acid_local_defaults.m`; default: 0.8).<br>*Input:* a single brain image or tissue probability maps for gray matter, white matter, and cerebrospinal fluid, and optionally a dMRI dataset to be masked.<br>*Output:* binary brain mask and optionally a masked dMRI dataset.<br>*Application:* to restrict the estimation of DTI, DKI, and biophysical parameters to the brain for increased speed and efficiency. |
| **Reliability masking** | Aims to identify "unreliable" voxels, i.e., voxels irreversibly corrupted by artifacts. Reliability masks are generated by thresholding the root-mean-square model-fit error ($\text{rms}(\epsilon)$) map (David et al., 2017).<br>*Input:* $\text{rms}(\epsilon)$ maps (output by tensor fitting methods with label: RMS-ERROR) and the desired threshold value. The optimal threshold can be determined using the *Determine threshold* submodule.<br>*Output:* a binary reliability mask.<br>*Application:* Reliability masks can serve as binary masks in region-of-interest-based analyses. In principle, reliability masking as an outlier rejection technique is applicable after each model fitting method. It is particularly useful in situations where many data points are affected by outliers (often the case in spinal cord dMRI), which could otherwise lead to unstable tensor fits and inaccurate tensor estimates (see David et al., 2017 for examples). |
| **DWI series browser** | Enables browsing through the slices of the dMRI data for quality assessment. Slices with low SNR and/or artifacts can be identified and labeled.<br>*Input:* the dMRI dataset, b-values, and b-vectors.<br>*Output:* list of labeled slices.<br>*Application:* The saved labels can be used to inform ECMOCO about unreliable slices (see *Exclusion mode* in Appendix B). |
| **DWI series movie** | Enables simultaneous streaming of images from multiple dMRI datasets in video mode for quality assessment.<br>*Input:* a reference image and up to three dMRI datasets.<br>*Output:* a video file containing the image streams.<br>*Application:* useful for visual assessment of a single dMRI dataset or for comparing images before and after a specific processing step (e.g., ECMOCO). |
| **Noise estimation** | Estimates the noise standard deviation ($\sigma$) in the dMRI data using either the *standard* or the *repeated measures method*. The *standard method* uses the formula $\sigma \approx \sqrt{\sum_{i \in \text{mask}} S_i^2 / (2Ln)}$, where $S_i$ is the voxel intensity within a background mask defined outside the body, $L$ is the number of voxels within the background mask, and $n$ is the effective number of coil elements that contributed to the measured signal (Constantinides et al., 1997). The *repeated measures method* uses the formula: $\sigma \approx \text{mean}_{i \text{ in ROI}} \left( \text{std}_k(S(i, k)) \right)$, where $S(i, k)$ is the voxel intensity at voxel $i$ in the $k$th repeated image (Dietrich et al., 2007). The standard deviation and mean operators are performed across the repetitions and voxels, respectively. The set of repeated images can be either the non-diffusion-weighted ($b \approx 0$) or strongly diffusion-weighted (the highest b-value) images (see Appendix C for recommendations).<br>*Input:* the raw (unprocessed) dMRI dataset, a mask (*standard method*: background mask; *repeated measures method*: see Appendix C), $n$ (for the *standard method* |
|  | only), and b-values (for the *repeated measures method* only).<br>*Output*: a single $\sigma$ (assuming a homogeneous variance).<br>*Application*: $\sigma$ serves as input for msPOAS, Rician bias correction, and diffusion tensor imaging (for fitting methods WLS and robust fitting). |
| **Rician bias simulation** | Simulates diffusion-weighted MRI signals at specified SNR values in voxels within the brain white and gray matter. The simulated signals are corrected using the specified Rician bias correction (RBC) methods (for details, see Oeschger et al., 2023a).<br>*Input*: a voxel from a list of 27 pre-defined voxels, each with different diffusion and kurtosis tensor metrics<sup>1</sup> (for details, see Oeschger et al., 2023a), a list of SNR values, and the number of noise samples.<br>*Output*: a figure showing the distance between the estimated metric and the ground truth value for each RBC method.<br>*Application*: useful for computing the required SNR for DTI, DKI, and biophysical parameter estimation. |
| **ROI analysis** | Calculates the mean value within a specified region of interest (ROI).<br>*Input*: list of images and various types of ROIs including (i) global ROIs, applied to all images in the list, (ii) subject-specific ROIs, applied only to the corresponding image, and (iii) subject-specific reliability masks, again applied only to the corresponding image (see *Reliability masking*).<br>*Output*: an array containing the mean values within the specified ROIs per subject, ROI, and (optionally) slice. When multiple types of ROIs are specified, their intersection is applied.<br>*Application*: the function offers flexibility for a range of ROI-based analyses; for example, ROI-based analysis in the native space requires a set of subject-specific ROIs, while a single global mask is sufficient in the template space (with optional reliability masks in both cases). An example application including reliability masks can be found in David et al., 2017. |

**Table 3.**
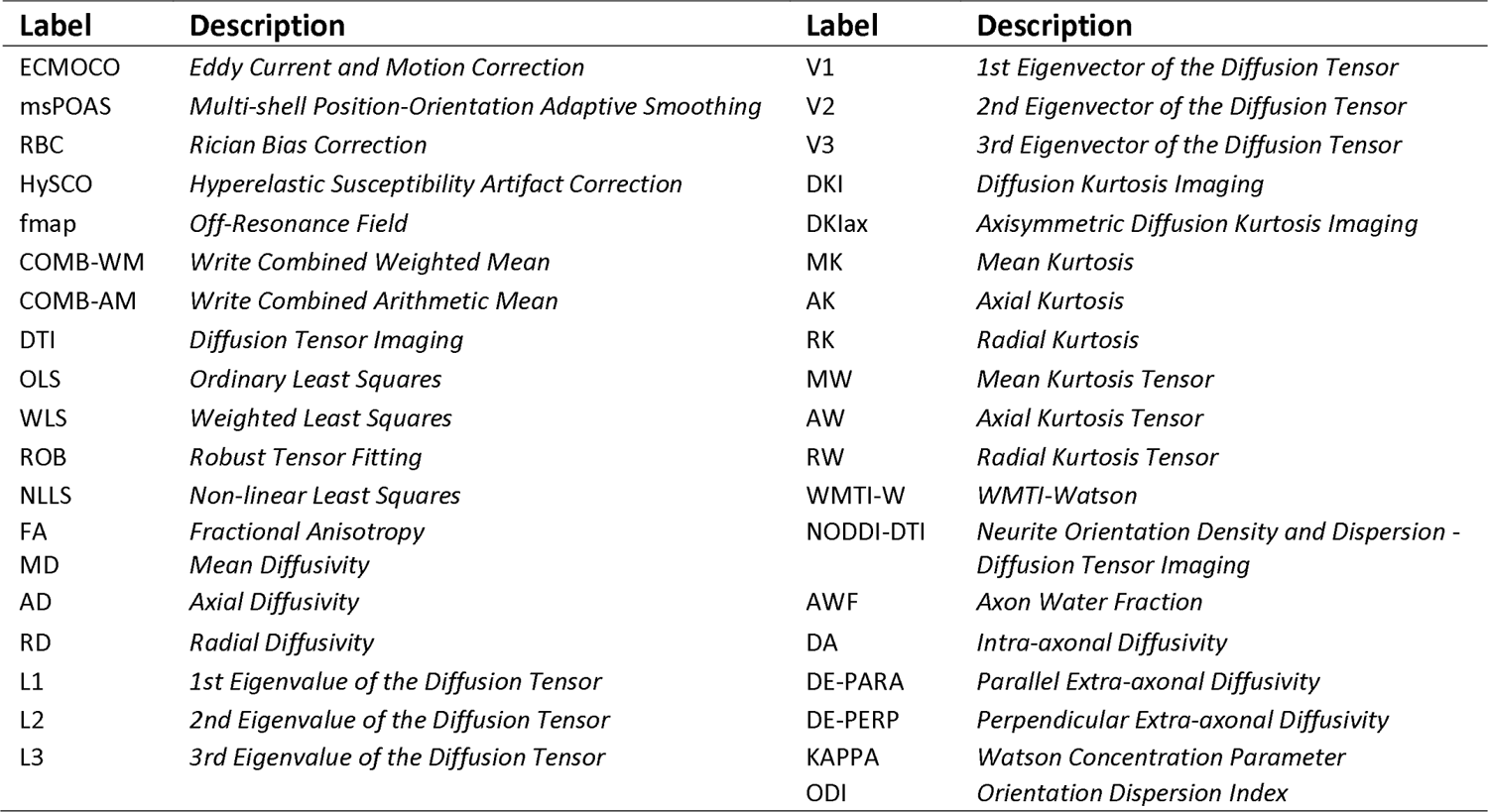
List of labels in the output filename’s desc field (not comprehensive).

#### 2.2.3 Rician bias correction

The voxel intensities of MRI magnitude images exhibit a Rician distribution in case of a single receiver coil (Gudbjartsson & Patz, 1995) and a non-central χ distribution in case of multiple receiver coils (Aja-Fernandez et al., 2014). When fitting diffusion signal models (Section 2.3), this distribution leads to a bias, known as the Rician bias, in the estimated tensor (Basser & Pajevic, 2000; Gudbjartsson & Patz, 1995; Jones & Basser, 2004) and kurtosis parameters (Veraart et al., 2011; Veraart et al., 2013a), as well as in biophysical parameter estimates (M. Andersson et al., 2022; Fan et al., 2020; Howard et al., 2022). This Rician bias is particularly relevant in low SNR situations (Polzehl & Tabelow, 2016). Two approaches of Rician bias correction (RBC) are implemented in ACID. The M2 approach, introduced in Miller & Joseph, 1993 and later extended to multi-channel receiver coil (André et al., 2014), operates on the dMRI data and uses the second moment of the non-central χ distribution of the measured intensities and noise estimates to estimate the true voxel intensities. The second approach modifies the parameter estimation by considering the non-central *χ* distribution to account for the Rician bias during model fitting (Oeschger et al., 2023a). Note that the latter approach assumes uncorrected data, therefore it must not be combined with the first method and is currently only available for non-linear least squares fitting. Both methods require an estimate of the noise standard deviation, which can be obtained using either the *standard* or the *repeated measures* method within the *Noise estimation* utility function (Table 2). Details on noise estimation are available in Appendix C. In addition, ACID offers the *Rician bias simulation* utility function to determine the optimal RBC method for the dMRI dataset and SNR at hand (Table 2). An example of how RBC influences the estimation of biophysical parameters is illustrated in Fig. F1.

#### 2.2.4 Susceptibility artifact correction (HySCO)

Hyperelastic Susceptibility Artifact Correction (HySCO) is a technique used to correct for geometric distortions caused by susceptibility artifacts (Ruthotto et al., 2012, 2013). These artifacts can occur at interfaces between tissues with different magnetic susceptibilities, such as those found near paranasal sinuses, temporal bone, and vertebral bodies. To correct for these artifacts, HySCO estimates the bias field based on a reversed-gradient spin-echo echo planar imaging (EPI) acquisition scheme. This requires the acquisition of at least one image with identical acquisition parameters as the dMRI data but with reversed phase-encoding direction, also referred to as “blip-up” or “blip-down” acquisitions. The bias field map, estimated from the blip-up and blip-down images, is applied to the entire dMRI data to unwarp the geometric distortions (see Fig. 3 for examples). For datasets that include full blip-reversed acquisition, i.e., each image was acquired with two phase-encoding directions (blip-up and blip-down), the reverse phase-encoded images can be combined using the submodule *HySCO: combine blip-up and blip-down images*.

## Diffusion signal models

The dependence of dMRI signal on the direction and strength of diffusion-weighting is commonly described by mathematical models. Two of the most widely used models are DTI (Basser et al., 1994) and DKI (Hansen et al., 2016; Jensen et al., 2005).

### 2.3.1 Diffusion tensor imaging (DTI)

DTI describes the anisotropic water diffusion in the white matter by a diffusion tensor with six independent diffusion parameters. The eigenvalues of the tensor can be used to compute rotationally invariant DTI scalar metrics including fractional anisotropy (FA) and mean (MD), axial (AD), and radial diffusivities (RD). The interpretation of DTI assumes that the direction of axial diffusivity is aligned with the white matter tracts, which may not be the case in complex fiber geometry such as crossing or fanning fibers.

ACID provides four algorithms to obtain the diffusion tensor (see Appendix E for details). Ordinary least squares (OLS) fits the tensor model by minimizing the sum of squared model-fit errors, while weighted least squares (WLS) minimizes the *weighted* sum of squared model-fit errors, accounting for the distortion of noise distribution in the linearized (logarithmic) data. Robust fitting is similar to WLS but factorizes the weights into three components to account for local and slice-specific artifacts as well, while also featuring Tikhonov regularization to handle ill-conditioned weighting matrices resulting from a high occurrence of outliers. Robust fitting is designed to downweight outliers in the model fit, which can otherwise introduce a bias in the fitted model parameters (Mohammadi et al., 2013) (Fig. E1). Unlike the linearized models, the non-linear least squared (NLLS) method is based on an implementation (Modersitzki, 2009) of the Gauss-Newton algorithm and operates on the non-logarithmic data, avoiding the distortion of noise distribution.

### 2.3.2 Diffusion kurtosis imaging (DKI)

DKI expands the diffusion tensor model by the kurtosis tensor, a fourth-order tensor with 15 independent parameters, which captures the effects of non-Gaussian water diffusion. From the 15 kurtosis parameters, several kurtosis metrics can be estimated including the mean (MK), axial (AK), and radial kurtosis (RK), as well as the mean (MW), axial (AW), and radial (RW) kurtosis tensor (Tabesh et al., 2011) (Fig. 1). These metrics provide additional information about tissue complexity beyond what can be captured by diffusion tensor metrics alone. DKI requires the acquisition of a second diffusion shell with higher b-value (typically between 2000 and 2500 s/mm^2^). ACID also includes the axisymmetric DKI model, a recent modification of DKI which reduces the parameter space to 8 independent parameters by imposing the assumption of axisymmetrically distributed axons (Hansen et al., 2016). Currently, ACID offers the OLS and NLLS algorithms for fitting the kurtosis tensor, and the NLLS algorithm for fitting the axisymmetric kurtosis tensor. Note that the diffusion tensor parameters from DKI might differ from standard DTI parameters. In particular, diffusivities (AD, MD, and RD) derived from the DTI model are often underestimated compared to those derived from the DKI model (referred to as kurtosis bias) (Edwards et al., 2017). By incorporating higher-order moments of the diffusion signal, DKI can address kurtosis bias, resulting in more accurate diffusivity estimates (see Fig. S3 in the Supplementary material for a comparison of MD derived from DTI and DKI).

**Fig. 1.**
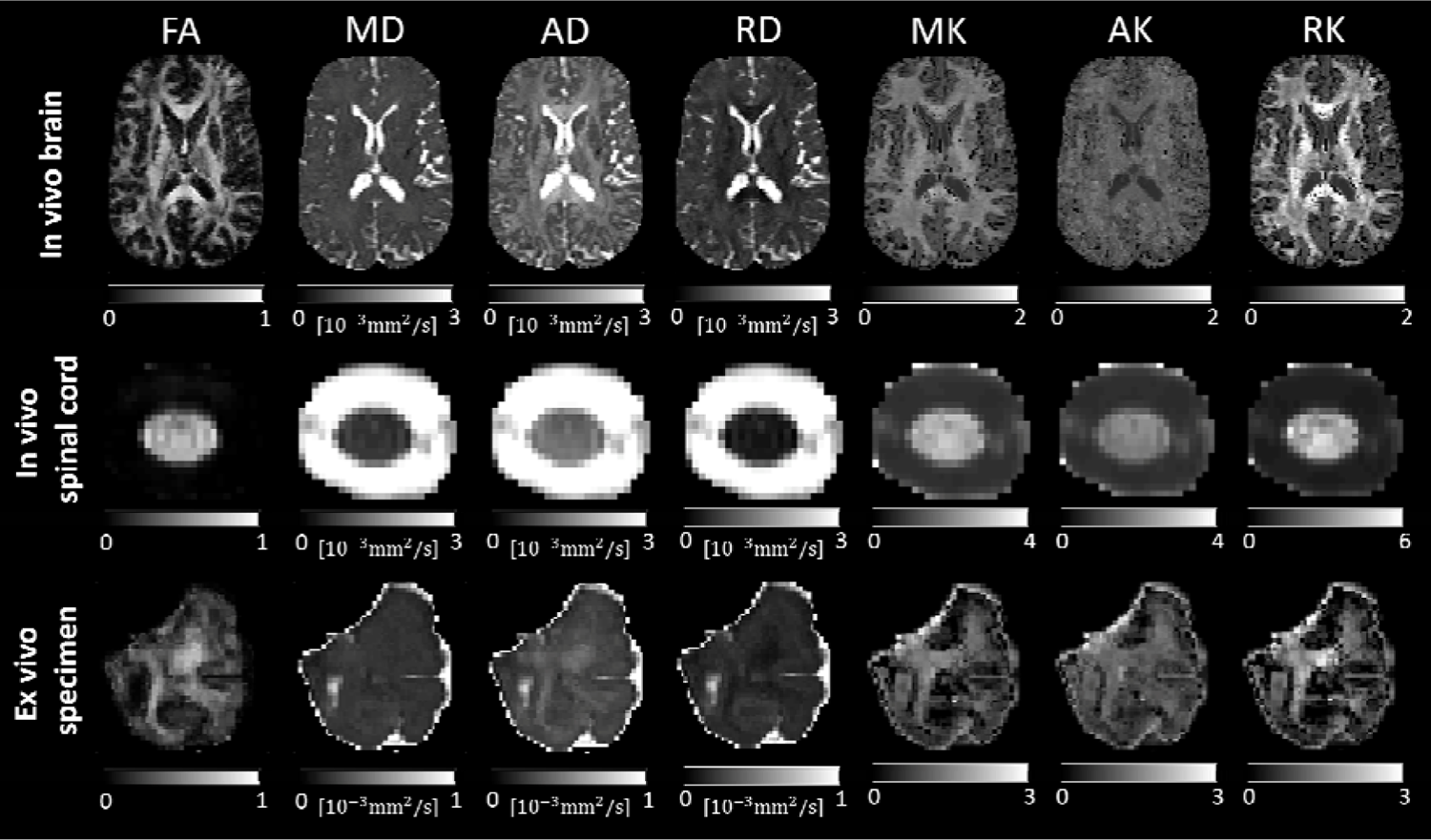
Selected maps derived from diffusion kurtosis imaging (DKI) using an in vivo brain, in vivo spinal cord, and ex vivo dMRI dataset (refer to Table 4 for details on the dataset). Shown are maps of fractional anisotropy (FA), mean diffusivity (MD), axial diffusivity (AD), radial diffusivity (RD), mean kurtosis (MK), axial kurtosis (AK), and radial kurtosis (RK).

#### 2.4 Biophysical models

Biophysical models separate the dMRI signal into distinct signal components from various tissue compartments, each with their own underlying assumptions. Biophysical models provide more specific and biologically interpretable metrics that are linked to tissue microstructure (Jelescu et al., 2020). The application of biophysical models is often referred to as dMRI-based in vivo histology (Mohammadi & Callaghan, 2021; Weiskopf et al., 2021) or microstructural dMRI (Jelescu et al., 2020; Novikov, 2021; Novikov et al., 2019). In the following, we briefly describe the two white matter biophysical models currently implemented in ACID (WMTI-Watson and NODDI-DTI), while recommendations on their usage are provided in Section 4.2.2. Example maps are shown in Fig. 2, and specific values obtained from the brain and spinal cord are presented and discussed in Fig. S5 (Supplementary material).

**Fig. 2.**
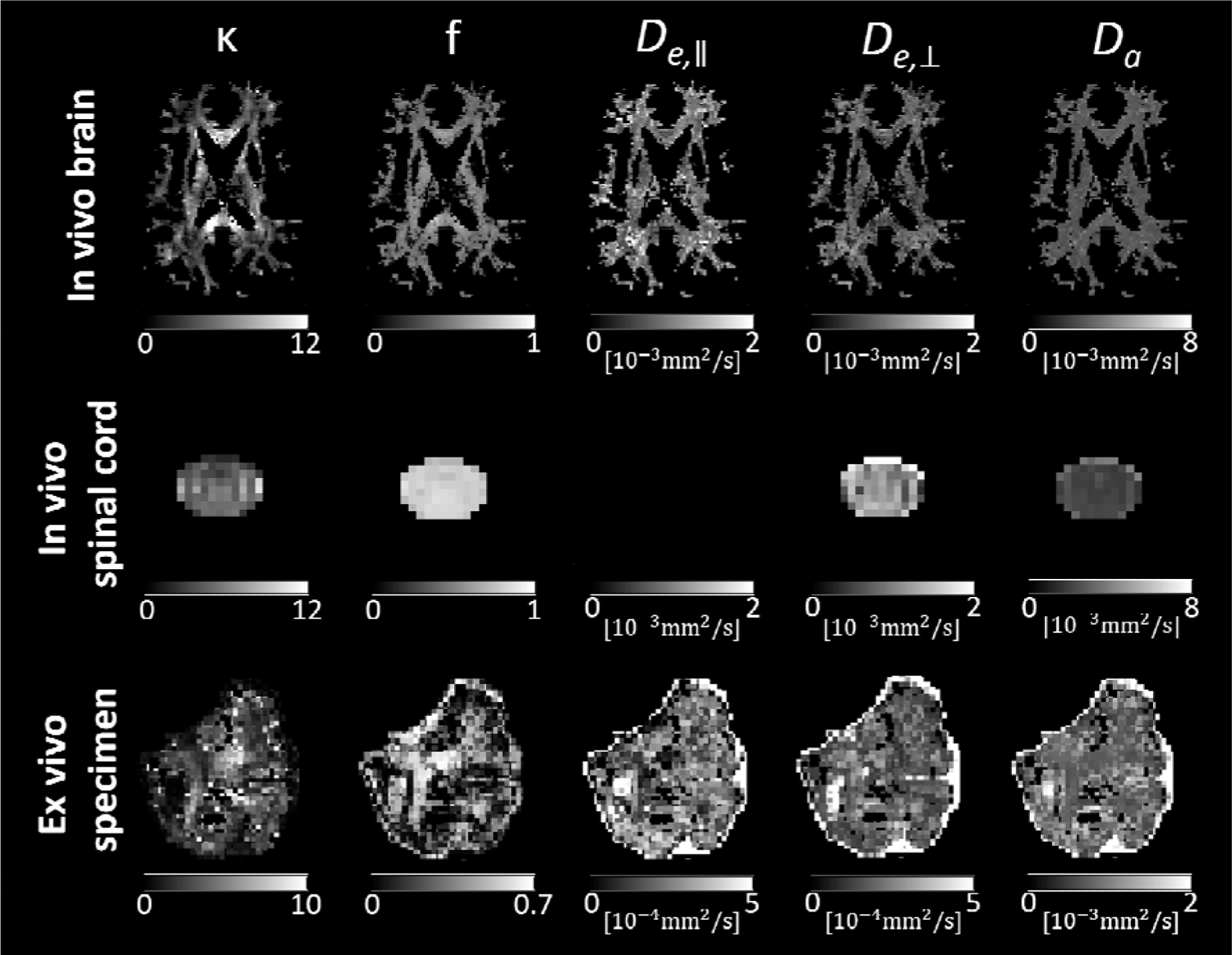
Maps of biophysical parameters derived from the WMTI-Watson model using an in vivo brain, in vivo spinal cord and ex vivo dMRI dataset (refer to Table 4 for details on the dataset). Shown are maps of Watson concentration parameter (), axonal water fraction (), parallel and perpendicular extra-axonal diffusivities (and), and intra-axonal diffusivity (). Note that for the in vivo spinal cord dataset, the maximum b-value (b=1500 s/mm^2^) was probably too low for an accurate estimation of, resulting in voxels with negative (hence unphysical) values within the spinal cord. Since WMTI-Watson is a white matter biophysical model, the parameter maps were masked for the white matter in the brain dataset. For the spinal cord and ex vivo specimen, we refrained from masking for the white matter due to the difficulty of obtaining an accurate white matter mask.

**Fig. 3.**
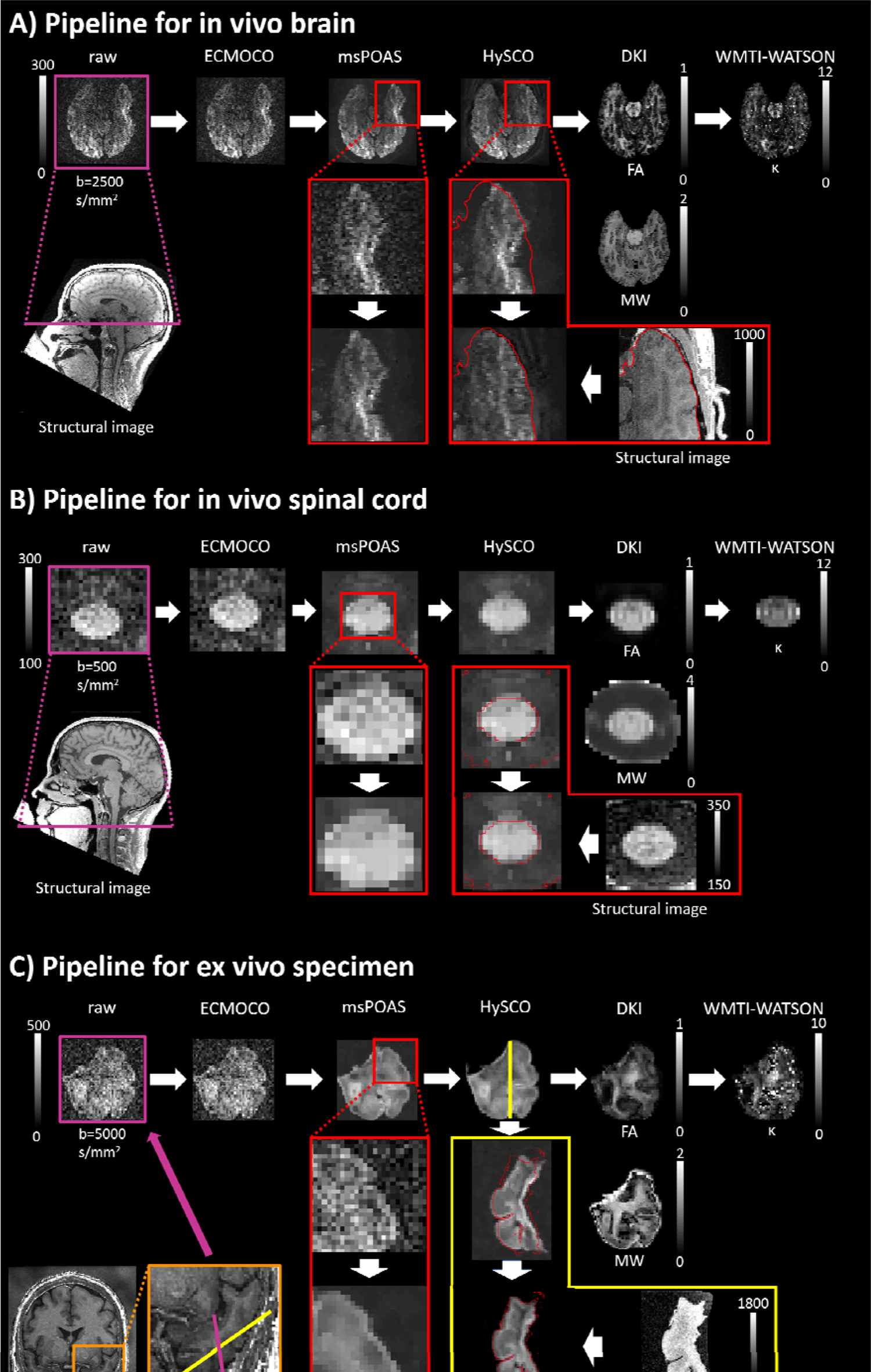
Standard processing pipelines for typical (A) in vivo brain, (B) in vivo spinal cord, and (C) ex vivo dMRI datasets (refer to Table 4 for details on the datasets and Table 5 for details on the pipeline settings). Example batches for each types of dMRI data are stored in the Example_Batches folder of the toolbox. The positions of the displayed slices of the dMRI data are indicated in purple on the corresponding structural images. For the ex vivo specimen (C), the brain region from which the sample was extracted is highlighted in an orange box. Although not explicitly shown here, noise estimation should be performed on the unprocessed data (see Appendix C), which serves as input for msPOAS, Rician bias correction, and diffusion tensor fitting (for fitting methods WLS and robust fitting). However, in case of substantial misalignments across volumes, and when using the *repeated measures* noise estimation method, it might be beneficial to perform this step after ECMOCO to prevent an overestimation of noise. For msPOAS, a zoomed-in visual comparison is shown between a diffusion-weighted (DW) image before (middle row) and after applying msPOAS (bottom row); the msPOAS-corrected image appears less noisy while preserving tissue edges. For HySCO, contour lines of the corresponding structural image (displayed as red lines) are overlaid on a zoomed-in DW image both before (middle row) and after applying HySCO (bottom row). HySCO improves the alignment between the DW and the structural image. For the in vivo brain dMRI dataset (A), an inferior slice is shown that presents high susceptibility-related distortions, making the effect of HySCO more visible. For the ex vivo dMRI dataset (C), the effect of HySCO is shown in a slice (illustrated in yellow) orthogonal to the original one (illustrated in purple) to better visualize susceptibility-related distortions and their correction. Note that HySCO is applied as the final pre-processing step, i.e., after applying msPOAS; however, the HySCO field map used for “unwrapping” the diffusion-weighted images is estimated on the ECMOCO-corrected datasets, i.e., before applying msPOAS. Rician bias correction (not explicitly shown here) should be applied either before (recommended: between msPOAS and HySCO, using the RBC module) or during model fitting (using the Rician bias correction option in NLLS). Diffusion signal models are fitted on the processed dataset; here, we display the maps of fractional anisotropy (FA) and mean kurtosis tensor (MW) from diffusion kurtosis imaging (DKI). The output from DKI can be used to compute biophysical parameters of the white matter; shown here is the map of Watson concentration parameter (;<) from the WMTI-Watson biophysical model. Note that for the in vivo brain dMRI dataset, the inferior slice displayed contains relatively little white matter; hence, we refrained from using a white matter mask. The less smooth appearance of the *к* map is due to the low values in the gray matter.

### 2.4.1 WMTI-Watson model

The white matter tract integrity (WMTI)-Watson model as an implementation of the Standard Model assumes two non-exchanging water compartments (intra- and extra-axonal tissue water) (Alexander et al., 2019; Novikov et al., 2019). The model characterizes the intra-axonal compartment as “sticks” of zero radius, with an intra-axonal diffusivity *D_a_* and axonal water fraction *f.* Axonal alignment is characterized by the Watson concentration parameter k, where higher values indicate higher axonal alignment, and the orientation dispersion index (ODI), where higher values indicate lower alignment. While *к* and ODI are mathematically related (Mollink et al., 2016), ACID outputs both for convenience. The extra-axonal space is modeled as a homogenous medium, described by an axisymmetrical diffusion tensor with parallel (D_e_ ц) and perpendicular (D_e ±_) extra-axonal diffusivities. The five biophysical parameters *(D_a_, f, к, D_e_* ц, *D_e_*_±_) are derived from the axisymmetric DKI tensor metrics (AD, RD, MW, AW, RW) according to the formulas described in (Jespersen et al., 2018; Novikov et al., 2018). Being derived from the biophysical Standard Model, the estimation of WMTI-Watson biophysical parameters is generally degenerate, which leads to two solutions: the plus branch, which assumes *D_a_ > D_e_* ц, and the minus branch, which assumes *D_a_ < D_e_* ц (Novikov et al., 2018). We recommend using the plus branch (default in the toolbox), as in our experience, and also reported by others (Jelescu et al., 2020; Jespersen et al., 2017), the minus branch is the biologically invalid solution.

### 2.4.2 NODDI-DTI

NODDI-DTI (Edwards et al., 2017) is based on the neurite orientation dispersion and density imaging (NODDI) model (Zhang et al., 2012). While NODDI is a three-compartment biophysical model with intra- and extra-axonal space, and cerebrospinal fluid compartments, NODDI-DTI assumes that the latter compartment can be neglected in normal appearing white matter. NODDI-DTI further assumes a fixed diffusivity of the intra-neurite compartment (). In our implementation, users can either choose from two fixed values tailored for in vivo (=1.7-10 ^3^ mm^2^/s) and ex vivo (=0.6-10 ^3^ mm^2^/s) datasets, or select their own value. NODDI-DTI estimates the intra-neurite (here:) and extra-neurite () signal fraction, as well as the Watson concentration parameter and the orientation dispersion index (ODI), from the FA and MD maps. While WMTI-Watson requires specific multi-shell dMRI data for robust parameter estimation, NODDI-DTI parameters can also be obtained from single-shell DTI acquisitions.

#### 2.5 Utilities

ACID utilizes SPM’s utility functions, available under SPM -> Util in the SPM12 Batch Editor, for handling and manipulating NlfTI images. These functions include mathematical operations on single or multiple images, reorienting images, and concatenating 3D volumes and separating 4D volumes. Additionally, ACID provides its own set of utility functions for image manipulation, mask generation, quality assessment, and other related tasks (refer to Table 2 for more details).

#### 2.6 External tools

ACID provides the option to integrate external tools from other packages, which can be accessed directly from the ACID graphical user interface (GUI) *(External tools* module), ensuring a seamless integration into ACID pipelines. We included the following external tools in the current release: (i) FSL eddy^1^ (J. L. R. Andersson & Sotiropoulos, 2016); (ii) FSL topup^2^ (Smith et al., 2004); (iii) dwidenoise^3^ (based on the Marchenko-Pastur principal component analysis (MP-PCA), part of the MRtrix toolbox) (Veraart et al., 2016); (iv) denoising^4^ (based on the local principal component analysis (LPCA), part of the DWI Denoising Software) (Manjon et al., 2013); (v) Koay’s noise estimation^5^; (vi) mrdegibbs^6^ for Gibbs ringing removal, part of the MRtrix toolbox (Kellner et al., 2016); and (vii) the WMTI model (part of the DESIGNER toolbox) (Fieremans et al., 2011). ACID also allows expert users to incorporate their own external tools into the toolbox, using the aforementioned examples as a template.

#### 2.7 Output structure and BIDS naming convention

ACID supports the BIDS standard, while also being compatible with non-BIDS data. Following BIDS recommendations, ACID appends a label to the output filename’s desc field, which indicates the applied processing step (refer to Table 3 for a list of labels used in the modules *Pre-processing, Diffusion tensor/kurtosis imaging,* and *Biophysical models).* For instance, after applying ECMOCO to sub01_dwi.nii, the output file becomes sub01_desc-ECMOCO_dwi.nii. When multiple processing steps are involved, the labels are concatenated, as in sub01_desc-ECMOCO-msPOAS_dwi.nii. Model fitting appends three labels indicating the type of diffusion model, algorithm, and parametric map, such as sub01_desc-ECMOCO-POAS-DKI-OLS-FA_dwi, nii. For BIDS-compliant input, ACID generates a bval and bvec file after each processing step. ACID stores all output in the derivatives folder, with separate subfolders for each module’s output (e.g., derivatives/POAS-Run). ACID retains the same folder structure and naming convention even when non-BIDS input is provided.

## 3. Results

### 3.1 Pipelines

ACID is fully integrated into the SPM12 batch system, allowing users to execute its functions individually or combined into linear pipelines with multiple steps. Each step can receive the output of any of the previous steps via flexible and easy-to-use dependencies. While pipelines are typically set up in the SPM batch system, they can also be converted into MATLAB code (SPM batch script) for automation and further customization. In addition to its own functions, ACID integrates seamlessly with a range of standard SPM features, including segmentation, co-registration (based on affine transformation), spatial normalization (including non-linear registration), and voxel-based statistical analyses, as well as a growing number of SPM extensions^7^. For example, combining ACID with the hMRI toolbox enables multi-contrast analysis of dMRI and other quantitative MRI data, such as relaxation rates (Tabelow et al., 2019).

### 3.2 Example applications

To demonstrate the application of ACID toolbox on different types of dMRI data, here we provide three example pipelines for in vivo brain, in vivo spinal cord, and ex vivo dMRI (Fig. 3). Details of these three datasets are summarized in Table 4. The gradient schemes used for all datasets were based on the configurations proposed by (Caruyer et al., 2013), available online^8^. The design of the sampling schemes followed a uniform coverage on a sphere. Note that data with reverse phase­encoding direction were available for all three datasets, which refers to the acquisition of either a single bO volume or all volumes with identical geometry and sequence parameters but opposite phase encoding direction. All example pipelines consist of artifact correction (ECMOCO, msPOAS, RBC, HySCO) and model fitting steps. While Gibbs ringing removal is often part of dMRI processing pipelines (Ades-Aron et al., 2018; Kellner et al., 2016; Tournier et al., 2019) and is also available in ACID as an external tool, we refrained from including it in the example pipelines because the interaction between denoising and the interpolation associated with Gibbs ringing removal is not well characterized yet. We emphasize that these example pipelines might not be optimal for all cases; users might find that another combination of pre-processing steps, which might also include Gibbs ringing removal, works even better for their data.

While the pipelines for in vivo brain, in vivo spinal cord, and ex vivo dMRI follow similar concepts, recommended settings for each region may differ (Table 5). It is important to note that the settings listed in Table 5 serve as initial values for typical datasets. The optimal settings for a particular dataset depend on the sequence parameters, the subject, and the imaged region. Model fitting may be followed by spatial processing, such as co-registration to the structural image or spatial normalization to a template in a standard space (e.g. MNI152 space), and statistical analysis (e.g., ROI- or voxel-based analysis).

## 4. Discussion

We have developed the ACID toolbox, which extends the capabilities of the SPM framework by providing comprehensive pre-processing and model fitting techniques for in vivo brain, spinal cord, and ex vivo dMRI data. Besides commonly used diffusion signal models such as DTI and DKI, ACID also offers biophysical models that provide parameters of white matter tissue microstructure such as axonal water fraction and axon orientation dispersion. Being seamlessly integrated into the SPM batch system, ACID allows for user-friendly access to SPM’s powerful spatial processing tools and statistical framework. In addition to offering recommended pipelines for in vivo brain, spinal cord, and ex vivo dMRI, ACID provides the flexibility for users to create customized pipelines tailored to their specific data. Adhering to the BIDS conventions facilitates data sharing, enhances data comprehension for investigators, and makes ACID compliant with software requiring BIDS input (https://bids-apps.neuroimaging.io).

### 4.1 Pre-processing dMRI data

ACID offers artifact correction steps typically applied to dMRI data, including image realignment (ECMOCO), adaptive denoising (msPOAS), Rician bias correction (RBC), and correction for susceptibility-induced geometric distortions (HySCO). Here, we discuss specific considerations regarding their use for various applications.

Correcting for displacements within the dMRI data through image realignment is one of the most important but also challenging tasks. ECMOCO provides users with the flexibility to choose the degrees of freedom for image realignment based on the anticipated type of displacement, but also offers a selection of pre-defined degrees of freedom that are optimized for brain, spinal cord, and ex vivo dMRI.

In brain dMRI, motion can be approximated as a rigid body displacement with 6 degrees of freedom (DOF). Eddy-current spatial displacements, to a first-order approximation, result in translation and scaling along the phase-encoding direction (typically, the y-axis), and in-plane and through-plane shearing (Mohammadi et al., 2010). Since these displacements affect the entire brain, we recommend employing a 9-DOF volume-wise (volume to volume) registration with translation and rotation along x, y, and z, scaling along y, and shearing in the x-y and y-z plane. First-order approximation of eddy-current displacements might not always be sufficient, as dMRI data can also be affected by higher-order eddy-current field inhomogeneities causing non-linear distortions (J. L. R. Andersson & Sotiropoulos, 2016; Rohde et al., 2004). For example, in our observations, ECMOCO was not effective in removing pronounced eddy-current displacements present in the dMRI data of the Human Connectome Project (Van Essen et al., 2012). In such cases, we recommend using FSL eddy, which incorporates higher-order eddy-current correction terms (J. L. R. Andersson & Sotiropoulos, 2016) and can be called directly from ACID as an external tool (Section 2.6). In cases where ECMOCO is sufficient, an advantage of ECMOCO is that its performance is largely independent of the number of diffusion directions, whereas FSL eddy requires a minimum number of diffusion directions for good performance (see FSL website^9^ for recommendations).

In spinal cord dMRI, volume-wise registration has been found to be less effective (Cohen-Adad et al., 2009; Mohammadi et al., 2013) due to displacements that vary along the rostro-caudal axis of the spinal cord. These displacements appear mostly in the phase-encoding direction and are caused by physiological factors such as respiration and cardiac pulsation (Kharbanda et al., 2006; Summers et al., 2006). We recommend applying volume-wise registration for rough alignment and correction of through-slice displacements, followed by slice-wise (slice to slice) registration for correcting any remaining slice-dependent displacement. This combined approach has demonstrated effectiveness in realigning not only volumes but also individual slices (Fig. B2), as well as improving the contrast-to-noise ratio between gray and white matter and reducing test-retest variability in DTI maps of the spinal cord (Mohammadi et al., 2013). Eddy-current distortions are typically less severe in the spinal cord compared to the brain, because the in-plane field of view is smaller and located near the scanner isocenter. This makes the first-order approximation of eddy-current displacements, as supported by ECMOCO, generally adequate. We recommend employing a 4-DOF volume-wise registration (translation along x, y, z; scaling along y) followed by a 3-DOF slice-wise registration (translation along x, y; scaling along y). In datasets with low SNR, slice-wise correction along x can be omitted, given the smaller range of movement which makes reliable estimation difficult. We advise against correcting for in-plane rotation and shearing, as their expected range is very small. Correction for these DOFs might introduce spurious displacements during realignment, a risk we consider greater than not applying correction at all.

Structures surrounding the spinal cord (bones, ligaments, etc.) may move independently from the spinal cord, potentially leading to inaccuracies in transformation parameters. Moreover, as these structures typically occupy a larger portion of the image, they can dominate the estimation of transformation parameters. To address this challenge, ECMOCO provides the option of specifying a spinal cord mask to restrict the estimation of transformation parameters to the spinal cord and its immediate surroundings (Fig. 3). Any residual misalignments can be manually corrected using the *Slice-wise realignment* utility function (Table 2).

In ex vivo dMRI, specimen motion is not anticipated if the specimen is appropriately fixed, for instance, by using a sample holder or embedding it in agarose. Thus, we recommend correcting only for the four first-order eddy-current displacements (y-translation, y-scaling, x-y shearing, y-z shearing). The first-order approximation is typically adequate for small specimens where eddy-current displacements are not severe.

In general, the performance of msPOAS and HySCO is largely independent of the anatomical features present in the image; therefore, default parameters are expected to work well for in vivo brain, spinal cord, and ex vivo dMRI data. Nevertheless, the default regularization parameters for HySCO (alpha “diffusion” and beta “Jacobian” regulator), accessible through the script `config/local/acid_local_defaults.m,` are optimized for the brain and may require adjustment for the spinal cord if performance is inadequate.

Applying HySCO is particularly important for acquisitions with severe susceptibility-related distortions, such as multi-band EPI without parallel imaging, and for multi-contrast analyses where dMRI data or other quantitative maps are combined with structural reference images, e.g., the dMRI-based axonal water fraction and magnetization transfer saturation maps in g-ratio mapping (Mohammadi & Callaghan, 2021) or multi-contrast MRI in the spinal cord (David et al., 2019). In these cases, HySCO improves the overlap between the undistorted structural image and the dMRI data, improving the performance of subsequent co-registration and spatial normalization algorithms. HySCO has also been shown to improve the accuracy of g-ratio mapping (Clark et al., 2021; Mohammadi et al., 2015b). While HySCO is far more efficient than FSL topup in terms of computation time (Macdonald & Ruthotto, 2018), it does not integrate movement and susceptibility artifact correction into a single model. To mitigate the effects of subject movement, we propose acquiring images with reversed phase-encoding direction (the blip-up and blip-down images) in close succession.

The application of adaptive denoising (msPOAS) is important as it reduces the variance and therefore improves the precision of the tensor and kurtosis parameter estimates (see Fig. S4 for an example illustrating the effect of msPOAS on DKI parameters, and refer to Becker et al., (2014) for more examples and details). For high-SNR data, denoising might not be advantageous; instead, denoising methods could even introduce additional error (see analysis in Appendix G). For low-SNR data, Rician bias correction (RBC), either applied to the dMRI data or during model fitting, must be performed in addition to msPOAS to mitigate the Rician bias in parameter estimates (see Appendix F for an example). An in-depth analysis of the impact of Rician bias correction on DKI and axisymmetric DKI can be found in Oeschger et al., 2023a.

### 4.2 Model fitting on dMRI data

#### 4.2.1 Physical diffusion models

At a given b-value, the SNR in spinal cord dMRI is typically lower than in brain dMRI due to (i) the smaller cross-sectional area that requires higher in-plane resolution (see Fig. 4A for a size comparison), (ii) the high signal attenuation for diffusion-gradient directions parallel to the highly aligned fibers in the head-feet direction (Fig. 4B), and (iii) the suboptimal coil configuration in the thoracic and lumbar regions, which are not covered by the head and neck coil. Lower SNR increases the variance of parameter estimates and makes spinal cord dMRI more susceptible to Rician bias. Consequently, SNR is often prohibitively low at higher b-values necessary for fitting the kurtosis tensor, making the application of DKI in the spinal cord very challenging.

**Fig. 4.**
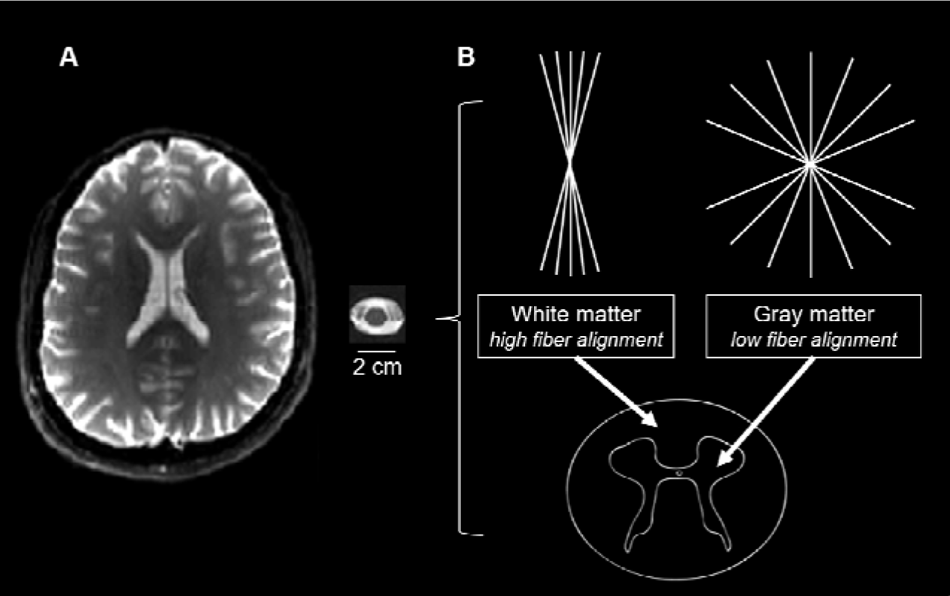
(A) Illustration of differences in the cross-sectional area between the brain and spinal cord, displaying a single axial slice of the mean T2-weighted (bO) image (refer to Table 4 for details on the datasets). (B) Schematic visualization of the spinal cord, highlighting the “butterfly-shaped” gray matter, which is located in the middle of the spinal cord and contains neuronal cell bodies and loosely aligned fibers, and the surrounding white matter, which contains highly aligned fibers.

The bias in parameters estimates induced by signal outliers from cardiac, respiratory, and other physiological artifacts (Mohammadi, Hutton, et al., 2013) can be mitigated by applying robust fitting as a tensor fitting method (Appendix E.3). Given the higher occurrence of signal outliers in the spinal cord, robust fitting holds particular relevance for spinal cord dMRI. In a previous study, we demonstrated that robust fitting leads to higher FA values within the white matter and lower FA values within the gray matter in spinal cord dMRI data, resulting in an approximately 8% enhancement in contrast-to-noise ratio (Mohammadi, Freund, et al., 2013). While robust fitting demonstrated high resistance to contamination (presence of outliers) compared to OLS and NLLS estimations, it is important to note that robust fitting requires a sufficiently large number of artifact-free data points. Simulations suggested that robust tensor estimates begin to break down when the frequencies of moderately intense cardiac pulsation artifacts exceed 27-30% (Zwiers, 2010; Fig. 5).

**Fig. 5.**
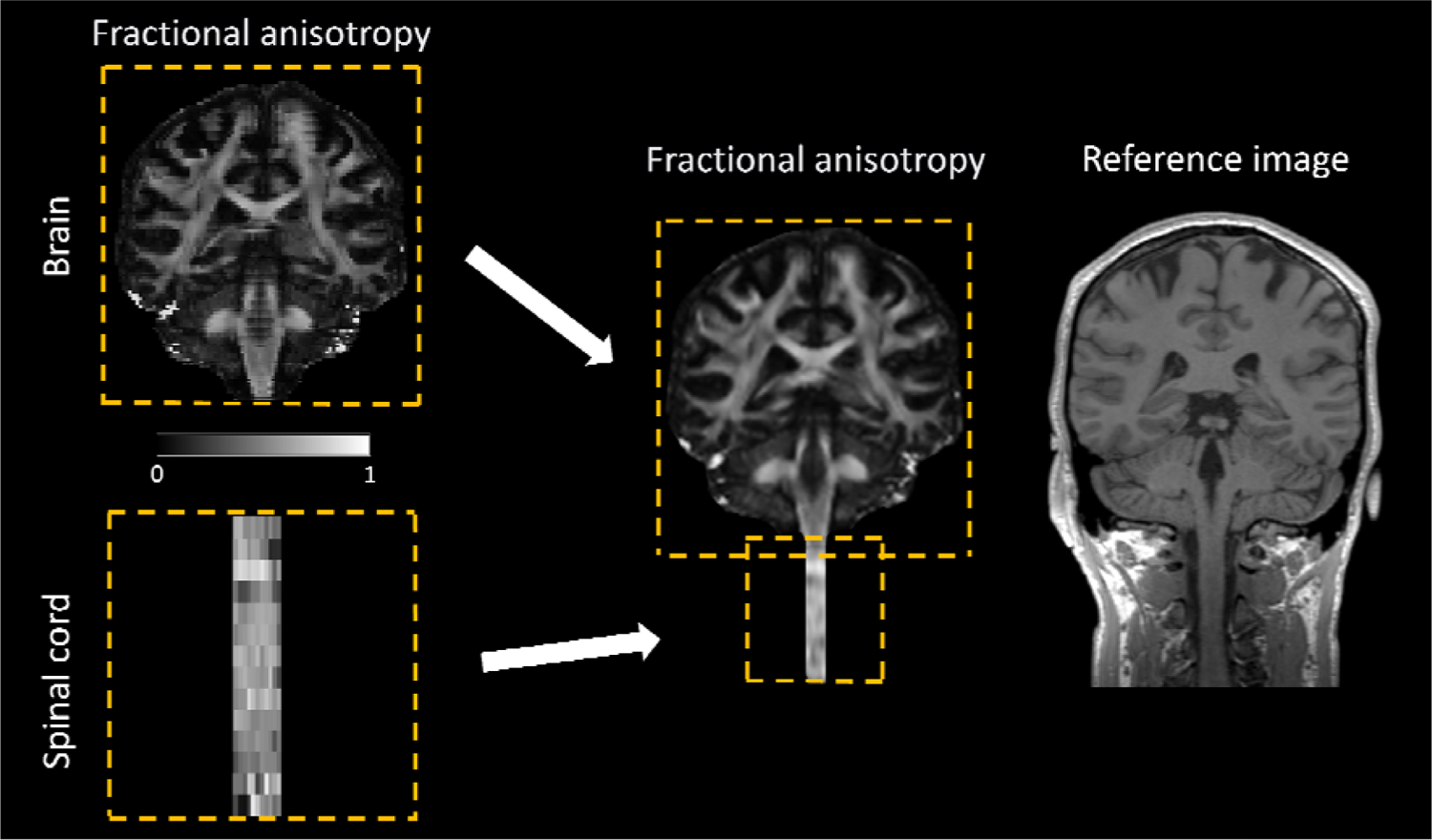
Merging of two fractional anisotropy (FA) maps, covering the brain and cervical cord, respectively, into a unified FA map using the Fusion utility function (Table 2). The two images should ideally share an overlapping region, but they may have different geometric properties such as resolution and number of slices. In the overlapping region, the voxel intensity values are computed as the average of the intensity values from the two images. The merging process requires a structural image as the registration target. The combined FA map is resampled onto the higher-resolution structural image, resulting in a smoother appearance.

One potential limitation of linearized fitting methods is their operation on logarithmically transformed signals, where the assumption of Gaussian (or Rician) error distribution may not hold. The presence of logarithmically distorted Rician noise distribution not only restricts validity but can also impact the accuracy of the parameter estimates (J. L. R. Andersson, 2008; Chang et al., 2005; Koay et al., 2006), particularly in the low-SNR regime such as in spinal cord dMRI. The WLS and robust fitting algorithms incorporate the signal intensity into the weights of the estimator function (Appendix E.2 and E.3), which was shown to reduce the effect of log-Rician distortion (Salvador et al., 2005). Alternatively, the NLLS algorithm (Appendix E.4) can be used, which circumvents the distortion of the Rician distribution by operating on the original (non-logarithmic) signals, and is therefore expected to yield more accurate parameter estimates, provided that the numerical fitting problem is sufficiently well-conditioned.

In summary, for data with relatively high SNR and a frequent occurrence of outliers, we recommend using robust fitting to mitigate the influence of outliers. NLLS, particularly when combined with Rician bias correction, may be more suitable for dMRI data with lower SNR, which is often encountered in acquisitions for DKI (refer to Oeschger et al., 2023a for recommended minimum SNR values and the *Rician bias simulation* utility function in Table 2 for simulating the Rician bias on dMRI data with a given SNR). Low-SNR data with a frequent occurrence of outliers pose challenges for model fitting, where a combination of msPOAS with RBC might reach their limits. In such cases, reliability masking can assist in identifying and excluding corrupted, thus unreliable, voxels from the parameter maps (David et al., 2017).

#### 4.2.2 Biophysical diffusion models

Of the biophysical models implemented in ACID, WMTI-Watson relies on DKI metrics (requiring at least two diffusion shells), while NODDI-DTI relies on DTI metrics (requiring a single diffusion shell only). This implies that the challenges associated with the estimation of DTI and DKI metrics, as discussed earlier, also apply to derived biophysical models. Accurate and precise estimation of DKI and DTI metrics is essential for the successful application of WMTI-Watson and NODDI-DTI, respectively.

In general, we recommend the DKI-based WMTI-Watson model over NODDI-DTI due to the fewer model assumptions, allowing it to better capture diffusion patterns in complex axonal configurations within the brain white matter. This aligns with the results from our example multi­shell brain dMRI dataset, where WMTI-Watson yielded more accurate estimates of κ and AWF compared to NODDI-DTI (Fig. S5). For a more in-depth comparison of biophysically-derived values with histological values, refer to Papazoglou et al., 2023.

On the other hand, complex models are more “data-hungry” and more susceptible to noise due to the higher number of fitted parameters, which can lead to poorly conditioned optimization problems when the amount and/or the quality of input data are insufficient. Therefore, for low-SNR data, as is often the case in spinal cord dMRI, the less complex but better-conditioned NODDI-DTI model might be the preferred choice. The low b-values often used in spinal cord could also lead to inadequate parameter estimation when using the WMTI-Watson model. Indeed, NODDI-DTI yielded a more accurate estimation of κ in the example spinal cord dMRI dataset, whereas WMTI-Watson highly overestimated it (Fig. S5). A drawback of the NODDI-DTI model in the spinal cord is its assumption of fixed intra- and extra-cellular diffusivities, which were optimized for the brain and might not be valid for the spinal cord. Both low SNR (Veraart et al., 2011) and kurtosis bias (Edwards et al., 2017) can lead to an underestimation of MD. The lower SNR can also lead to an underestimation of MD due to kurtosis bias (Fig. S3), impacting the model parameter estimation when MD falls outside the range where the NODDI-DTI model provides a valid representation (refer to Equation (4) in Edwards et al., 2017). This was evident in the estimation of AWF, which proved unfeasible in the spinal cord dataset (see Figs. FI and S5). We anticipate that future improvements in acquisition methods will enhance the SNR in spinal cord dMRI, enabling the acquisition of higher fa-values, which would alleviate many of the above-mentioned drawbacks.

A compromise between these two models could be the WMTI model, which is included as an external tool in ACID (Section 2.6). WMTI assumes highly aligned fibers, which holds true in white matter regions with high fiber alignment, such as the corpus callosum or the spinal cord white matter, but is less appropriate in regions with more complex axonal configurations, such as parts of the superior longitudinal fasciculus.

Ex vivo neuronal tissues exhibit different diffusivities compared to in vivo tissues due to various factors, including the effect of fixation, changes in chemical properties, and differences in temperature and composition of the embedding fluid. For example, white matter diffusivity was reported to reduce by approximately 85% from in vivo to ex vivo conditions, while the ratio between gray and white matter diffusivities remains similar, around 2-3 (Roebroeck et al., 2019). To accommodate the reduced diffusivities under ex vivo conditions, ACID offers the option to utilize compartmental diffusivities tailored for ex vivo datasets within the NODDI-DTI model. Such an adjustment is not necessary for WMTI and WMTI-Watson, as their compartmental diffusivities are fitted rather than fixed.

We emphasize that NODDI-DTI, WMTI, and WMTI-Watson have been developed to characterize diffusion in the white matter. Recently, several efforts have been made to extend biophysical models to the gray matter (Jelescu et al., 2020). Notable examples include the SANDI (Palombo et al., 2020) and NEXI (Jelescu et al., 2022) biophysical models. However, these models thus far, no study using these protocols on a clinical MRI system has been published.

### 4.3 Studies quantitatively evaluating the performance of ACID pipelines

Here, we briefly summarize and discuss the studies that quantitatively evaluated the performance of ACID tools individually or in comparison with other tools.

#### 4.3.1 Evaluating pre-processing pipelines

In a previous study, we assessed the performance of ECMOCO as well as the combination of ECMOCO and msPOAS in simulated high- and low-SNR multi-shell brain dMRI datasets with added motion and eddy current artifacts (i.e., perturbed data) (Mohammadi, Tabelow, et al., 2015). We found that the performance of ECMOCO in correcting the perturbed volumes was dependent on SNR, with the number of incorrectly registered volumes increasing at lower SNR (SNR < 16). However, the combined application of msPOAS and ECMOCO effectively reduced the number of incorrectly registered volumes even at low SNR (Mohammadi, Tabelow, et al., 2015; Fig. 3). Additionally, correcting the perturbed volumes with ECMOCO and msPOAS yielded FA maps closer to the “ground truth”, i.e., the FA map computed on the unperturbed data (Mohammadi, Tabelow, et al., 2015; Fig. 5). In another study utilizing clinical spinal cord dMRI data, we evaluated the impact of ECMOCO on the group differences observed in FA between patients with degenerative cervical myelopathy and healthy controls (David et al., 2017; Fig. 7). Our analysis revealed that ECMOCO had only a minimal effect on the two-sample t-score computed between the FA values of the two groups.

We also tested the effects of different denoising methods (msPOAS, LPCA, and MP-PCA) on the accuracy of DKI metrics, with the details and results described in Appendix G. In short, we found that denoising (using any of the three methods) is beneficial only in the low-SNR domain (below an SNR of approximately 30). In high-SNR data, denoising did not lead to further improvements with MP-PCA and even introduced additional errors with msPOAS and LPCA. In terms of susceptibility artifacts, we previously found in a brain dMRI dataset that FSL topup was more efficient in correcting susceptibility-related distortions than HySCO, even when including a motion correction step between the reverse phase-encoded (blip-up and blip-down) images before running HySCO (Clark et al., 2021; Fig. 3). This is potentially because the HySCO pipeline involved multiple interpolation steps, introducing additional blurring effects, while FSL topup incorporates motion and susceptibility distortion correction within the same model. The same study found that combining reverse phase-encoded images using the “weighted average” method *(HySCO: combine blip-up and blip-down images* module), as opposed to the “arithmetic average” method, reduces image blurring in the corrected brain dMRI data and achieves greater overlap between the dMRI data and the corresponding structural image. In fact, when using the “weighted average” method, HySCO performed comparably to FSL topup and even outperformed it in regions suffering from high levels of distortion (Clark et al., 2021; Fig. 5). In spinal cord dMRI, a previous study found that HySCO is comparable to other distortion correction tools such as FSL topup (Schilling et al., 2024).

#### 4.3.2 Evaluating diffusion signal models

In brain dMRI datasets, we found that robust tensor fitting can reduce the effect of signal outliers due to motion, eddy current artifacts, incorrectly registered volumes (Mohammadi, Tabelow, et al., 2015; Fig. 5C-D), or physiological noise (Mohammadi, Hutton, et al., 2013; Fig. 9). In spinal cord dMRI, we quantified the performance of robust fitting and showed that it can reduce the bias in FA, especially at tissue boundaries (Mohammadi, Freund, et al., 2013; Fig. 7). On the other hand, robust fitting had only a minor effect on group differences in FA between patients with degenerative cervical myelopathy and healthy controls, regardless whether using the ACID implementation of robust fitting or using RESTORE (part of the CAMINO toolbox, Chang et al., 2012) (David et al., 2017; Fig. 7). However, within the same study, we also found that supplementing the pipeline with reliability masking to exclude outlier voxels (Section 2.5) considerably increased the statistical differences between patients and controls (David et al., 2017; Fig. 7)).

### 4.4 Applications

For all applications, it is highly recommended to assess the data quality before and after each processing step. In addition to the quality assessment utility functions *DWI series browser* and *DWI series movie* (Table 2), multiple ACID modules generate diagnostic plots to identify the presence and type of artifacts in the dMRI data. Example diagnostic plots are provided in Figs. S1-S2.

#### 4.4.1 Integration with SPM modules

ACID can be readily combined with SPM tools for segmentation, spatial processing, and voxel-based analysis of parametric maps. Segmenting the brain or spinal cord is often necessary for co­registration, spatial normalization, or tissue-specific analyses. In the brain, tissue probability maps of white matter, gray matter, and cerebrospinal fluid can be created by unified segmentation, the default segmentation routine in SPM12 (Ashburner & Friston, 2005). These tissue probability maps can also be used to create a binary brain mask using the *Create brain mask* utility function (Table 2). To enable SPM’s unified segmentation in the spinal cord, the brain tissue priors need to be substituted with the joint brain and spinal cord tissue priors from the probabilistic brain and spinal cord atlas (Blaiotta et al., 2017). However, this atlas only covers the upper cervical cord down to C3; for other spinal levels, the user is referred to automatic (e.g., deepseg (Perone et al., 2018)) or semi­automatic (e.g., active surface method (Horsfield et al., 2010)) segmentation techniques.

Brain dMRI data can be co-registered to the corresponding structural image using *spm_coreg.* For non-linear spatial registration to the MNI space, we recommend SPM DARTEL (Ashburner, 2007) or Geodesic Shooting (Ashburner & Friston, 2011). As SPM registration tools often rely on brain tissue priors, they cannot be applied directly on spinal cord dMRI. For the spinal cord, we recommend utilizing the PAM50 template (De Leener et al., 2018) and the corresponding normalization tools integrated into the Spinal Cord Toolbox (De Leener et al., 2017).

While brain and spinal cord images are typically analyzed separately, there are scenarios where combining them into a single image can be beneficial. For example, when registering the brain and spinal cord image to a joint brain-spinal cord template, such as the probabilistic atlas of the brain and spinal cord (Blaiotta et al., 2017), the warping field is often obtained using a structural image with a large field of view (FOV) covering both regions (Fig. 5). To apply this warping field to the brain and spinal cord images, they need to be fused into a single image. ACID provides the *Fusion* utility function (Table 2) which merges two distinct images, acquired with different FOV and geometric properties, into a unified large-FOV image (Fig. 5).

**Table 4.** Scan parameters of the in vivo brain, in vivo spinal cord, and ex vivo dMRI datasets used in this paper.

| *Dataset* | **In vivo brain** | **In vivo spinal cord** | **Ex vivo specimen** |
| --- | --- | --- | --- |
| *Imaged body part or tissue* | entire brain (incl. cerebellum) of a 34-year-old healthy volunteer | upper cervical cord (appr. C1-C4) of a 43-year-old healthy volunteer | ex vivo specimen of the temporal lobe from a 46-year-old patient diagnosed with drug-resistant temporal lobe epilepsy; specimen embedded in glucose for 2h and fixed with 4% paraformaldehyde for 12h before measurement |
| *Scanner* | 3T Siemens Prisma Fit | 3T Siemens Prisma Fit | 3T Siemens Prisma Fit |
| *Receive coils* | 64-channel Head/Neck | 64-channel Head/Neck | 16-channel Hand/Wrist |
| *Sequence* | 2D single-shot spin-echo EPI | 2D single-shot spin-echo EPI | pulse gradient spin echo |
| *Volumes and b-values [s/mm<sup>2</sup>] (number of gradient directions)* | b=0 (18); b=600 (30); b=1100 (45); b=2500 (60) | b=0 (11); b=500 (30); b=1000 (30); b=1500 (30) | b=0 (36); b=550 (30); b=1100 (75); b=2200 (45); b=2500 (60); b=5000 (60) |
| *Cardiac gating* | - | 2 slices per cardiac cycle, trigger delay of 260 ms | - |
| *Number of slices* | 100 (interleaved, no gap) | 14 (interleaved, no gap) | 160 |
| *Resolution [mm<sup>3</sup>]* | 1.7 x 1.7 x 1.7 | 1.0 x 1.0 x 5.0 | 0.8 x 0.8 x 0.8 |
| *Field of view [mm<sup>3</sup>]* | 204 x 170 x 201 | 128 x 36 x 70 | 128 x 48 x 48 |
| *Echo time* | 75 ms | 73 ms | 99 ms |
| *Repetition time* | 5800 ms | pulse-dependent (cardiac gated) | 8700 ms |
| *Parallel imaging* | 2x (GRAPPA) | - | - |
| *Multi-band imaging* | - | - | - |
| *Phase partial Fourier* | 7/8 | - | 7/8 |
| *Phase-encoding dir.* | A-P | A-P | A-P |
| *Readout bandwidth* | 1842 Hz/pixel | 1396 Hz/pixel | 802 Hz/pixel |
| *EPI spacing* | 0.77 ms | 0.93 ms | 1.37 ms |
| *EPI factor* | 120 | 36 | 60 |
| *Acquisition time [min:sec]* | 17:46 | 06:51 (nominal) | 93:10 |
| *Additional data with reversed phase-encoding direction* | a single b0 volume acquired with reversed phase-encoding direction | full blip-reversed acquisition (reversed phase-encoding available for each volume) | full blip-reversed acquisition (reversed phase-encoding available for each volume) |

**Table 5.**
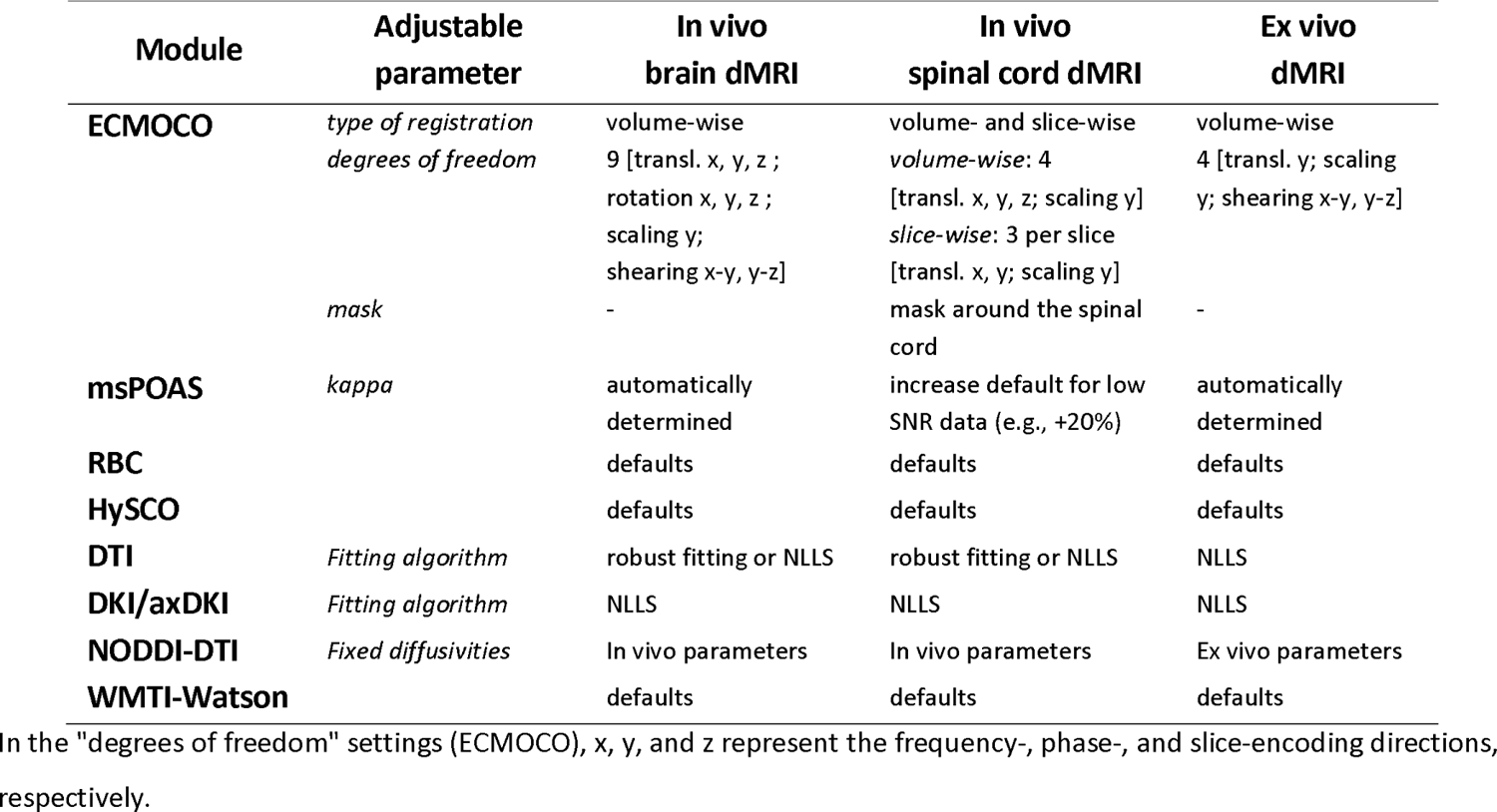
Settings of selected modules for in vivo brain, in vivo spinal cord, and ex vivo dMRI datasets.

ACID benefits from SPM’s rich statistical framework for voxel-based analysis. SPM’s second-level analysis tool (SPM -> Specify 2nd-level) performs voxel-based statistical tests on the parametric maps using t-test, ANOVA, or general linear model. In the SPM -> Results module, the framework also offers (i) multiple comparison correction in the form of family-wise error rate and false discovery rate, (ii) thresholding the test statistics at cluster- and voxel-level and providing a list of significant clusters/voxels, and (iii) various visualization tools for displaying and saving the significant clusters. Furthermore, ACID’s *ROI analysis* utility function (Table 2) can be used to extract mean metrics within subject-specific ROIs in the native space or perform atlas-based analysis in the template space. For atlas-based analysis in the spinal cord, the user is referred to the PAM50 white and gray matter atlas (De Leener et al., 2018).

Although ACID does not provide tractography or tract-based analysis tools, the output of its model fitting methods can be input into tractography tools such as FSL or the SPM12-based Fibertools toolbox (see Wiki^11^ on the git repository for more details).

#### 4.4.2 Computation time

To speed up the processing and analysis of dMRI data, parallel computing is implemented wherever applicable. This technique can substantially accelerate the most time-consuming ACID modules, including ECMOCO and DTI/DKI fit. Note that parallel computing requires the Parallel Computing Toolbox in MATLAB. Table 6 provides the computation times for selected ACID functions on a typical brain and spinal cord dMRI dataset.

**Table 6.** Computation times of selected ACID modules on an example In vivo brain and In vivo spinal cord dMRI dataset (refer to Table 4 for details on the datasets), when run on a MacBook Ml laptop (4 cores, 16 GB RAM).

| Module | In vivo<br>brain dMRI | In vivo<br>spinal cord dMRI |
| --- | --- | --- |
| ECMOCO | 9 min | 2 min |
| msPOAS | 92 min | 1 min |
| RBC | < 1 min | < 1 min |
| HySCO | 2 min | 1 min |
| DKI (using NLLS) | 4 min | 2 min |
| WMTI-Watson | < 1 min | 1 min |

#### 4.4.3 Research applications

ACID has been used in a variety of clinical and neuroscience research, e.g., in dMRI studies assessing cerebral changes in patients with multiple sclerosis (Deppe et al., 2016a, 2016b; Dossi et al., 2018; Kugler & Deppe, 2018) and Parkinson’s disease (Szturm et al., 2021), and to assess gliomas (Paschoal et al., 2022; Raja et al., 2016). We have also used ACID to investigate spinal cord white matter following spinal cord injury (Büeler et al., 2024; David et al., 2019, 2021, 2022; Grabher et al., 2016; Huber et al., 2018; Seif et al., 2020; Vallotton et al., 2021). A non-comprehensive list of studies using the ACID toolbox can be found on the project website^10^. Note that certain ACID functions can be applied to MRI data beyond dMRI as well; for instance, HySCO has been used to correct brain fMRI data for susceptibility artifacts (De Groote et al., 2020). It is important to note that ACID has not been approved for clinical applications by any health agency and it comes with no warranty. Therefore, it should not be used for diagnosis in clinical settings.

### 4.5 Limitations and future directions

Comparing the tools within the ACID toolbox with alternative implementations in other software presents challenges because their performance depends on the specific dMRI data and the chosen parameter settings from a potentially large parameter space, which necessitates a systematic exploration of the parameter space. In addition, the evaluation of entire processing pipelines would drastically increase the number of parameters to test. While we have outlined the comparisons conducted so far in Section 4.3, we assert that a thorough quantitative comparison between toolboxes warrants a dedicated future study. In general, we encourage users to undertake such comparisons on their own datasets.

The ACID toolbox is the result of a collaborative effort to extend the SPM ecosystem with state-of-the-art processing and modeling tools for dMRI data. Our aim is to make the toolbox widely accessible, leveraging SPM’s large and vibrant community. Users can submit their questions, bug reports, and suggestions via the dedicated mailing list or by opening an issue on the git website. This paper offers an overview of the current state of the toolbox, with several ongoing developments not covered here. The modularity of the toolbox allows for integration of newly developed methods, even when used concurrently with old ones. Biophysical modeling is an emerging field, and we expect many methodological advancements to occur in the coming years. To align with this ongoing development, our goal is to consistently integrate state-of-the art biophysical models into ACID. We also plan to add the Rician maximum likelihood estimator (Sijbers et al., 1998) as an alternative to the existing quasi-likelihood estimators (Polzehl & Tabelow, 2016).

## 5. Conclusion

ACID is an open-source extension to SPM12 that provides a comprehensive framework for processing and analyzing in vivo brain, spinal cord, and ex vivo dMRI data. The toolbox was developed to meet the increasing demand for studies involving spinal cord dMRI, research employing biophysical models, and validation studies utilizing ex vivo dMRI. ACID leverages the core SPM tools and other SPM extensions, which can be easily integrated into the ACID pipeline.

## Ethics statement

Three dMRI datasets from previous studies were re-used in this paper. These studies complied with the principles of the Declaration of Helsinki and were approved by the local ethics committee (Ärztekammer Hamburg). The whole-brain dataset was measured in vivo (ethics approval number: PV5600). The dataset of the temporal lobe specimen was acquired ex vivo (PV5034). The spinal cord dataset was measured in vivo (PV5141).

## Data and Code Availability

The source code of ACID is freely available at https://bitbucket.org/siawoosh/acid-artefact-correction-in-diffusion-mri/src/master/. The authors will make the raw data used for the visualizations in this article available in an associate publication.

## Author Contributions

**Gergely David:** Conceptualization, Data curation, Formal analysis, Investigation, Methodology,

Resources, Software, Visualization, Writing - original draft, Writing - review & editing

**Björn Fricke:** Conceptualization, Data curation, Formal analysis, Investigation, Methodology, Software, Validation, Visualization, Writing - original draft, Writing - review & editing

**Jan Malte Oeschger:** Formal analysis, Methodology, Software, Writing - original draft, Writing - review & editing

**Lars Ruthotto:** Methodology, Software, Writing - review & editing

**Francisco Javier Fritz:** Data curation, Resources

**Ora Ohana:** Data curation, Resources’

**Laurin Mordhorst:** Software

**Thomas Sauvigny:** Data curation, Resources

**Patrick Freund:** Conceptualization, Project administration, Writing - review & editing

**Karsten Tabelow:** Conceptualization, Investigation, Methodology, Project administration, Software, Writing - review & editing

**Siawoosh Mohammadi:** Conceptualization, Formal analysis, Funding acquisition, Investigation, Methodology, Project administration, Resources, Software, Supervision, Writing - original draft, Writing - review & editing

## Acknowledgments

This work was supported by the German Research Foundation (DFG Priority Program 2041 “Computational Connectomics” (MO 2397/5-1, MO 2397/5-2)), the Emmy Noether Stipend (MO 2397/4-1 and 2397/4-2), the BMBF (01EW1711A and B) in the framework of ERA-NET NEURON, and the ERC (Acronym: MRStain, Grant agreement ID: 101089218, DOI: 10.3030/101089218). Views and opinions expressed are however those of the authors only and do not necessarily reflect those of the European Union or the European Research Council Executive Agency. Neither the European Union nor the granting authority can be held responsible for them. L.R. is supported in part by NSF awards (DMS 1751636 and DMS 2038118). P.F. is funded by an SNF Eccellenza Professorial Fellowship grant (PCEFP3_181362/1).

## Appendix A. Implementation and organization

### Appendix A.1. Installation and toolbox documentation

The ACID toolbox is an extension of SPM12 that requires existing MATLAB and SPM12 installations. To run the toolbox without a Matlab license, ACID is also available as a compiled standalone version which only requires MATLAB Runtime (David et al., 2024). The toolbox has been developed and tested with MATLAB versions R2017b to R2024a, and SPM12 from versions r6906 onwards. It is recommended to use the latest SPM release, which can be downloaded from the SPM website^11^, as developments in ACID are synchronized with those in SPM.

Information about the toolbox can be found on the main project website^12^. The source code is available on Bitbucket^13^, where the latest version as well as all previous versions of the toolbox can be downloaded. There are four ways to install the toolbox: (i) by cloning the repository (recommended for staying up-to-date with the latest release), (ii) by downloading the toolbox as a zip file and placing the unzipped directory into the `spm12/toolbox` directory, (iii) by downloading the toolbox as a zip file and using a redirection script that enables switching between different local versions of ACID, or (iv) by downloading the compiled standalone version. The full documentation of the toolbox is available as a Wiki on the git repository^14^, which provides detailed installation instructions, module descriptions, and step-by-step instructions for typical analysis pipelines.

ACID is free but copyrighted software, distributed under the terms of the GNU General Public License as published by the Free Software Foundation (either version 2 of the License or, at your option, any later version). Further details on “copyleft” can be found at the GNU website^15^. It should be noted that ACID is supplied as is and no formal support or maintenance is provided. The toolbox was developed for academic research purposes only and comes with no warranty, nor is it intended for clinical and diagnostics use.

### Appendix A.2. Organization of the toolbox

The ACID modules can be found in the SPM12 Batch Editor by navigating to `SPM -> Tools -> ACID Toolbox.` The toolbox is divided into six modules, as shown in Fig. Al: *Startup, Pre­processing, Diffusion tensor/kurtosis imaging, Biophysical models, Utilities,* and *External tools*.

**Fig. A1.**
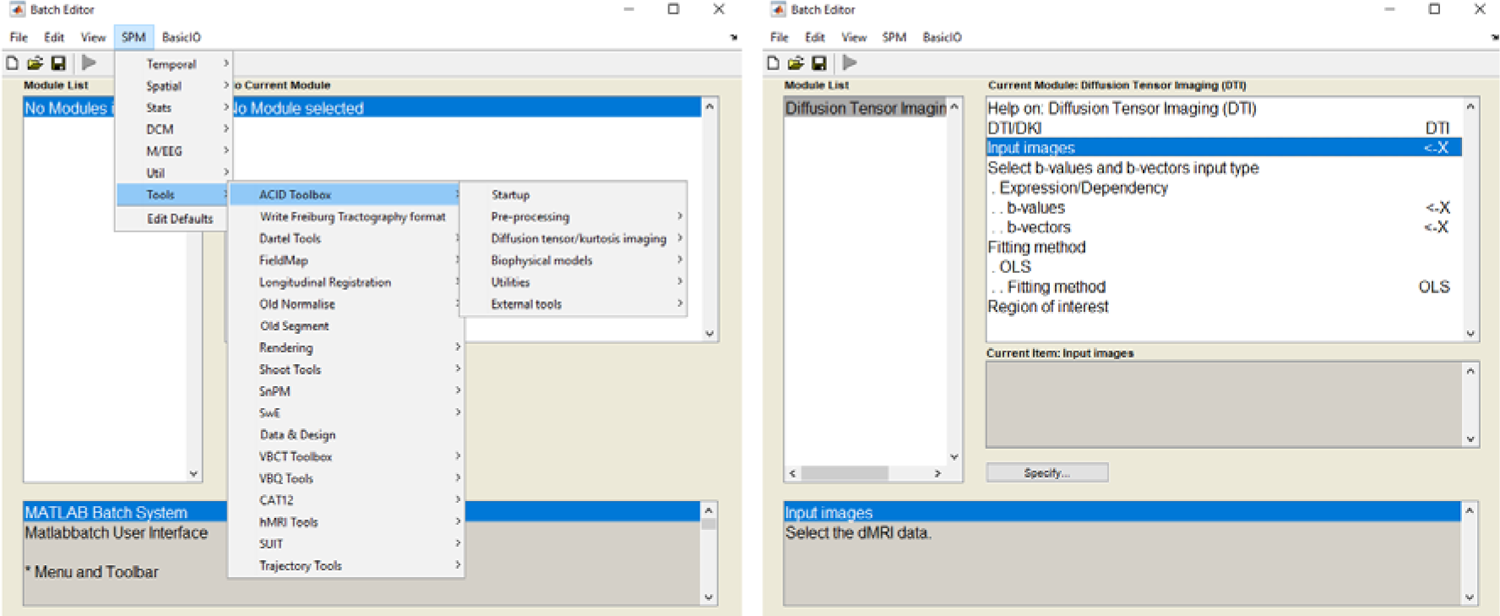
The left panel shows the location of ACID toolbox in the SPM Batch Editor after successful installation (SPM -> Tools). The toolbox is organized into six modules, each of which may be further divided into submodules. The right panel provides an example of a submodule *{Diffusion Tensor Imaging* within the *Diffusion tensor/kurtosis imaging*module). Each (sub-) module requires at least one mandatory input, indicated by “X”, as well as several optional inputs and parameter settings, which can be adjusted for customization. Recommended settings for typical in vivo brain, in vivo spinal cord, and ex vivo dMRI datasets are presented in Table 5.

### Appendix A.3. Startup

The ACID modules rely on a set of default settings, which were selected to yield reasonable results for typical dMRI data. However, adjustments may be necessary depending on the specific dataset (see Section 3.2 for recommendations). For convenience, the module’s graphical user interface (GUI) only presents the settings that are likely to be modified. Advanced users can access and modify all settings through the script config/local/acid_local_defaults.m. To use modified settings, the *Startup* module must be executed with the customized file provided as input; these settings will remain in effect even after restarting SPM or MATLAB until new settings are specified.

ACID requires all input images to be in NIfTI format (either NifTl-1 or NIfTI-2), with dMRI data required to be in 4D NifTI format. ACID also supports compressed NifTI images with the extension .nii.gz and outputs compressed images for compressed input and uncompressed images for uncompressed input. Users can convert from DICOM to NifTI format using SPM’s DICOM Import function, which can also export metadata into JSON files if the “Export metadata” option is enabled. To bring dMRI data into the required format, the *Startup* module can be utilized to (i) convert a set of 3D NifTI files into a single 4D NifTI file, (ii) generate corresponding bval/bvec files from the JSON files (if not already available), (iii) create an additional metadata file containing the most commonly reported subject and acquisition parameters (such as TE and TR) to provide a concise overview of the dataset, and (iv) set an output directory alternative to the default one. The output from *Startup* can be automatically passed to subsequent processing steps through dependencies.

## Appendix B. Details on ECMOCO

ECMOCO consists of four steps (Fig. BI):

1. The type of registration *{slice-wise* or *volume-wise)* and the degrees of freedom (DOF) for the affine transformation are specified by the user.
2. Shell-specific target volumes are generated, and transformation parameters are obtained between all non-diffusion-weighted (bO) volumes and their corresponding target. The parameter iteration for a given bO volume can be initialized by the transformation parameters of the preceding bO volume *(initialized registration,* see details below). Only the DOF associated with rigid-body transformation are estimated for bO volumes, as eddy currents are expected to be negligible in bO volumes due to the absence of diffusion-sensitizing gradients.
3. Transformation parameters are obtained between all diffusion-weighted (DW) volumes and their corresponding target. The parameter iteration for a given DW volume can be initialized by the interpolated transformation parameters from the bO volumes *(initializedregistration,* see details below).
4. The obtained transformation parameters are applied to reslice all volumes.

In addition to *slice-wise* registration, introduced in Section 2.2.1 and demonstrated in Fig. B2, ACID incorporates two additional recent features: *initialized registration* and *exclusion mode. Initialized registration* is based on the observation that transformation parameters obtained from high-SNR bO volumes tend to be more accurate than those obtained from low-SNR DW volumes. With *initialized registration,* the parameter iteration for each bO volume starts with the transformation parameters obtained from the preceding bO volume. Once all the bO volumes have been registered, their rigid-body transformation parameters are interpolated to the positions of the DW volumes situated between the bO volumes. Subsequently, the parameter iteration for each DW starts with these interpolated values. If interpolation is not feasible (e.g., the DW volume is situated before the first or after the last bO volume), the parameter iteration starts with the parameters obtained from the nearest bO volume. This approach is particularly useful for correcting slow spatial drifts across volumes.

**Fig. B1.**
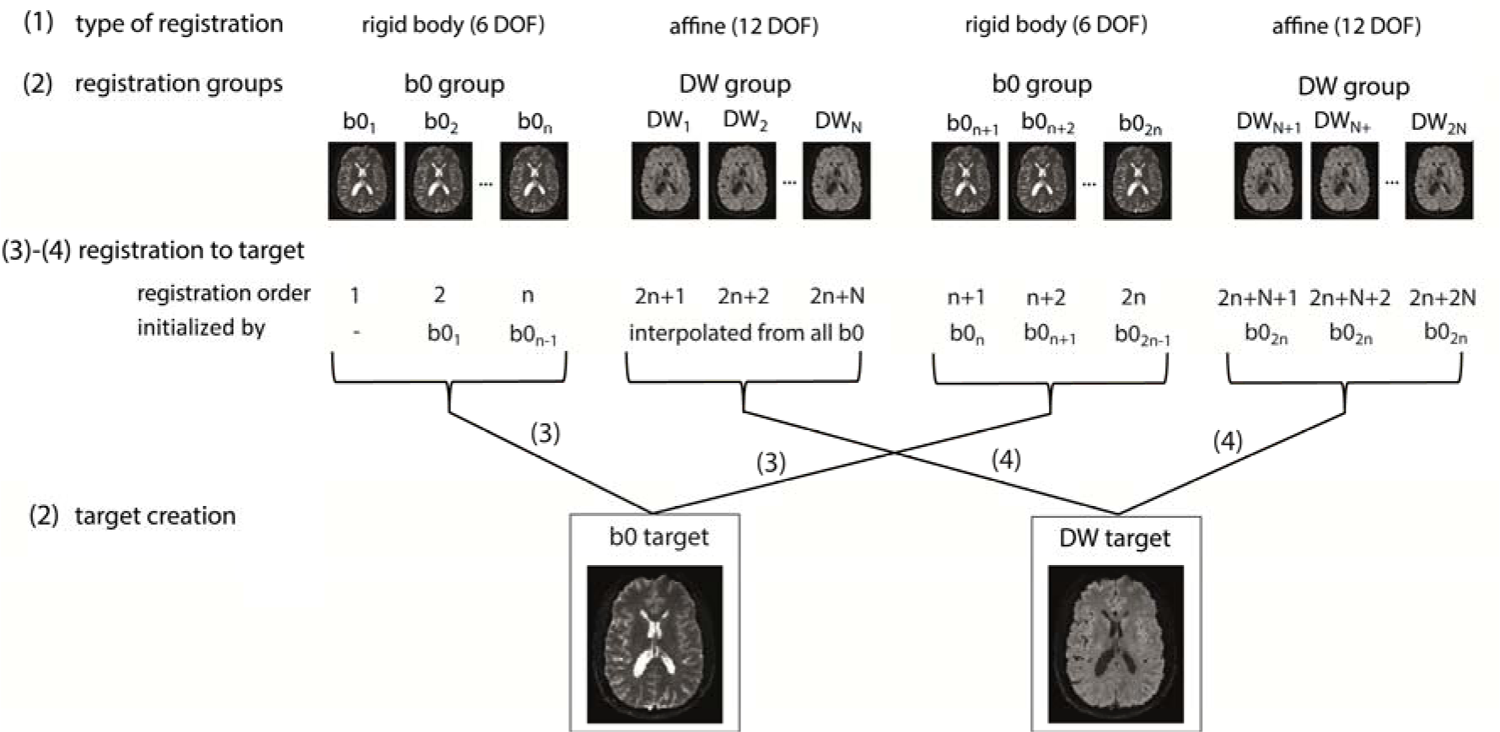
Registration scheme for an example dMRI dataset, which consists of two sets of non-diffusion-weighted (bO) volumes (volumes each) and two sets of diffusion-weighted (DW) volumes (volumes each) interspersed with each other. The bO and DW volumes form separate registration groups and are registered to their corresponding target volumes. First, the bO volumes are registered using the rigid-body components of the specified degrees of freedom (DOF), followed by the registration of the DW volumes using all specified DOF. The parameter iteration for a given bO or DW can be initialized using previously obtained transformation parameters *(initialized registration)*.

The *exclusion mode* is designed to address volumes with very low SNRs, which can make obtaining reliable transformation parameters difficult. Volumes that are considered not feasible for registration can be identified through visual inspection, e.g., using the *DWI series browser* utility function, and can be input into ECMOCO. For these volumes, the rigid-body transformation parameters from the preceding non-excluded volume are applied instead.

**Fig. B2.**
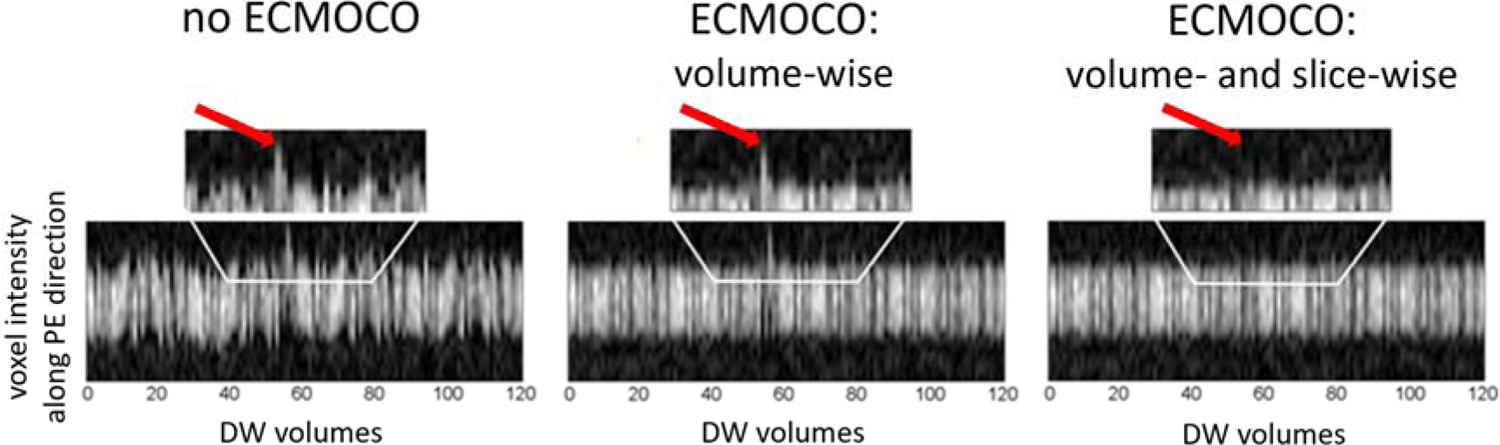
Qualitative comparison of different motion correction techniques including no correction, volume-wise ECMOCO, and the combination of volume- and slice-wise ECMOCO. The plots show the concatenation of ID cross-sections along the phase-encoding (PE) direction (anterior-posterior), extracted at fixed x- and z-coordinates from each of the 120 diffusion-weighted (DW) volumes in an in vivo spinal cord dMRI dataset. Additionally, zoomed-in views of a subset of DW volumes are provided to facilitate the assessment of improvements by ECMOCO. Substantial motion along the y-direction was initially observed, which was notably reduced after applying ECMOCO. Importantly, volume-wise ECMOCO did not entirely correct for spatial misalignments in all volumes (an example of failed correction is indicated by the red arrow). Conversely, the combination of volume- and slice-wise ECMOCO effectively corrected spatial misalignments in all DW volumes.

## Appendix C. Regions for noise estimation

For optimal denoising (msPOAS, Section 2.2.2) and Rician bias correction (Section 2.2.3), it is crucial to accurately estimate the image noise within the appropriate region of interest. Noise measurements taken from regions outside the body are often suboptimal due to the lower parallelization factor (g-factor) at the edge compared to the center of the field of view. Instead, we recommend estimating the noise by considering two distinct scenarios, employing the *repeated measures* method in each case (see *Noise estimation* in Table 2). In datasets affected by (temporally varying) physiological artifacts, such as in in vivo brain and spinal cord datasets, we recommend estimating the noise across images with high b-values and within regions where the signal reaches the noise plateau (i.e., within cerebrospinal fluid compartments). For automatic ventricle segmentation within the brain, ACID provides an example segmentation batch located at `ACID_TPM/acid-ventricles-batch.m,` which utilizes the *spm_segment* function. In datasets unaffected by physiological artifacts, such as in ex vivo dMRI, we recommend estimating the noise across non-diffusion-weighted (bO) images within either the entire specimen or a specific part. The latter recommendation, however, requires repeated measurements of bO images. Example noise regions are shown in Fig. Cl.

**Fig. C1.**
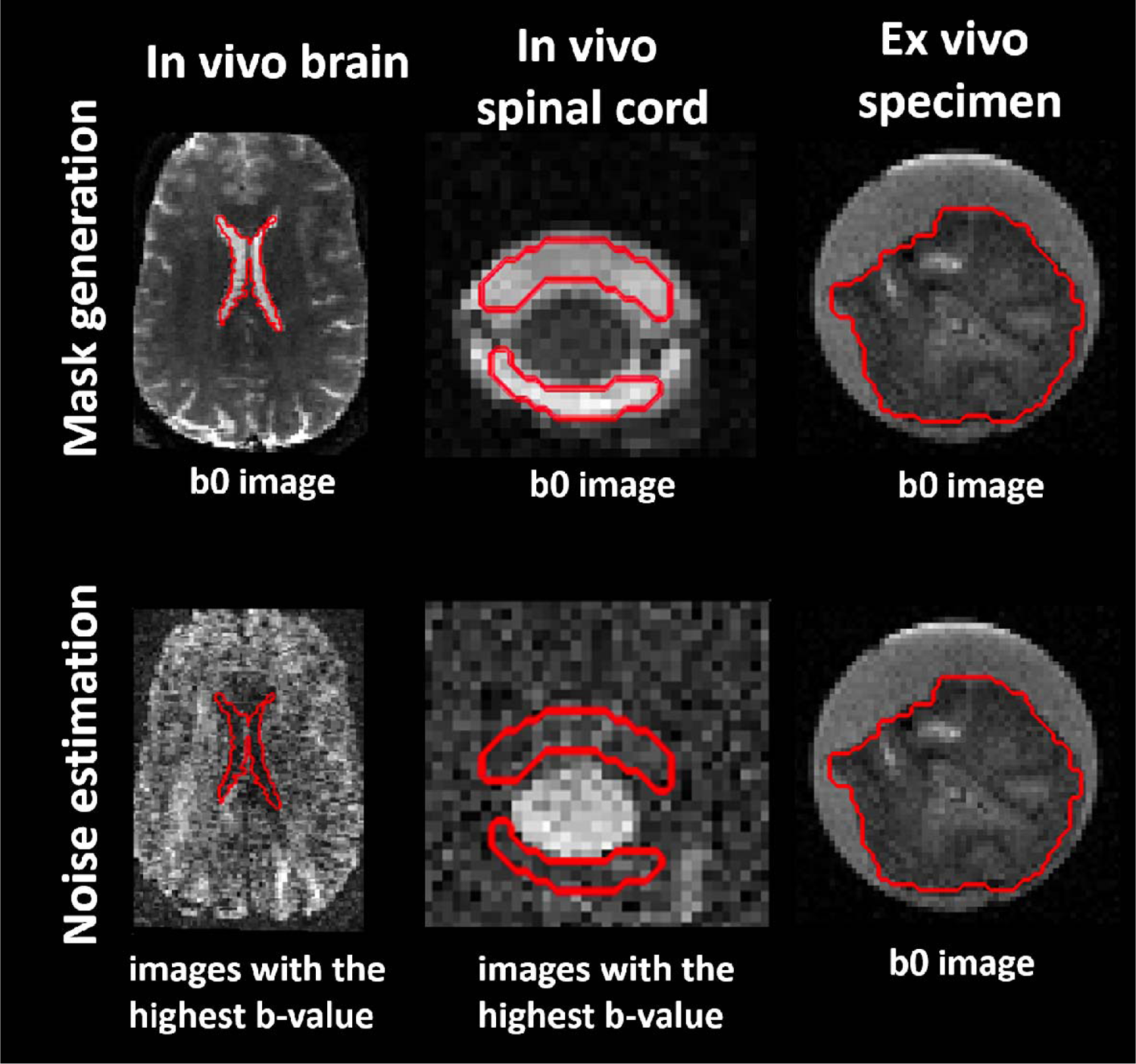
Definition of noise regions of interest (ROI) for the *repeated measures* noise estimation method (see *Noise estimation* in Table 2). Binary noise ROIs are outlined in red. For in vivo brain and spinal cord dMRI, we recommend creating a noise ROI within the cerebrospinal fluid (CSF), such as the lateral ventricles in the brain and the subarachnoid space in the spinal cord, on the bO images. Subsequently, we recommend estimating the noise on the images with the highest b-value (ideally above 1500 s/mm^2^) within the CSF mask. For ex vivo dMRI, the noise ROI is recommended to encompass the specimen itself, but noise estimation should be applied only on the bO images. Since ex vivo dMRI is not affected by physiological artifacts, signal variations across the bO images are considered noise.

## Appendix D. Recommendations for adaptive denoising (msPOAS)

If the overall noise reduction is insufficient, *kstar* can be increased at the cost of longer computation time (Tabelow et al., 2015). It is important to note that msPOAS assumes a single global value of *sigma,* which may not always hold. If *sigma* is correctly estimated, the default *lambda* value will ensure optimal adaptation. Incorrect estimation of *sigma* can be compensated by the choice of *lambda,* which makes msPOAS robust against misspecification of *sigma* (Becker et al., 2014). We recommend determining *kappa* automatically based on the number of diffusion directions (Tabelow et al., 2015). However, manual adjustment of *kappa* may be necessary in cases where the SNR is low (e.g., for spinal cord dMRI) or if the dataset has more images with high b-values than with low b-values. The effective number of coils *(ncoils)* is 1 when using SENSEI reconstructions (Polzehl & Tabelow, 2016; Sotiropoulos et al., 2013), but the correct value is more difficult to determine when using multiple receiver channels (Aja-Fernandez et al., 2014). It is important to use the same *ncoils* for the estimation of *sigma* and in msPOAS to ensure the same number of degrees of freedom.

## Appendix E. Model fitting methods implemented in ACID

### Appendix E.1. Ordinary Least Squares

Tensor fitting involves solving the linear regression problem ***y=Bα+ε***, where ***y*** contains the logarithmic signals, ***B*** (b-matrix) contains the gradient directions and strengths, ***α*** contains the elements of the diffusion tensor, and ε contains the model-fit errors (the difference between the actual and fitted signal). The ordinary least squares (OLS) approach employs the estimator function ρ(ε_i_)=ε i^2^, where ε_i_ represents the model-fit error of acquisition *i.* The solution is obtained by minimizing *Σ_i_ ε_i_^2^*, yielding ***α_ols_ = (B ^T^B)^-1^B^T^y*.**

**Fig. E1.**
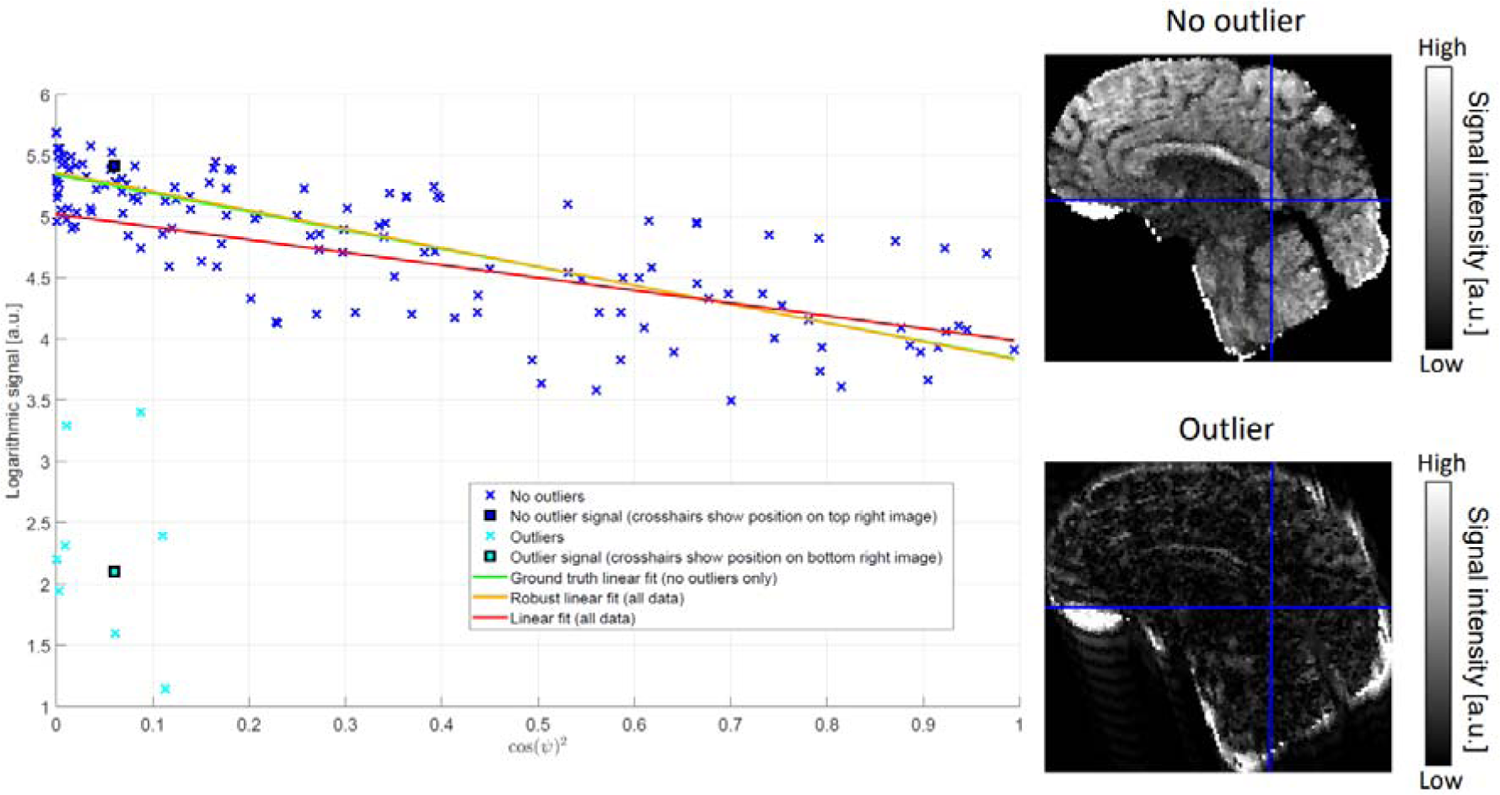
Schematic illustration of how robust fitting downweighs outliers in the model fit. The scatter plot shows the logarithm of diffusion-weighted voxel intensities against the squared cosine of the angle between the diffusion gradient direction (bvec) and the direction of the first eigenvector in a corpus callosum voxel (see blue crosshairs for location). Blue crosses in the scatter plot indicate data points not affected by artifacts (“No outliers”), while cyan crosses indicate data points affected by strong artifacts (“Outliers”). Outliers were generated by removing the center of the k-space of the original image to illustrate the effect of strong motion artifacts. Two example images corresponding to a non-artifactual (“No outlier”, top image) and an artifactual data point (“Outlier”, bottom image) are shown on the right. During the model fit, a linear curve is fitted to the logarithmic voxel intensities. The presence of outlier data points leads to a biased model fit (red line) and consequently biased tensor estimates when using ordinary least squares (OLS) model fitting. In contrast, robust fitting downweighs the influence of outliers, leading to a more accurate model fit (orange line) which is closer to the ground truth (green line) obtained by an OLS fit to the non-artifactual data points (blue crosses) only.

### Appendix E.2. Weighted Least Squares

The weighted least squares (WLS) approach addresses the heteroscedasticity of the logarithmic data by assigning individual weights to each image in the form of *ω_i_ =S_i_/σ_i_* where Ŝ_i_ represents the unknown true signal (without noise) and σ_i_ is the background noise for acquisition *i.* The estimator function now becomes ρ(ε_i_)=|ω_i_ε_i_~^2^, yielding the solution ***α_wls_ =$V ^T^B^T^WB)^^1^W^T^B^T^Wy,*** with ***W*** being the diagonal matrix of «j. Note that OLS is a special case of WLS, where *ω_i_,* =1 for all *i.* A practical consideration in obtaining ***a_wls_*** is related to estimating S_ε_. One approach is to use the measured noisy signal *S,* as an estimate of S_ε_. Another approach is to start with the OLS solution and use the fitted signal as an estimate of which was shown to be more accurate (Veraart et al., 2013b).

### Appendix E.3. Robust fitting

The concept behind robust fitting is to assign lower weights to data points with higher model-fit errors during the fitting process (Mangin et al., 2002). The robust fitting method implemented in ACID is based on the “Patching ArTefacts from Cardiac and Head motion” (PATCH) technique introduced by Zwiers, 2010. While the form of the estimator function is similar to that of WLS, PATCH factorizes the weighting function into three components as *=a>* where each component is designed to address different types of artifacts: cu_ε1_ and *a>_i2_* account for regional and slice-wise artifacts, respectively, while *u>_i3_* is identical to the weight term in WLS. w_ε1_ and *a>_i2_* are exponentially decaying functions of 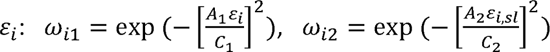, where 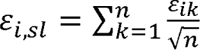 is the slice-average model-fit error, with *n* being the number of voxels within the slice. *A_1_* and Л_2_ ^are^ model parameters, by default set to 0.3 and 0.1, respectively, with higher values resulting in a faster exponential decay. C_t_ and C_2_ are estimates of the standard deviation of ε_ε_ and ε_εsε_, respectively, in the absence of outliers, and are computed as C_t_ = 1.4826 ▪ median(|ε_ε_|) and C_2_ = 1.4826 ▪ median(|ε_εsε_|) (Hampel, 1974; Rousseeuw & Croux, 1993). Note that accurate estimation of C_x_ and C_2_ is crucial for effectively downweighting outliers. This holds true as long as outliers are sparsely distributed and the median of the model-fit errors remains unaffected. However, a frequent occurrence of outliers can increase *C,* leading to a less effective downweighing of outliers.

While OLS and WLS independently fit the tensor in each voxel, PATCH makes use of the observation that physiological noise represents a structured, spatially correlated noise. To accommodate the anticipated smoothness of C_t_, the median operator is spatially smoothed using a 2D Gaussian kernel before computing C_x_ (Zwiers, 2010).

As a modification to PATCH, the robust fitting method incorporates Tikhonov regularization to handle ill-conditioned weighting matrices resulting from a high occurrence of outliers. This leads to the solution ***а_л_= [W^T^B^T^WB + ЛB^T^B]^-1^W^T^B^T^W^y^,*** where *W* represents the diagonal matrix of factorized weights, and A is the Tikhonov regularization factor. Notably, in the two extreme cases, the Tikhonov solution either becomes **α_wols_** (albeit with a different ***W)*** (A=0) or converges to **α_ols_** (λ→∞). The above equation cannot be solved readily, as ***W*** is a function of ε, which is only available after obtaining the solution. This is addressed by using an iteratively re-weighted least squares (IRLS) algorithm. In the first iteration, ω_i_ is set to 1 for all **t** to obtain the OLS solution **α_ols_** and the initial ε. In the second iteration, an updated ***W*** is computed based on the initial ε, which is then used to compute ***а_л_.*** In each further iteration, εfrom the preceding iteration is used to update ***W,*** which is in turn used to compute the updated ***а_л_.*** This iterative process is repeated until convergence or until the predefined number of iterations is exceeded.

### Appendix E.4. Non-linear least squares

The non-linear least squares (NLLS) method solves the optimization problem _, where represents the signal model (DTI or DKI), the model parameters (elements of the diffusion and/or kurtosis tensors), and the measured signal intensities for a specific diffusion weighting and diffusion gradient direction. The NLLS optimization problem is solved using a Gauss-Newton algorithm.

## Appendix F. Effect of Rician bias correction on biophysical parameter estimates

Here, we demonstrate the influence of Rician bias correction on the estimation of Watson concentration parameter (κ) and axonal water fraction (AWF) (Fig. FI). These biophysical parameters were estimated on the fully processed dataset using either the NODDI-DTI model applied on a single (lower b-value) shell or the WMTI-Watson model applied on two shells. For an in-depth analysis of the impact of Rician bias correction on DKI and axisymmetric DKI, refer to Oeschger et al., 2023a.

**Fig. F1.**
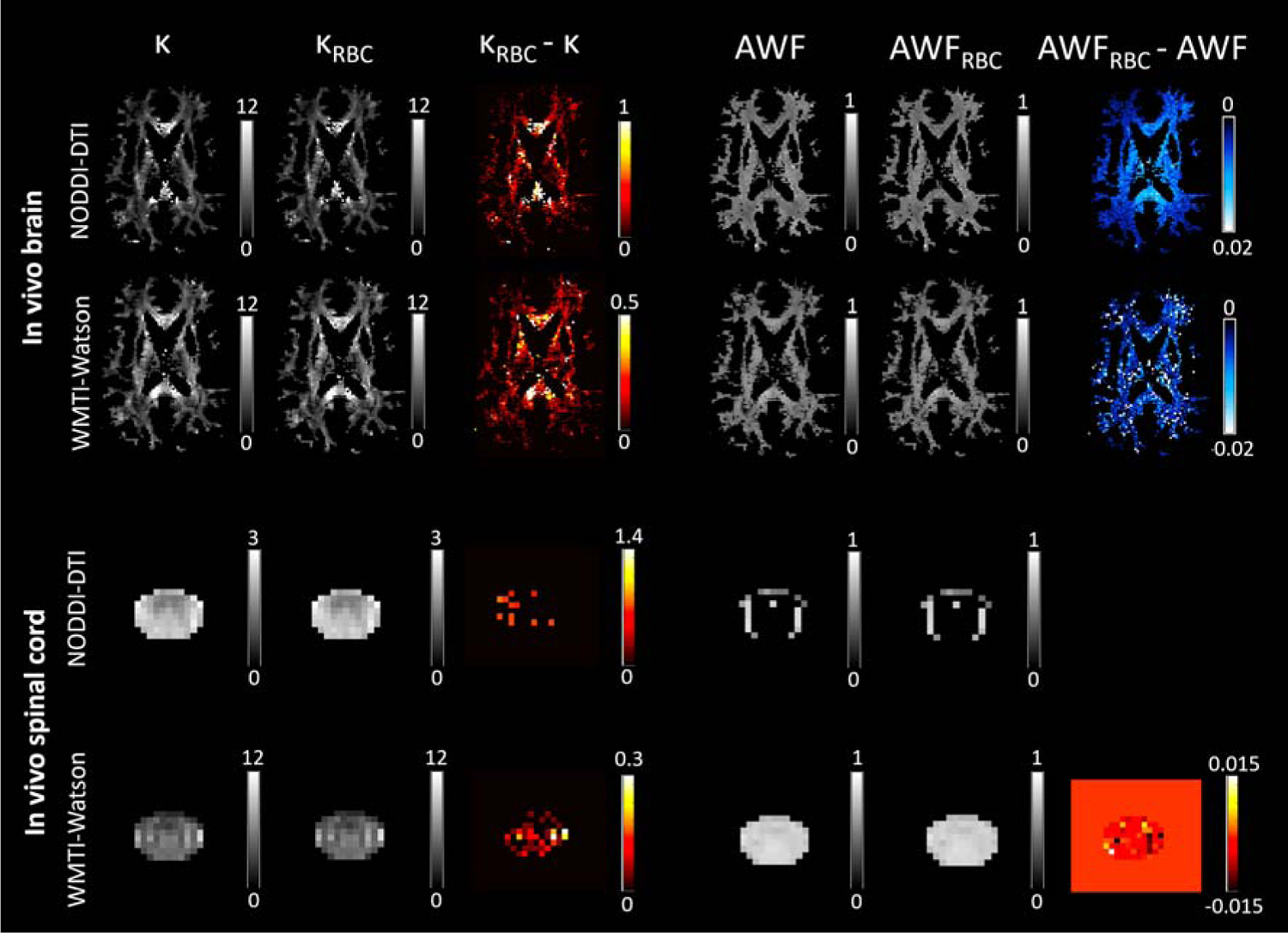
The impact of Rician bias correction (RBC) on maps of biophysical parameter estimates, derived from the NODDI-DTI and WMTI-Watson models, including Watson concentration parameter () and axonal water fraction (AWF), in an in vivo brain and spinal cord dataset (refer to Table 4 for details on the datasets). Being derived from white matter biophysical models, the parameter maps were masked for the white matter in the brain dataset. For the spinal cord and ex vivo specimen, we refrained from masking due to the difficulty of obtaining an accurate white matter mask. These maps were computed both without (left column) and with (middle column) RBC; their voxel-wise difference, referred to as the Rician bias, is shown in the right column. RBC slightly decreased the mean of the kurtosis tensor in both the brain and spinal cord, which resulted in an increase in. The estimation of AWF using the NODDI-DTI model was not feasible, as the mean diffusivity (MD) values derived from DTI fell below the range where the NODDI-DTI model provides a valid representation (refer to Equation (4) in Edwards et al., 2017). This discrepancy could be attributed to either the underestimation of **МГ)** rinp tn kurtnqiq hia«; (Fio nr thp invalidity nf fixpri rnmnartmpntal riiffiiqivitip<; in thp NODDI-DTI mnripl

## Appendix G. Evaluating denoising methods

Several denoising methods have been developed, including the Multi-Shell Position-Orientation Adaptive Smoothing (msPOAS, Section 2.2.2) (Becker et al., 2014), as well as methods based on local principal component analysis (LPCA) (Manjon et al., 2013) and Marchenko-Pastur principal component analysis (MP-PCA) (Veraart et al., 2016). Here, we evaluated these three denoising methods using a simulated dMRI dataset of the human brain. Specifically, we fitted the kurtosis model to an in vivo brain dMRI dataset (refer to Table 4 for details on the dataset) and considered the fitted dMRI signal as the “noise-free” ground truth. Then, we added varying amounts of noise to the ground truth, drawn from a circularly-symmetric complex normal distribution *CN(0,* c^2^) with 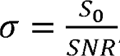 using the same set of SNR values (SNR=5, 15, 30, 39, 52, 100) as in our previous study (Oeschger et al., 2023b). The code for the simulation is available online^16^. For each SNR, the kurtosis model was fitted to the noisy magnitude dMRI data, both before (No denoising) and after denoising (msPOAS, LPCA, MP-PCA), using the non-linear least squares (NLLS) algorithm implemented in ACID. Slices of axial diffusivity (AD), radial diffusivity (RD), mean kurtosis tensor (MW), axial kurtosis tensor (AW), and radial kurtosis tensor (RW) maps obtained from the dMRI data with the lowest SNR (SNR=5) are shown in Fig. Gl. The kurtosis model was also fitted to the noise-free dMRI data for comparison (Ground truth, Fig. Gl). Deviations from the ground truth were quantified by computing the normalized root-mean-square error (NRMSE) between the obtained DKI metrics and the ground truth across white matter voxels for one noise realization (Fig. G2). The white matter mask applied is overlaid on the ground truth DKI metric maps in Fig. Gl.

In general, denoising methods proved beneficial in reducing NRMSE from the ground truth compared to the “no denoising” scenario in the low-SNR domain, although not consistently across all DKI metrics. Specifically, denoising reduced NRMSE for RD and RW below an SNR of 15, and for AW below an SNR of 30. However, it did not reduce NRMSE for AD, and the trend was not clear for MW. At higher SNRs (above 30-40), denoising increased NRMSE for all DKI metrics compared to the non-denoised data, except for the MP-PCA denoising method, which yielded results comparable to the non-denoised scenario. The relative difference between the maps generated using denoising and those generated without denoising is shown in Fig. G3. These results suggest that denoising (using any of the three methods) is beneficial for increasing the similarity to ground truth DKI metrics only in the low-SNR domain. In the high-SNR domain, denoising either does not lead to further improvements (MP-PCA) or even introduces additional errors (msPOAS and LPCA).

**Fig. G1.**
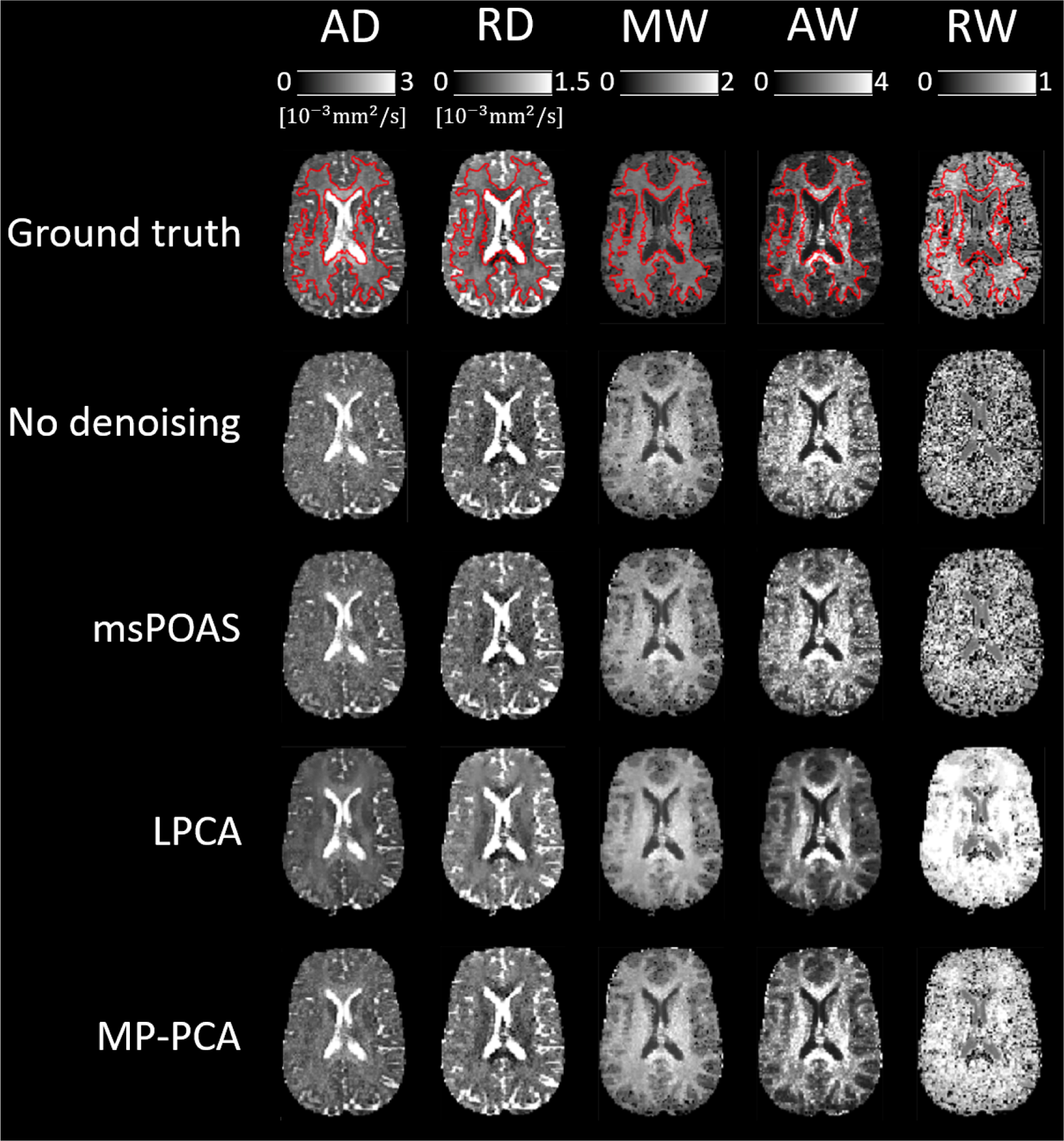
Qualitative illustration of the effect of denoising on maps derived from diffusion kurtosis imaging (DKI). Shown are maps of axial diffusivity (AD), radial diffusivity (AR), mean kurtosis tensor (MW), axial kurtosis tensor (AW), and radial kurtosis tensor (MW). The maps were obtained by fitting the kurtosis model to simulated noisy dMRI data (signal + noise) with a signal-to-noise ratio (SNR) of 5, both before (No denoising) and after employing different denoising methods (msPOAS, LPCA, MP-PCA). The DKI metric maps obtained from the simulated noise-free dMRI data (signal only) are also shown for comparison (Ground truth). The white matter mask used for calculating the normalized root-mean-square error (NRMSE) between the obtained DKI metrics and the ground truth is overlaid as a red segmentation line on the Ground truth maps.

**Fig. G2.**
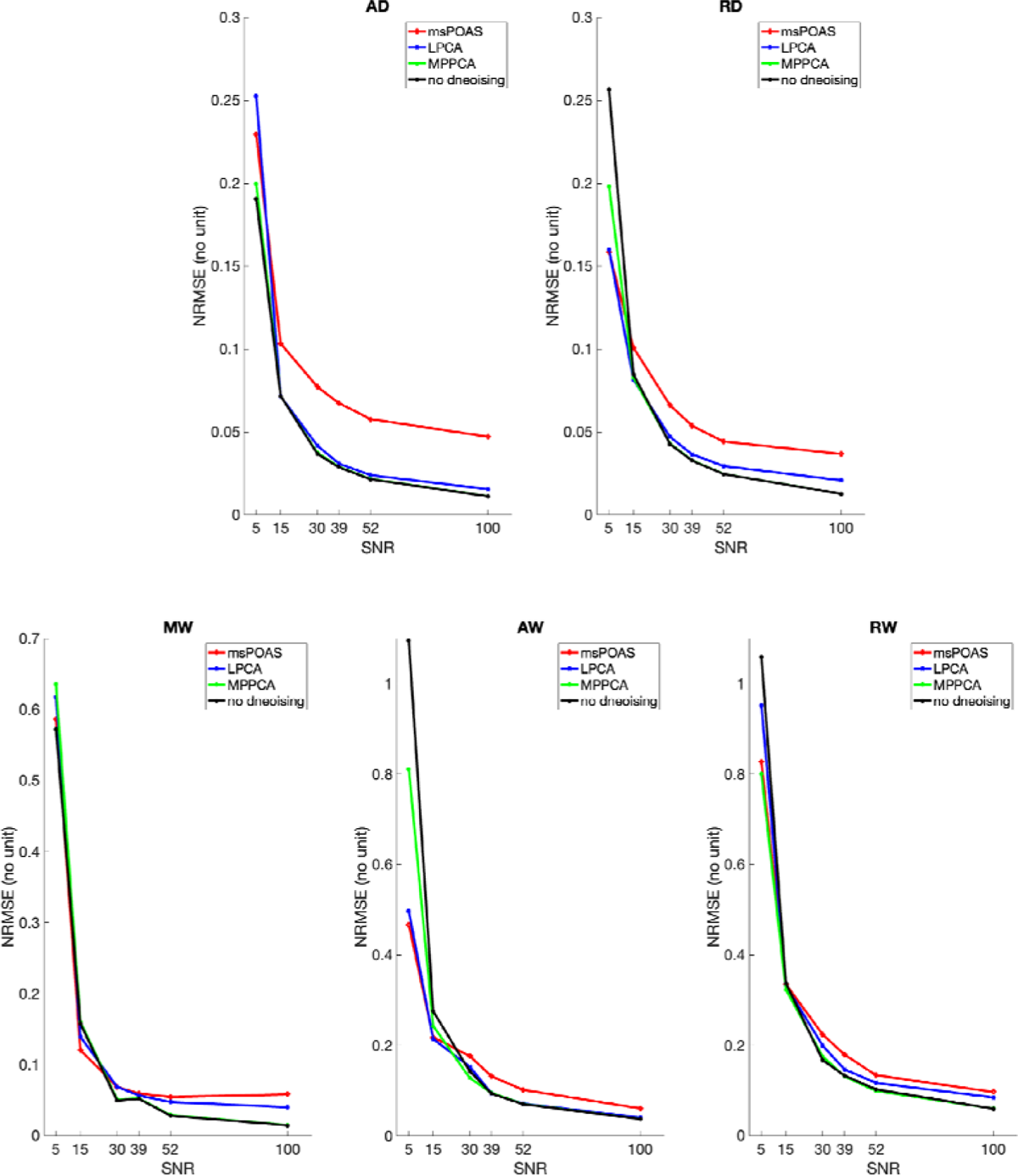
Quantitative illustration of the effect of denoising on maps derived from diffusion kurtosis imaging (DKI) (one noise realization). The plots show the normalized root-mean-square error (NRMSE) between (i) DKI metrics obtained from simulated noisy dMRI data (signal + noise) with varying signal-to-noise ratios (SNR), both before (no denoising) and after employing different denoising methods (msPOAS, LPCA, MP-PCA), and (ii) DKI metrics obtained from noise-free dMRI data (signal only). NRMSE was computed across white matter voxels (see Fig. G1 for the white matter mask) for the following DKI metrics: axial diffusivity (AD), radial diffusivity (RD), mean kurtosis tensor (MW), axial kurtosis tensor (AW), and radial kurtosis tensor (RW). Denoising methods reduced NRMSE from the ground truth compared to the “no denoising” scenario only in the low-SNR domain, although not consistently for all DKI metrics. At high SNRs (above 30-40), denoising increased NRMSE for all DKI metrics, except for the MP-PCA method, which yielded results comparable to the “no denoising” scenario.

**Fig. G3.**
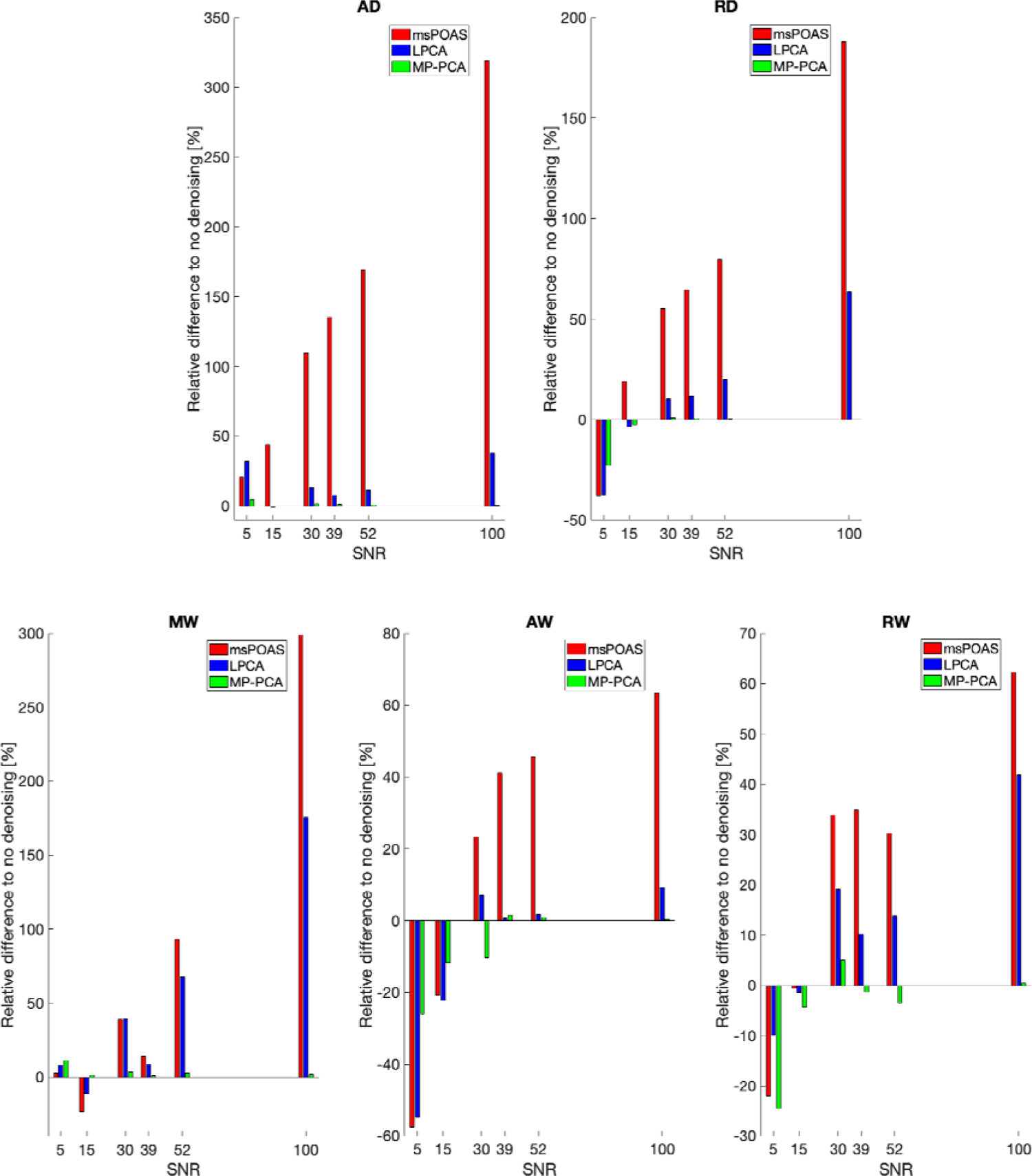
Quantitative illustration of the effect of denoising on maps derived from diffusion kurtosis imaging (DKI). The plots show the relative difference in DKI metrics obtained from simulated noisy dMRI data with varying signal-to-noise ratios (SNR) after employing different denoising methods (msPOAS, LPCA, MP-PCA) to those obtained without denoising (one noise realization). The relative difference was computed across white matter voxels (see Fig. Gl for the white matter mask) for the following DKI metrics: axial diffusivity (AD), radial diffusivity (RD), mean kurtosis tensor (MW), axial kurtosis tensor (AW), and radial kurtosis tensor (RW). Denoising introduced substantial improvements in the investigated DKI metrics only in the low-SNR domain, although not consistently across all DKI metrics. When using msPOAS and LPCA, the relative differences were greater compared to using MP-PCA, with msPOAS introducing the highest bias. At high SNRs (above 30-40), the relative difference to the “no denoising” scenario was negligible for MP-PCA.

## Supplementary material

**Fig. S1.**
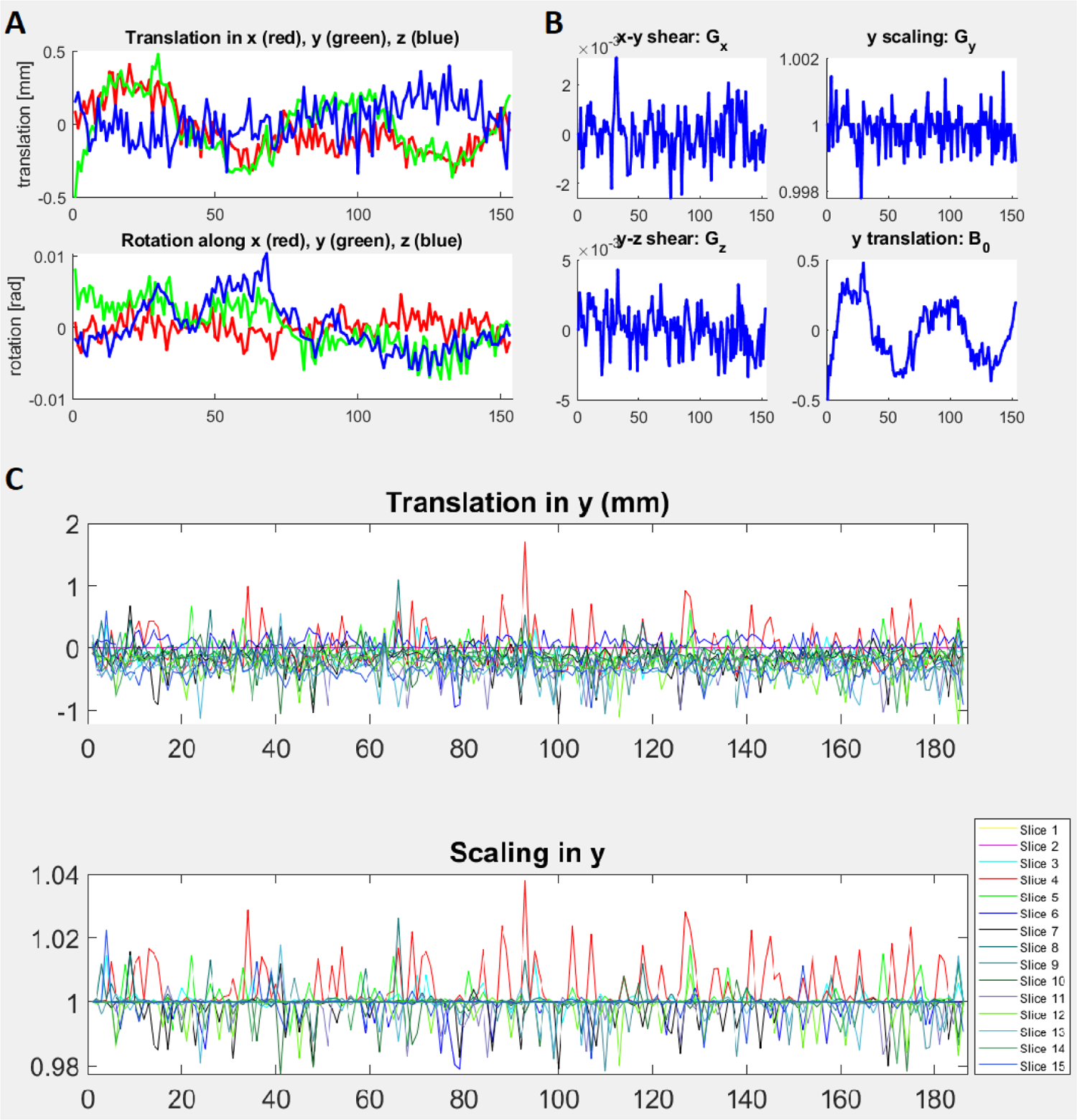
Diagnostic plots, optionally generated by ECMOCO, displaying the transformation parameters for all volumes (in the case of volume-wise registration) or slices (in the case of slice-wise registration). In volume-wise registration, demonstrated here with an in vivo brain dMRI dataset, two figures are created to plot the transformation parameters associated with motion (A) and eddy-current-related displacements (B). In slice-wise registration, shown here with an in vivo spinal cord dMRI dataset, a single figure is created to plot the transformation parameters with separate subfigures for each estimated degree of freedom (C). Excessive displacements in volumes/slices indicate either extreme movements, eddy-current artifacts, or a failed estimation of transformation parameters.

**Fig. S2.**
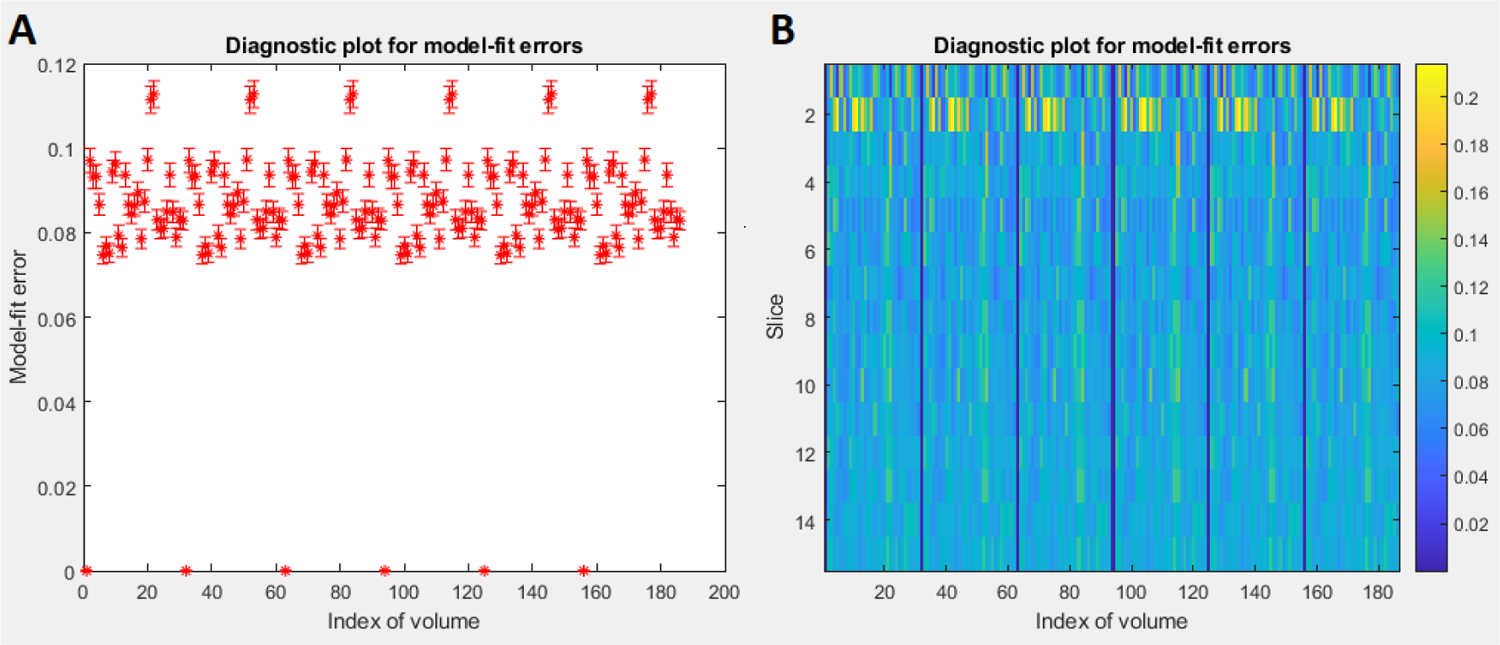
Diagnostic plots, optionally generated by the *Diffusion tensor/kurtosis imaging* module, displaying the average (logarithmic) model-fit error within the provided region of interest for each volume and slice, demonstrated here with an in vivo spinal cord dataset and a spinal cord mask. Volumes/slices with high model-fit error (outliers) indicate a high number of corrupted volumes (e.g., due to misregistration, physiological, or other artifacts) or an inadequate model for capturing the underlying complexity of diffusion. Here, periodically occurring pairs of volumes with high model-fit errors result from an inadequate model fit due to the low signal-to-noise ratio caused by the diffusion-sensitizing gradient aligned parallel to the spinal cord (A). Also, notice that the model-fit error is highest within slice 2, which could be attributed to the presence of physiological artifacts in that location. For an even more precise diagnosis of signal outliers, the voxel-wise root-mean-square of the model-fit error map (suffix: RMSE-LOG_map.nii) or the 4D model-fit error map (suffix: ERROR-LOG_map.nii) can be visually inspected to help identify individual outlier voxels or data points.

**Fig. S3.**
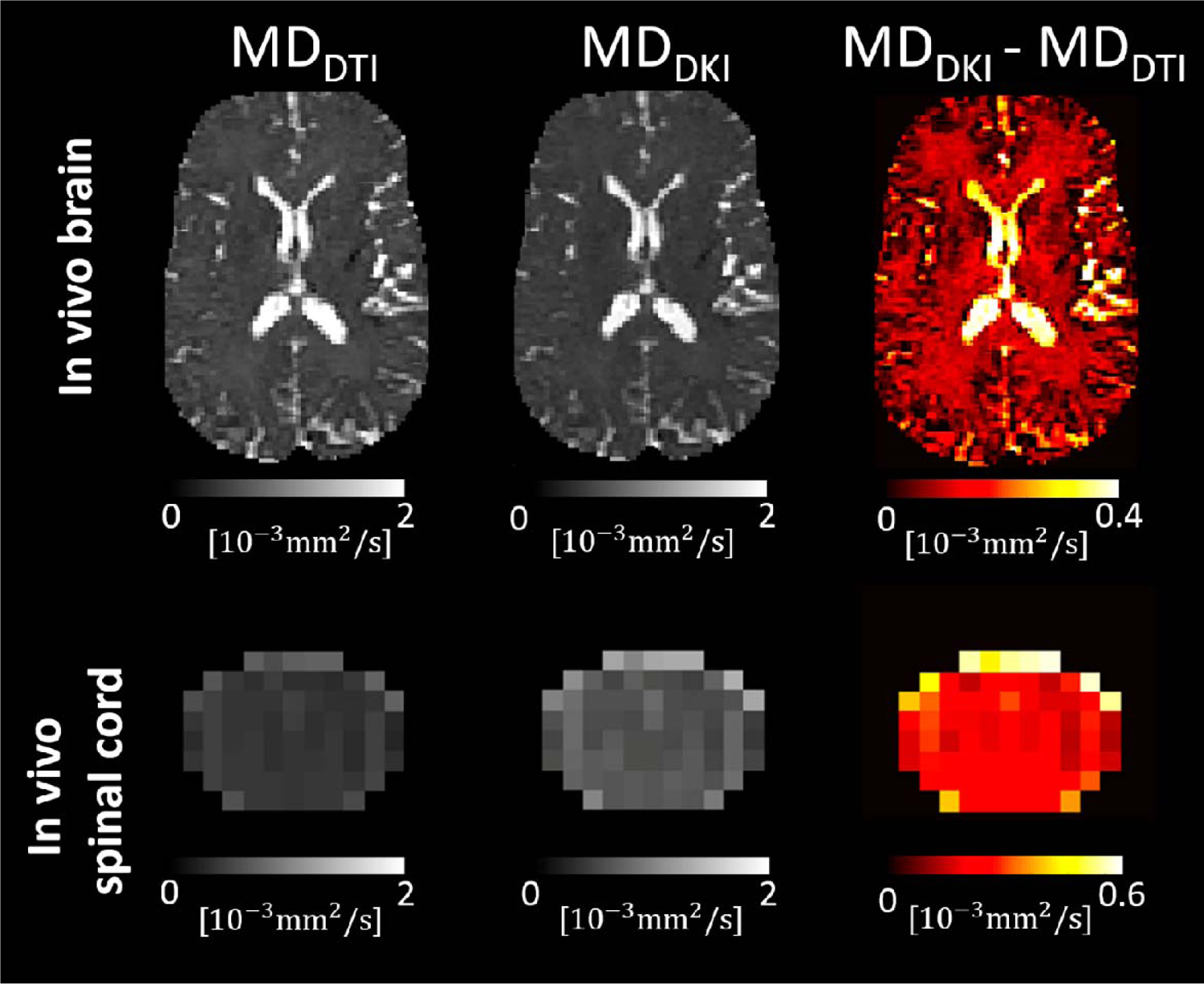
Kurtosis bias in the mean diffusivity (MD) maps in an in vivo brain and in vivo spinal cord dataset (refer to Table 4 for details on the datasets). This bias, shown in the right column, refers to the difference in the estimated diffusivity values when using the lower diffusion shells only (, tensor model, left column) or both the lower and higher diffusion shells (, kurtosis model, middle column). On average, the kurtosis bias was 12% and 54% within the brain white matter and the whole spinal cord, respectively.

**Fig. S4.**
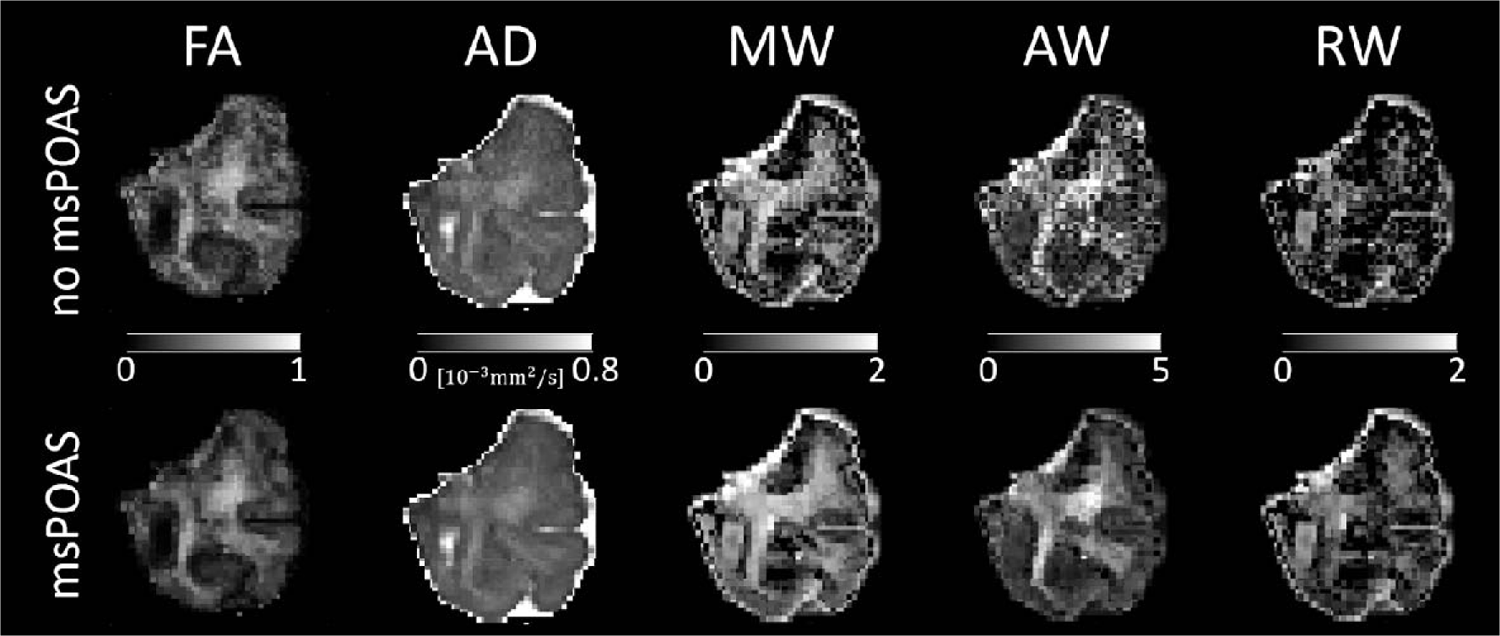
Comparison between maps of fractional anisotropy (FA), axial diffusivity (AD), mean kurtosis tensor (MW), axial kurtosis tensor (AW), and radial kurtosis tensor (RW) with and without applying adaptive denoising (msPOAS). The msPOAS-corrected maps appear less noisy while preserving tissue edges.

**Fig. S5.**
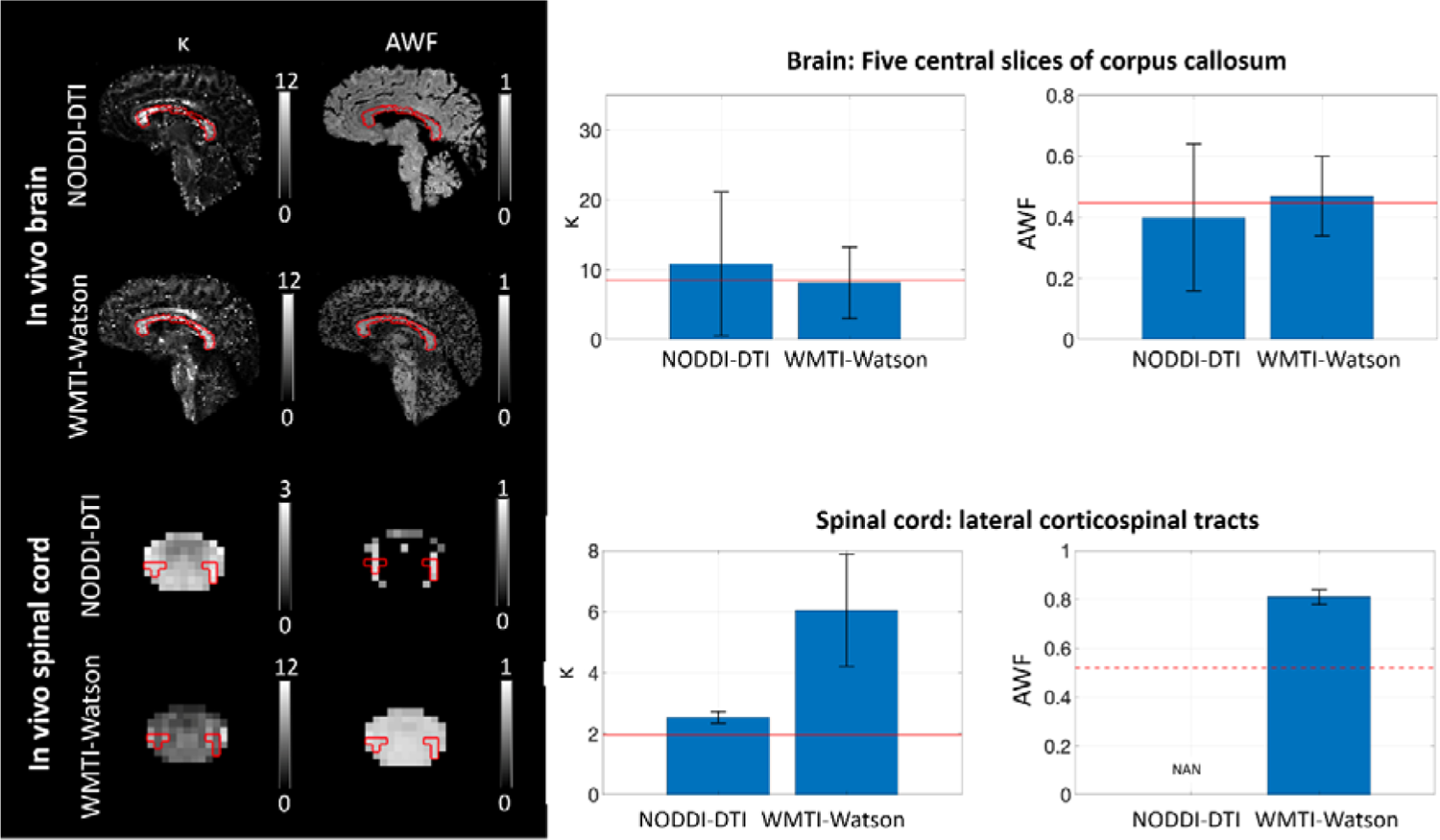
Bar plots displaying the Watson concentration parameter (κ) and axonal water fraction (AWF) within the five central slices of the corpus callosum and the lateral corticospinal tracts in the spinal cord (refer to Table 4 for details on the datasets). The corpus callosum was manually segmented, while the lateral corticospinal tracts were segmented using the PAM50 spinal cord white matter atlas. The ROI of an example slice is shown on the left side for each parameter. Red horizontal lines represent literature values obtained by histology, while the red dotted line represents a literature value obtained in the brain, given the absence of a corresponding value in the spinal cord. Orientation dispersion index values reported in the literature were converted to κ using Equation (1) in Mollink et al., 2017. Within the corpus callosum, к values were (mean ± std) 10.82 ± 10.31 and 8.14 ± 5.13 when derived from the NODDI-DTI (single shell) and WMTI-Watson model (two shells), respectively. These values fall within the range of literature values obtained post-mortem using polarized light imaging (Mollink et al., 2017). AWF values derived from NODDI-DTI (0.40 ± 0.24) and WMTI-Watson model (0.47 ± 0.13) were similar to literature values obtained using electron microscopy in a cynomolgus macaque (Stikov et al., 2015). Within the lateral corticospinal tracts, к values derived from NODDI-DTI were notably lower than those derived from WMTI-Watson (2.53 ± 0.19 vs. 6.04 ± 1.84) and were consistent with literature values obtained in a post-mortem specimen (Grussu et al., 2017). AWF values derived from the WMTI-Watson model in the spinal cord were substantially higher (0.81 ± 0.03) compared to a literature value obtained in the brain (red dotted line). The estimation of AWF was not feasible using the NODDI-DTI model, as DTI-derived mean diffusivity (MD) values fell below the range where the NODDI-DTI model provides a valid representation (refer to Equation (4) in Edwards et al., 2017). This discrepancy could be attributed to either the underestimation of MD due to kurtosis bias (Fig. S3) or the invalidity of fixed compartmental diffusivities in the NODDI-DTI model. These results indicate that WMTI-Watson yields more accurate estimation of κ and AWF for the brain, while NODDI-DTI yields a more accurate estimation of κ for the spinal cord. This could be a consequence of non-optimal b-values for kurtosis estimation in the spinal cord.

## Footnotes

[1] https://fsl.fmrib.ox.ac.uk/fsl/fslwiki/eddy

[2] https://fsl.fmrib.ox.ac.uk/fsl/fslwiki/topup

[3] https://mrtrix.readthedocs.io/en/dev/dwi_preprocessing/denoising.html

[4] https://sites.google.com/site/pierrickcoupe/softwares/denoising/dwi-denoising/dwi-denoising-software

[5] https://github.com/jan-martin-mri/koays-inversion

[6] https://mrtrix.readthedocs.io/en/dev/reference/commands/mrdegibbs.html

[7] https://www.fil.ion.ucl.ac.uk/spm/ext/

[8] http://www.emmanuelcaruyer.com/q-space-sampling.php

[9] https://fsl.fmrib.ox.ac.uk/fsl/fslwiki/eddy

[10] http://www.diffusiontools.org/sidebar/studies-using-acid.html

[11] http://www.fil.ion.ucl.ac.uk/spm/software/spml2/

[12] http://www.diffusiontools.com/

[13] https://bitbucket.org/siawoosh/acid-artefact-correction-in-diffusion-mri

[14] https://bitbucket.org/siawoosh/acid-artefact-correction-in-diffusion-mri/wiki/Home

[15] http://www.gnu.org/copyleft/

[16] https://github.com/quantitative-mri-and-in-vivo-histology/esmrmb2024

## References

Ades-Aron, B., Veraart, J., Kochunov, P., McGuire, S., Sherman, P., Kellner, E., Novikov, D. S., & Fieremans, E. (2018). Evaluation of the accuracy and precision of the diffusion parameter Estimation with Gibbs and NoisE removal pipeline. NeuroImage 183, 532–543. 10.1016/j.neuroimage.2018.07.066

Aja-Fernandez, S., Vegas-Sanchez-Ferrero, G., & Tristan-Vega, A. (2014). Noise estimation in parallel MRI: GRAPPA and SENSE. Magnetic Resonance Imaging 32(3), 281–290. 10.1016/j.mri.2013.12.001

Alexander, D. C., Dyrby, T. B., Nilsson, M., & Zhang, H. (2019). Imaging brain microstructure with diffusion MRI: practicality and applications. NMR in Biomedicine 32(*4*), e3841. 10.1002/nbm.3841

Andersson, J. L. R. (2008). Maximum a posteriori estimation of diffusion tensor parameters using a Rician noise model: Why, how and but. NeuroImage 42(*4*), 1340–1356. 10.1016/j.neuroimage.2008.05.053

Andersson, J. L. R., & Sotiropoulos, S. N. (2016). An integrated approach to correction for off-resonance effects and subject movement in diffusion MR imaging. NeuroImage 125, 1063­1078. 10.1016/j.neuroimage.2015.10.019

Andersson, M., Pizzolato, M., Kjer, H. M., Skodborg, K. F., Lundell, H., & Dyrby, T. B. (2022). Does powder averaging remove dispersion bias in diffusion MRI diameter estimates within real 3D axonal architectures? NeuroImage 248, 118718. 10.1016/j.neuroimage.2021.118718

André, E. D., Grinberg, F., Farrher, E., Maximov, I.I., Shah, N. J., Meyer, C., Jaspar, M., Muto, V., Phillips, C., & Balteau, E. (2014). Influence of noise correction on intra- and inter-subject variability of quantitative metrics in diffusion kurtosis imaging. PLoS ONE 9(4). 10.1371/journal.pone.0094531

Ashburner, J. (2007). A fast diffeomorphic image registration algorithm. NeuroImage 38(1), 95–113. 10.1016/j.neuroimage.2007.07.007

Ashburner, J., & Friston, K. J. (2005). Unified segmentation. NeuroImage 26(*3*), 839–851. 10.1016/j.neuroimage.2005.02.018

Ashburner, J., & Friston, K. J. (2011). Diffeomorphic registration using geodesic shooting and Gauss-Newton optimisation. NeuroImage 55(*3*), 954–967. 10.1016/j.neuroimage.2010.12.049

Barazany, D., Basser, P. J., & Assaf, Y. (2009). In vivo measurement of axon diameter distribution in the corpus callosum of rat brain. Brain 132(*5*), 1210–1220. 10.1093/brain/awp042

Barker, G. J. (2001). Diffusion-weighted imaging of the spinal cord and optic nerve. Journal of the Neurological Sciences, 186, 45–49. 10.1016/S0022-510X(01)00490-7

Basser, P. J., Mattiello, J., & LeBihan, D. (1994). Estimation of the effective self-diffusion tensor from the NMR spin echo. Journal of Magnetic Resonance, Series B, 103(*3*), 247–254. 10.1006/jmrb.1994.1037

Basser, P. J., & Pajevic, S. (2000). Statistical artifacts in diffusion tensor MRI (DT-MRI) caused by background noise. Magnetic Resonance in Medicine 44(*1*), 41–50. 10.1002/1522-2594(200007)44:1<41::AID-MRM8>3.0.CO;2-0

Becker, S. M. A., Tabelow, K., Mohammadi, S., Weiskopf, N., & Polzehl, J. (2014). Adaptive smoothing of multi-shell diffusion weighted magnetic resonance data by msPOAS. NeuroImage 95, 90­105. 10.1016/j.neuroimage.2014.03.053

Becker, S. M. A., Tabelow, K., Voss, H. U., Anwander, A., Heidemann, R. M., & Polzehl, J. (2012). Position-orientation adaptive smoothing of diffusion weighted magnetic resonance data (POAS). Medical Image Analysis, 16(6), 1142–1155. 10.1016/j.media.2012.05.007

Blaiotta, C., Freund, P., Cardoso, M. J., & Ashburner, J. (2017). Generative diffeomorphic atlas construction from brain and spinal cord MRI data. *ArXiv.* http://arxiv.org/abs/1707.01342

Büeler, S., Freund, P., Kessler, T. M., Liechti, М. D., & David, G. (2024). Improved inter-subject alignment of the lumbosacral cord for group-level in vivo gray and white matter assessments: A scan-rescan MRI study at 3T. PLOS ONE 19(*A*), e0301449. 10.1371/JOURNALPONE.0301449

Callaghan, P. T., Eccles, C. D., & Xia, Y. (1988). NMR microscopy of dynamic displacements: K-space and q-space imaging. Journal of Physics E: Scientific Instruments 21(*8*), 820–822. 10.1088/0022-3735/21/8/017

Caruyer, E., Lenglet, C., Sapiro, G., & Deriche, R. (2013). Design of multishell sampling schemes with uniform coverage in diffusion MRI. Magnetic Resonance in Medicine, 69(*6*), 1534–1540. 10.1002/mrm.24736

Chang, L. C., Jones, D. K., & Pierpaoli, C. (2005). RESTORE: Robust estimation of tensors by outlier rejection. Magnetic Resonance in Medicine 53(5), 1088–1095. 10.1002/mrm.20426

Chang, L. C., Walker, L, & Pierpaoli, C. (2012). Informed RESTORE: A method for robust estimation of diffusion tensor from low redundancy datasets in the presence of physiological noise artifacts. Magnetic Resonance in Medicine 68(*5*), 1654–1663. 10.1002/mrm.24173

Chun, S. Y., Li, K. C., Xuan, Y., Xun, M. J., & Qin, W. (2005). Diffusion tensor tractography in patients with cerebral tumors: A helpful technique for neurosurgical planning and postoperative assessment. European Journal of Radiology 56(*2*), 197–204. 10.1016/j.ejrad.2005.04.010

Clark, I. A., Callaghan, M. F., Weiskopf, N., Maguire, E. A., & Mohammadi, S. (2021). Reducing susceptibility distortion related image blurring in diffusion MRI EPI data. Frontiers in Neuroscience 15, 955. 10.3389/fnins.2021.706473

Coelho, S., Baete, S. H., Lemberskiy, G., Ades-Aron, B., Barrol, G., Veraart, J., Novikov, D. S., & Fieremans, E. (2022). Reproducibility of the Standard Model of diffusion in white matter on clinical MRI systems. NeuroImage 257, 119290. 10.1016/j.neuroimage.2022.119290

Cohen-Adad, J., Rossignol, S., & Hoge, R. (2009). Slice-by-slice motion correction in spinal cord fMRI: SliceCorr. Proceedings of the 17th Scientific Meeting, International Society for Magnetic Resonance in Medicine, Honolulu, USA, 3181.

Cohen, Y., Anaby, D., & Morozov, D. (2017). Diffusion MRI of the spinal cord: from structural studies to pathology. NMR in Biomedicine 30(3). 10.1002/nbm.3592

Constantinides, C. D., Atalar, E., & McVeigh, E. (1997). Signal-to-noise measurements in magnitude images from NMR phased arrays. Proceedings of the Annual International Conference of the IEEE Engineering in Medicine and Biology 1, 456–459. 10.1109/iembs.1997.754578

David, G., Freund, P., & Mohammadi, S. (2017). The efficiency of retrospective artifact correction methods in improving the statistical power of between-group differences in spinal cord DTI. NeuroImage 158, 296–307. 10.1016/j.neuroimage.2017.06.051

David, G., Fricke, B., Oeschger, Jan, M., Ruthotto, L., Fritz, Francisco, J., Ohana, O., Sauvigny, T., Freund, P., Tabelow, K., & Mohammadi, S. (2024). ACID: A Comprehensive Toolbox for Image Processing and Modeling of Brain, Spinal Cord, and Ex Vivo Diffusion MRI Data - Software. 10.20347/ACIDTBX

David, G., Pfyffer, D., Vallotton, K., Pfender, N., Thompson, A., Weiskopf, N., Mohammadi, S., Curt, A., & Freund, P. (2021). Longitudinal changes of spinal cord grey and white matter following spinal cord injury. *Journal of Neurology*, Neurosurgery and Psychiatry 92, 1222–1230. 10.1136/jnnp-2021-326337

David, G., Seif, M., Huber, E., Hupp, M., Rosner, J., Dietz, V., Weiskopf, N., Mohammadi, S., & Freund, P. (2019). In vivo evidence of remote neural degeneration in the lumbar enlargement after cervical injury. Neurology 92(12), E1367–E1377. 10.1212/WNL.0000000000007137

David, G., Vallotton, K., Hupp, M., Curt, A., Freund, P., & Seif, M. (2022). Extent of cord pathology in the lumbosacral enlargement in non-traumatic versus traumatic spinal cord injury. Journal of Neurotrauma 39(9-10), 639–650. 10.1089/neu.2021.0389

De Groote, S., Goudman, L, Peeters, R., Linderoth, B., Vanschuerbeek, P., Sunaert, S., De Jaeger, M., De Smedt, A., & Moens, M. (2020). Magnetic resonance imaging exploration of the human brain during 10 kHz spinal cord stimulation for failed back surgery syndrome: A resting state functional magnetic resonance imaging study. Neuromodulation, 23(1), 46–55. 10.1111/ner.12954

De Leener, B., Fonov, V. S., Collins, D. L, Callot, V., Stikov, N., & Cohen-Adad, J. (2018). PAM50: Unbiased multimodal template of the brainstem and spinal cord aligned with the ICBM152 space. NeuroImage 165, 170–179. 10.1016/j.neuroimage.2017.10.041

De Leener, B., Lévy, S., Dupont, S. M., Fonov, V. S., Stikov, N., Collins, D. L., Callot, V., & Cohen-Adad, J. (2017). SCT: Spinal Cord Toolbox, an open-source software for processing spinal cord MRI data. Neurolmage 145, 24–43. 10.1016/j.neuroimage.2016.10.009

Deppe, M., Krämer, J., Tenberge, J. G., Marinell, J., Schwindt, W., Deppe, K., Groppa, S., Wiendl, H., & Meuth, S. G. (2016). Early silent microstructural degeneration and atrophy of the thalamocortical network in multiple sclerosis. Human Brain Mapping 37(5), 1866–1879. 10.1002/hbm.23144

Deppe, M., Tabelow, K., Krämer, J., Tenberge, J. G., Schiffler, P., Bittner, S., Schwindt, W., Zipp, F., Wiendl, H., & Meuth, S. G. (2016). Evidence for early, non-lesional cerebellar damage in patients with multiple sclerosis: DTI measures correlate with disability, atrophy, and disease duration. Multiple Sclerosis 22(1), 73–84. 10.1177/1352458515579439

Dietrich, O., Raya, J. G., Reeder, S. B., Reiser, M. F., & Schoenberg, S. 0. (2007). Measurement of signal-to-noise ratios in MR images: Influence of multichannel coils, parallel imaging, and reconstruction filters. Journal of Magnetic Resonance Imaging 26(2), 375–385. 10.1002/jmri.20969

Dossi, D. E., Chaves, H., Heck, E. S., Murúa, S. R., Ventrice, F., Bakshi, R., Quintana, F. J., Correale, J., & Farez, M. F. (2018). Effects of systolic blood pressure on brain integrity in multiple sclerosis. Frontiers in Neurology 9, 487. 10.3389/fneur.2018.00487

Draganski, B., Ashburner, J., Hutton, C., Kherif, F., Frackowiak, R. S. J., Helms, G., & Weiskopf, N. (2011). Regional specificity of MRI contrast parameter changes in normal ageing revealed by voxel-based quantification (VBQ). NeuroImage, 55(4), 1423–1434. 10.1016/j.neuroimage.2011.01.052

Dubois, J., Dehaene-Lambertz, G., Kulikova, S., Poupon, C., Hüppi, P. S., & Hertz-Pannier, L. (2014). The early development of brain white matter: A review of imaging studies in fetuses, newborns and infants. Neuroscience 276, 48–71. 10.1016/j.neuroscience.2013.12.044

Edwards, L. J., Pine, K. J., Ellerbrock, I., Weiskopf, N., & Mohammadi, S. (2017). NODDI-DTI: Estimating neurite orientation and dispersion parameters from a diffusion tensor in healthy white matter. Frontiers in Neuroscience 11, T2&. 10.3389/fnins.2017.00720

Fan, Q., Nummenmaa, A., Witzel, T., Ohringer, N., Tian, Q., Setsompop, K., Klawiter, E. C., Rosen, B. R., Wald, L. L., & Huang, S. Y. (2020). Axon diameter index estimation independent of fiber orientation distribution using high-gradient diffusion MRI. NeuroImage 222, 117197. 10.1016/j.neuroimage.2020.117197

Farbota, K. D., Bendlin, B. B., Alexander, A. L., Rowley, H. A., Dempsey, R. J., & Johnson, S. C. (2012). Longitudinal diffusion tensor imaging and neuropsychological correlates in traumatic brain injury patients. Frontiers in Human Neuroscience 6, 1–15. 10.3389/fnhum.2012.00160

Fieremans, E., Jensen, J. H., & Helpern, J. A. (2011). White matter characterization with diffusional kurtosis imaging. NeuroImage 53(1), 177–188. 10.1016/j.neuroimage.2011.06.006

Friston, K. J., & Ashburner, J. (1997). Multimodal image coregistration and partitioning—a unified framework. NeuroImage 6(3), 209–217. 10.1006/nimg.1997.0290

Garyfallidis, E., Brett, M., Amirbekian, B., Rokem, A., van der Walt, S., Descoteaux, M., & Nimmo-Smith, I. (2014). Dipy, a library for the analysis of diffusion MRI data. Frontiers in Neuroinformatics 3, 8. 10.3389/fninf.2014.00008

Gerstner, E. R., & Sorensen, A. G. (2011). Diffusion and diffusion tensor imaging in brain cancer. Seminars in Radiation Oncology 21(*2*), 141–146. 10.1016/j.semradonc.2010.10.005

Gorgolewski, K. J., Auer, T., Calhoun, V. D., Craddock, R. C., Das, S., Duff, E. P., Flandin, G., Ghosh, S. S., Glatard, T., Halchenko, Y. O., Handwerker, D. A., Hanke, M., Keator, D., Li, X., Michael, Z., Maumet, C., Nichols, B. N., Nichols, T. E., Pellman, J., … Poldrack, R. A. (2016). The brain imaging data structure, a format for organizing and describing outputs of neuroimaging experiments. Scientific Data 3(1), 1–9. 10.1038/sdata.2016.44

Grabher, P., Mohammadi, S., Trachsler, A., Friedl, S., David, G., Sutter, R., Weiskopf, N., Thompson, A. J., Curt, A., & Freund, P. (2016). Voxel-based analysis of grey and white matter degeneration in cervical spondylotic myelopathy. Scientific Reports, 6(1), 1–10. 10.1038/srep24636

Grussu, F., Schneider, T., Tur, C., Yates, R. L., Tachrount, M., lanuş, A., Yiannakas, M. C., Newcombe, J., Zhang, H., Alexander, D. C., DeLuca, G. C., & Gandini Wheeler-Kingshott, C. A. M. (2017). Neurite dispersion: a new marker of multiple sclerosis spinal cord pathology? Annals of Clinical and Translational Neurology 4(9), 663–679. 10.1002/acn3.445

Gu, X., & Eklund, A. (2019). Evaluation of six phase encoding based susceptibility distortion correction methods for diffusion MRI. Frontiers in Neuroinformatics, 13, 76. 10.3389/fninf.2019.00076

Gudbjartsson, H., & Patz, S. (1995). The rician distribution of noisy mri data. Magnetic Resonance in Medicine 34(6), 910–914. 10.1002/mrm.1910340618

Hampel, F. R. (1974). The Influence Curve and Its Role in Robust Estimation. Journal of the American Statistical Association 69(346), 383. 10.2307/2285666

Hansen, B., Shemesh, N., & Jespersen, S. N. (2016). Fast imaging of mean, axial and radial diffusion kurtosis. NeuroImage 142, 381–393. 10.1016/j.neuroimage.2016.08.022

Horsfield, M. A., Lai, M., Webb, S. L., Barker, G. J., Tofts, P. S., Turner, R., Rudge, P., & Miller, D. H. (1996). Apparent diffusion coefficients in benign and secondary progressive multiple sclerosis by nuclear magnetic resonance. Magnetic Resonance in Medicine 36(3), 393–400. 10.1002/mrm.1910360310

Horsfield, M. A., Sala, S., Neema, M., Absinta, M., Bakshi, A., Sormani, M. P., Rocca, M. A., Bakshi, R., & Filippi, M. (2010). Rapid semi-automatic segmentation of the spinal cord from magnetic resonance images: Application in multiple sclerosis. NeuroImage 50(2), 446–455. 10.1016/j.neuroimage.2009.12.121

Howard, A. F., Cottaar, M., Drakesmith, M., Fan, Q., Huang, S. Y., Jones, D. K., Lange, F. J., Mollink, J., Rudrapatna, S. U., Tian, Q., Miller, K. L., & Jbabdi, S. (2022). Estimating axial diffusivity in the NODDI model. NeuroImage 262, 119535. 10.1016/j.neuroimage.2022.119535

Huber, E., David, G., Thompson, A. J., Weiskopf, N., Mohammadi, S., & Freund, P. (2018). Dorsal and ventral horn atrophy is associated with clinical outcome after spinal cord injury. Neurology, 90(17), E1510–E1522. 10.1212/WNL.0000000000005361

Jelescu, I. O., de Skowronski, A., Geffroy, F., Palombo, M., & Novikov, D. S. (2022). Neurite Exchange Imaging ((NEXI): A minimal model of diffusion in gray matter with inter-compartment water exchange. NeuroImage 256. 10.1016/j.neuroimage.2022.119277

Jelescu, I. O., Palombo, M., Bagnato, F., & Schilling, K. G. (2020). Challenges for biophysical modeling of microstructure. Journal of Neuroscience Methods, 344,108861. 10.1016/j.jneumeth.2020.108861

Jensen, J. H., Helpern, J. A., Ramani, A., Lu, H., & Kaczynski, K. (2005). Diffusional kurtosis imaging: The quantification of non-Gaussian water diffusion by means of magnetic resonance imaging. Magnetic Resonance in Medicine 53(*6*), 1432–1440. 10.1002/mrm.20508

Jespersen, S. N., Olesen, J. L., Hansen, B., & Shemesh, N. (2018). Diffusion time dependence of microstructural parameters in fixed spinal cord. NeuroImage 182, 329–342. 10.1016/j.neuroimage.2017.08.039

Jezzard, P., Barnett, A. S., & Pierpaoli, C. (1998). Characterization of and correction for eddy current artifacts in echo planar diffusion imaging. Magnetic Resonance in Medicine 39(*5*), 801–812. 10.1002/mrm.1910390518

Jones, D. K., & Basser, P. J. (2004). ”Squashing peanuts and smashing pumpkins”: How noise distorts diffusion-weighted MR data. Magnetic Resonance in Medicine 52(5), 979–993. 10.1002/mrm.20283

Kellner, E., Dhital, B., Kiselev, V. G., & Reisert, M. (2016). Gibbs-ringing artifact removal based on local subvoxel-shifts. Magnetic Resonance in Medicine 76(5), 1574–1581. 10.1002/mrm.26054

Keim, N. D., West, K. L, Carson, R. P., Gochberg, D. F., Ess, K. C., & Does, M. D. (2016). Evaluation of diffusion kurtosis imaging in ex vivo hypomyelinated mouse brains. NeuroImage 124(0 0), 612­626. 10.1016/j.neuroimage.2015.09.028

Kharbanda, H. S., Alsop, D. C., Anderson, A. W., Filardo, G., & Hackney, D. B. (2006). Effects of cord motion on diffusion imaging of the spinal cord. Magnetic Resonance in Medicine 56(*2*), 334­339. 10.1002/mrm.20959

Koay, C. G., Chang, L. C., Carew, J. D., Pierpaoli, C., & Basser, P. J. (2006). A unifying theoretical and algorithmic framework for least squares methods of estimation in diffusion tensor imaging. Journal of Magnetic Resonance 182(1), 115–125. 10.1016/j.jmr.2006.06.020

Kugler, A. V., & Deppe, M. (2018). Non-lesional cerebellar damage in patients with clinically isolated syndrome: DTI measures predict early conversion into clinically definite multiple sclerosis. NeuroImage: Clinical, 19, 633–639. 10.1016/j.nicl.2018.04.028

Le Bihan, D., Breton, E., Lallemand, D., Aubin, M. L, Vignaud, J., & Laval-Jeantet, M. (1988). Separation of diffusion and perfusion in intravoxel incoherent motion MR imaging. Radiology, 168(*2*), 497–505. 10.1148/radiology.168.2.3393671

Leemans, A., Jeurissen, B., Sijbers, J., & Jones, D. K. (2009). ExploreDTI: A Graphical Toolbox for Processing, Analyzing, and Visualizing Diffusion MR Data. Proceedings of the 17th Scientific Meeting, International Society for Magnetic Resonance in Medicine, Honolulu, USA, 3537. https://archive.ismrm.org/2009/3537.html

Littlejohns, T. J., Holliday, J., Gibson, L. M., Garratt, S., Oesingmann, N., Alfaro-Almagro, F., Bell, J. D., Boultwood, C., Collins, R., Conroy, M. C., Crabtree, N., Doherty, N., Frangi, A. F., Harvey, N. C., Leeson, P., Miller, K. L., Neubauer, S., Petersen, S. E., Sellors, J.,… Allen, N. E. (2020). The UK Biobank imaging enhancement of 100,000 participants: rationale, data collection, management and future directions. Nature Communicatione 11(*1*), 1–12. 10.1038/s41467-020-15948-9

Macdonald, J., & Ruthotto, L. (2018). Improved Susceptibility Artifact Correction of Echo-Planar MRI using the Alternating Direction Method of Multipliers. Journal of Mathematical Imaging and Vision 60(*2*), 268–282. 10.1007/sl0851-017-0757-x

Mangin, J. F., Poupon, C., Clark, C., Le Bihan, D., & Bloch, I. (2002). Distortion correction and robust tensor estimation for MR diffusion imaging. Medical Image Analysie 6(3), 191–198. 10.1016/S1361-8415(02)00079-8

Manjon, J. V., Coupé, P., Concha, L., Buades, A., Collins, D. L., & Robles, M. (2013). Diffusion Weighted Image Denoising Using Overcomplete Local PCA. PLOS ONE 8(9), e73021. 10.1371/JOURNAL.PONE.0073021

Martin, A. R., Aleksanderek, L, Cohen-Adad, J., Tarmohamed, Z., Tetreault, L., Smith, N., Cadotte, D. W., Crawley, A., Ginsberg, H., Mikulis, D. J., & Fehlings, M. G. (2016). Translating state-of-the-art spinal cord MRI techniques to clinical use: A systematic review of clinical studies utilizing DTI, MT, MWF, MRS, and fMRI. NeuroImage: Clinical, 10,192–238. 10.1016/j.nicl.2015.ll.019

Meinzer, M., Mohammadi, S., Kugel, H., Schiffbauer, H., Flöel, A., Albers, J., Kramer, K., Menke, R., Baumgärtner, A., Knecht, S., Breitenstein, C., & Deppe, M. (2010). Integrity of the hippocampus and surrounding white matter is correlated with language training success in aphasia. NeuroImage 53(1), 283–290. 10.1016/j.neuroimage.2010.06.004

Miller, A. J., & Joseph, P. M. (1993). The use of power images to perform quantitative analysis on low SNR MR images. Magnetic Resonance Imaging 11(7), 1051–1056. 10.1016/0730-725X(93)90225-3

Miller, S. P., Vigneron, D. B., Henry, R. G., Bohland, M. A., Ceppi-Cozzio, C., Hoffman, C., Newton, N., Partridge, J. C., Ferriero, D. M., & Barkovich, A. J. (2002). Serial quantitative diffusion tensor MRI of the premature brain: Development in newborns with and without injury. Journal of Magnetic Resonance Imaging 16(*6*), 621–632. 10.1002/jmri.10205

Modersitzki, J. (2009). FAIR - Flexible Algorithms for Image Registration. Society for Industrial and Applied Mathematics (SIAM, 3600 Market Street, Floor 6, Philadelphia, PA 19104). https://epubs.siam.Org/doi/pdf/10.1137/l.9780898718843.bm

Mohammadi, S., & Callaghan, M. F. (2021). Towards in vivo g-ratio mapping using MRI: Unifying myelin and diffusion imaging. Journal of Neuroscience Methods, 348,108990. 10.1016/j.jneumeth.2020.108990

Mohammadi, S., Carey, D., Dick, F., Diedrichsen, J., Sereno, M. I., Reisert, M., Callaghan, M. F., & Weiskopf, N. (2015). Whole-brain in-vivo measurements of the axonal G-ratio in a group of 37 healthy volunteers. Frontiers in Neuroscience 9,1–13. 10.3389/fnins.2015.00441

Mohammadi, S., Freund, P., Feiweier, T., Curt, A., & Weiskopf, N. (2013). The impact of post­processing on spinal cord diffusion tensor imaging. NeuroImage 70, 377–385. 10.1016/j.neuroimage.2012.12.058

Mohammadi, S., Hutton, C., Nagy, Z., Josephs, O., & Weiskopf, N. (2013). Retrospective correction of physiological noise in DTI using an extended tensor model and peripheral measurements. Magnetic Resonance in Medicine 70(*2*), 358–369. 10.1002/mrm.24467

Mohammadi, S., Möller, H. E., Kugel, H., Müller, D. K., & Deppe, M. (2010). Correcting eddy current and motion effects by affine whole-brain registrations: Evaluation of three-dimensional distortions and comparison with slicewise correction. Magnetic Resonance in Medicine 64(*4*), 1047–1056. 10.1002/mrm.22501

Mohammadi, S., Tabelow, K., Ruthotto, L., Feiweier, T., Polzehl, J., & Weiskopf, N. (2015). High-resolution diffusion kurtosis imaging at 3T enabled by advanced post-processing. Frontiers in Neuroscience 9, 427. 10.3389/fnins.2014.00427

Mollink, J., Kleinnijenhuis, M., van Cappellen van Walsum, A.-M., Sotiropoulos, S. N., Cottaar, M., Mirfin, C., Heinrich, M. P., Jenkinson, M., Pallebage-Gamarallage, M., Ansorge, O., Jbabdi, S., & Miller, K. L. (2017). Evaluating fibre orientation dispersion in white matter: Comparison of diffusion MRI, histology and polarized light imaging. NeuroImage 157, 561–574. 10.1016/j.neuroimage.2017.06.001

Novikov, D. S. (2021). The present and the future of microstructure MRI: From a paradigm shift to normal science. Journal of Neuroscience Methode 351. 10.1016/j.jneumeth.2020.108947

Novikov, D. S., Fieremans, E., Jespersen, S. N., & Kiselev, V. G. (2019). Quantifying brain microstructure with diffusion MRI: Theory and parameter estimation. NMR in Biomedicine, 32(4). 10.1002/nbm.3998

Novikov, D. S., Veraart, J., Jelescu, I. O., & Fieremans, E. (2018). Rotationally-invariant mapping of scalar and orientational metrics of neuronal microstructure with diffusion MRI. NeuroImage, 174, 518–538. 10.1016/j.neuroimage.2018.03.006

Oeschger, J. M., Tabelow, K., & Mohammadi, S. (2023a). Axisymmetric diffusion kurtosis imaging with Rician bias correction: A simulation study. Magnetic Resonance in Medicine 89(*2*), 787–799. 10.1002/mrm.29474

Oeschger, J. M., Tabelow, K., & Mohammadi, S. (2023b). Investigating apparent differences between standard DKI and axisymmetric DKI and its consequences for biophysical parameter estimates. BioRxiv, 2023.06.21.545891. 10.1101/2023.06.21.545891

Palombo, M., Ianus, A., Guerreri, M., Nunes, D., Alexander, D. C., Shemesh, N., & Zhang, H. (2020). SANDI: A compartment-based model for non-invasive apparent soma and neurite imaging by diffusion MRI. NeuroImage 215,116835. 10.1016/j.neuroimage.2020.116835

Papazoglou, S., Ashtarayeh, M., Oeschger, J. M., Callaghan, M. F., Does, M. D., & Mohammadi, S. (2023). Insights and improvements in correspondence between axonal volume fraction measured with diffusion-weighted MRI and electron microscopy. NMR in Biomedicine. 10.1002/nbm.5070

Paschoal, A. M., Zotin, M. C. T., da Costa, L. M., dos Santos, A. C., & Leoni, R. F. (2022). Feasibility of intravoxel incoherent motion in the assessment of tumor microvasculature and blood-brain barrier integrity: a case-based evaluation of gliomas. Magnetic Resonance Materials in Physics, Biology and Medicine 35(1), 17–27. 10.1007/sl0334-021-00987-0

Perone, C. S., Calabrese, E., & Cohen-Adad, J. (2018). Spinal cord gray matter segmentation using deep dilated convolutions. Scientific Reports 8(1), 1–13. 10.1038/s41598-018-24304-3

Pierpaoli, C., Walker, L., Irfanoglu, M. O., Barnett, A., Basser, P., Chang, L.-C., Koay, C., Pajevic, S., Rohde, G., Sarlls, J., & Wu, M. (2010). TORTOISE: an integrated software package for processing of diffusion MRI data. Proceedings of the 18th Scientific Meeting, International Society for Magnetic Resonance in Medicine, Stockholm, Sweden, abstract #1597.

Polzehl, J., & Tabelow, K. (2016). Low SNR in diffusion MRI models. Journal of the American Statistical Association, 111(516), 1480–1490. 10.1080/01621459.2016.1222284

Raja, R., Sinha, N., Saini, J., Mahadevan, A., Rao, K. N., & Swaminathan, A. (2016). Assessment of tissue heterogeneity using diffusion tensor and diffusion kurtosis imaging for grading gliomas. Neuroradiology 58(12), 1217–1231. 10.1007/s00234-016-1758-y

Roebroeck, A., Miller, K. L., & Aggarwal, M. (2019). Ex vivo diffusion MRI of the human brain: Technical challenges and recent advances. NMR in Biomedicine 32(*4*), e3941. 10.1002/nbm.3941

Rohde, G. K., Barnett, A. S., Basser, P. J., Marenco, S., & Pierpaoli, C. (2004). Comprehensive approach for correction of motion and distortion in diffusion-weighted MRI. Magnetic Resonance in Medicine 51(1), 103–114. 10.1002/mrm.10677

Rousseeuw, P. J., & Croux, C. (1993). Alternatives to the median absolute deviation. Journal of the American Statistical Association 88(424), 1273–1283. 10.1080/01621459.1993.10476408

Ruthotto, L., Kugel, H., Olesch, J., Fischer, B., Modersitzki, J., Burger, M., & Wolters, C. H. (2012). Diffeomorphic susceptibility artifact correction of diffusion-weighted magnetic resonance images. Physics in Medicine and Biology 57(18), 5715–5731. 10.1088/0031-9155/57/18/5715

Ruthotto, L., Mohammadi, S., Heck, C., Modersitzki, J., & Weiskopf, N. (2013). Hyperelastic susceptibility artifact correction of DTI in SPM. *Proceedings of the German Workshop on Medical Image Computing (Informatik Aktuell), Heidelberg*, Germany, 344–349. 10.1007/978-3-642-36480-8_60

Salvador, R., Pena, A., Menon, D. K., Carpenter, T. A., Pickard, J. D., & Bullmore, E. T. (2005). Formal characterization and extension of the linearized diffusion tensor model. Human Brain Mapping, 24(*2*), 144–155. 10.1002/hbm.20076

Schilling, K. G., Combes, A. J. E., Ramadass, K., Rheault, F., Sweeney, G., Prock, L., Sriram, S., Cohen-Adad, J., Gore, J. C., Landman, B. A., Smith, S. A., & O’Grady, K. P. (2024). Influence of preprocessing, distortion correction and cardiac triggering on the quality of diffusion MR images of spinal cord. Magnetic Resonance Imaging 108, 11–21. 10.1016/j.mri.2024.01.008

Scholz, J., Klein, M. C., Behrens, T. E. J., & Johansen-Berg, H. (2009). Training induces changes in white-matter architecture. Nature Neuroscience 12(11), 1370–1371. 10.1038/nn.2412

Sébille, S. B., Rolland, A. S., Welter, M. L., Bardinet, E., & Santin, M. D. (2019). Post mortem high resolution diffusion MRI for large specimen imaging at 11.7 T with 3D segmented echo-planar imaging. Journal of Neuroscience Methods 311, 222–234. 10.1016/j.jneumeth.2018.10.010

Seif, M., David, G., Huber, E., Vallotton, K., Curt, A., & Freund, P. (2020). Cervical cord neurodegeneration in traumatic and non-traumatic spinal cord injury. Journal of Neurotrauma, 37(*6*), 860–867. 10.1089/neu.2019.6694

Sijbers, J., den Dekker, A. J., Scheunders, P., & Van Dyck, D. (1998). Maximum-likelihood estimation of rician distribution parameters. IEEE Transactions on Medical Imaging 17(3), 357–361. 10.1109/42.712125

Smith, S. M., Jenkinson, M., Woolrich, M. W., Beckmann, C. F., Behrens, T. E. J., Johansen-Berg, H., Bannister, P. R., De Luca, M., Drobnjak, I., Flitney, D. E., Niazy, R. K., Saunders, J., Vickers, J., Zhang, Y., De Stefano, N., Brady, J. M., & Matthews, P. M. (2004). Advances in functional and structural MR image analysis and implementation as FSL. NeuroImage 23(Suppl. 1), S208–S219. 10.1016/j.neuroimage.2004.07.051

Sotiropoulos, S. N., Moeller, S., Jbabdi, S., Xu, J., Andersson, J. L., Auerbach, E. J., Yacoub, E., Feinberg, D., Setsompop, K., Wald, L. L., Behrens, T. E. J., Ugurbil, K., & Lenglet, C. (2013). Effects of image reconstruction on fiber orientation mapping from multichannel diffusion MRI: Reducing the noise floor using SENSE. Magnetic Resonance in Medicine 70(*6*), 1682–1689. 10.1002/mrm.24623

Stejskal, E. O., & Tanner, J. E. (1965). Spin diffusion measurements: Spin echoes in the presence of a time-dependent field gradient. The Journal of Chemical Physice 42(1), 288–292. 10.1063/l.1695690

Stikov, N., Campbell, J. S. W., Stroh, T., Lavelée, M., Frey, S., Novek, J., Nuara, S., Ho, M. K., Bedell, B. J., Dougherty, R. F., Leppert, I. R., Boudreau, M., Narayanan, S., Duval, T., Cohen-Adad, J., Picard, P. A., Gasecka, A., Côté, D., & Pike, G. B. (2015). In vivo histology of the myelin g-ratio with magnetic resonance imaging. NeuroImage 118, 397–405. 10.1016/j.neuroimage.2015.05.023

Stroman, P. W., Wheeler-Kingshott, C., Bacon, M., Schwab, J. M., Bosma, R., Brooks, J., Cadotte, D., Carlstedt, T., Ciccarelli, O., Cohen-Adad, J., Curt, A., Evangelou, N., Fehlings, M. G., Filippi, M., Kelley, B. J., Koi lias, S., Mackay, A., Porro, C. A., Smith, S.,… Tracey, I. (2014). The current state-of-the-art of spinal cord imaging: Methods. NeuroImage 84, 1070–1081. 10.1016/j.neuroimage.2013.04.124

Sullivan, E. V., Rohlfing, T., & Pfefferbaum, A. (2010). Quantitative fiber tracking of lateral and interhemispheric white matter systems in normal aging: Relations to timed performance. Neurobiology of Aging 31(3), 464–481. 10.1016/j.neurobiolaging.2008.04.007

Summers, P., Staempfli, P., Jaermann, T., Kwiecinski, S., & Kollias, S. S. (2006). A preliminary study of the effects of trigger timing on diffusion tensor imaging of the human spinal cord. American Journal of Neuroradiology 27(*9*), 1952–1961.

Szturm, T., Kolesar, T. A., Mahana, B., Goertzen, A. L., Hobson, D. E., Marotta, J. J., Strafella, A. P., & Ko, J. H. (2021). Changes in metabolic activity and gait function by dual-task cognitive game-based treadmill system in Parkinson’s disease: Protocol of a randomized controlled trial. Frontiers in Aging Neuroscience 13, 283. 10.3389/fnagi.2021.680270

Tabelow, K., Balteau, E., Ashburner, J., Callaghan, M. F., Draganski, B., Helms, G., Kherif, F., Leutritz, T., Lutti, A., Phillips, C., Reimer, E., Ruthotto, L., Seif, M., Weiskopf, N., Ziegler, G., & Mohammadi, S. (2019). hMRI - A toolbox for quantitative MRI in neuroscience and clinical research. NeuroImage 194,191–210. 10.1016/j.neuroimage.2019.01.029

Tabelow, K., Mohammadi, S., Weiskopf, N., & Polzehl, J. (2015). POAS4SPM: a toolbox for SPM to denoise diffusion MRI data. Neuroinformatice 13(1), 19–29. 10.1007/sl2021-014-9228-3

Tabesh, A., Jensen, J. H., Ardekani, B. A., & Helpern, J. A. (2011). Estimation of tensors and tensor-derived measures in diffusional kurtosis imaging. Magnetic Resonance in Medicine 65(*3*), 823­836. 10.1002/mrm.22655

Taylor, P. A., & Saad, Z. S. (2013). FATCAT: (An Efficient) functional and tractographic connectivity analysis toolbox. Brain Connectivity 3(5), 523–535. 10.1089/brain.2013.0154

Tournier, J. D., Smith, R., Raffelt, D., Tabbara, R., Dhollander, T., Pietsch, M., Christiaens, D., Jeurissen, B., Yeh, C. H., & Connelly, A. (2019). MRtrix3: A fast, flexible and open software framework for medical image processing and visualisation. NeuroImage 202, 116137. 10.1016/j.neuroimage.2019.116137

Urbach, H., Flacke, S., Keller, E., Textor, J., Berlis, A., Hartmann, A., Reul, J., Solymosi, L., & Schild, H. H. (2000). Detectability and detection rate of acute cerebral hemisphere infarcts on CT and diffusion-weighted MRI. Neuroradiology 42(10), 722–727. 10.1007/s002340000401

Vallotton, K., David, G., Hupp, M., Pfender, N., Cohen-Adad, J., Fehlings, M. G., Samson, R. S., Gandini Wheeler-Kingshott, C. A. M., Curt, A., Freund, P., & Seif, M. (2021). Tracking white and gray matter degeneration along the spinal cord axis in degenerative cervical myelopathy. Journal of Neurotrauma 38(21), 2978–2987. 10.1089/neu.2021.0148

Van Essen, D. C., Smith, S. M., Barch, D. M., Behrens, T. E. J., Yacoub, E., & Ugurbil, K. (2013). The WU-Minn Human Connectome Project: An overview. NeuroImage 80, 62–79. 10.1016/j.neuroimage.2013.05.041

Van Essen, D. C., Ugurbil, K., Auerbach, E., Barch, D., Behrens, T. E. J., Bucholz, R., Chang, A., Chen, L, Corbetta, M., Curtiss, S. W., Della Penna, S., Feinberg, D., Glasser, M. F., Harel, N., Heath, A. C., Larson-Prior, L., Marcus, D., Michalareas, G., Moeller, S.,… Yacoub, E. (2012). The Human Connectome Project: A data acquisition perspective. NeuroImage 62, 2222–2231. 10.1016/j.neuroimage.2012.02.018

Veraart, J., Novikov, D. S., Christiaens, D., Ades-aron, B., Sijbers, J., & Fieremans, E. (2016). Denoising of diffusion MRI using random matrix theory. NeuroImage 142, 394–406. 10.1016/j.neuroimage.2016.08.016

Veraart, J., Rajan, J., Peeters, R. R., Leemans, A., Sunaert, S., & Sijbers, J. (2013). Comprehensive framework for accurate diffusion MRI parameter estimation. Magnetic Resonance in Medicine, 70(*4*), 972–984. 10.1002/mrm.24529

Veraart, J., Sijbers, J., Sunaert, S., Leemans, A., & Jeurissen, B. (2013). Weighted linear least squares estimation of diffusion MRI parameters: Strengths, limitations, and pitfalls. NeuroImage 81, 335–346. 10.1016/j.neuroimage.2013.05.028

Veraart, J., Van Hecke, W., & Sijbers, J. (2011). Constrained maximum likelihood estimation of the diffusion kurtosis tensor using a Rician noise model. Magnetic Resonance in Medicine 66(3), 678–686. 10.1002/mrm.22835

Weiskopf, N., Edwards, L. J., Helms, G., Mohammadi, S., & Kirilina, E. (2021). Quantitative magnetic resonance imaging of brain anatomy and in vivo histology. Nature Reviews Physics, 3, 570–588. 10.1038/s42254-021-00326-l

Woletz, M., Chalupa-Gantner, F., Hager, B., Rieke, A., Mohammadi, S., Binder, S., Baudis, S., Ovsianikov, A., Windischberger, C., & Nagy, Z. (2024). Toward Printing the Brain: A Microstructural Ground Truth Phantom for MRI. Advanced Materials Technologies, 2300176. 10.1002/admt.202300176

Yiannakas, M. C., Kearney, H., Samson, R. S., Chard, D. T., Ciccarelli, O., Miller, D. H., & Wheeler-Kingshott, C. A. M. (2012). Feasibility of grey matter and white matter segmentation of the upper cervical cord in vivo: A pilot study with application to magnetisation transfer measurements. NeuroImage 63(*3*), 1054–1059. 10.1016/j.neuroimage.2012.07.048

Zhang, H., Schneider, T., Wheeler-Kingshott, C. A., & Alexander, D. C. (2012). NODDI: Practical in vivo neurite orientation dispersion and density imaging of the human brain. NeuroImage 61(*4*), 1000–1016. 10.1016/j.neuroimage.2012.03.072

Zwiers, M. P. (2010). Patching cardiac and head motion artefacts in diffusion-weighted images. NeuroImage 53(*2*), 565–575. 10.1016/j.neuroimage.2010.06.014

